# Non-cell-autonomous removal of protein aggregates via trogocytosis

**DOI:** 10.64898/2026.10.13.758691

**Authors:** Yoshitaka Kimura, Kazuki Sato, Arun R. Chavan, William Kasberg, Malan Silva, Wei Qiu, Nirmala Iyer, Michael Innerberger, Igor Siwanowicz, Scott D. Pope, Athreya Rangavajhula, Gang-Qing Yao, Karl Insogna, Eric T. Trautman, Christopher K. E. Bleck, Xiaolei Su, Jennifer Lippincott-Schwartz, Ruslan Medzhitov

**Affiliations:** Department of Immunobiology, Yale University School of Medicine, New Haven, CT, USA; Department of Cell Biology, Yale University School of Medicine, New Haven, CT, USA; Janelia Research Campus, Howard Hughes Medical Institute, Ashburn, VA, USA; Enhanced FIB-SEM Imaging Shared Resource, Janelia Research Campus, Howard Hughes Medical Institute, Ashburn, VA, USA; Electron Microscopy Shared Resource, Janelia Research Campus, Howard Hughes Medical Institute, Ashburn, VA, USA; Project Technical Resources, Janelia Research Campus, Howard Hughes Medical Institute, Ashburn, VA, USA; Department of Internal Medicine, Yale University School of Medicine, New Haven, CT, USA; Yale Cancer Center, Yale University, New Haven, CT, USA; Howard Hughes Medical Institute, Yale University School of Medicine, New Haven, CT, USA; Tananbaum Center for Theoretical and Analytical Human Biology, Yale University School of Medicine, New Haven, CT, USA

## Abstract

Protein aggregate formation can cause cell damage and contribute to tissue pathologies including neurodegeneration. Protein aggregates are normally eliminated by cell-intrinsic mechanisms of lysosomal (autophagic) degradation. However, degradative capacity varies between cell types, raising the possibility that high-capacity cell types, such as macrophages, may contribute to aggregate disposal from the low-capacity cells. Here, we show that cellular protein aggregates are removed by macrophages via trogocytosis — a partial engulfment of the target cells. Lysosomal stress upon protein aggregate accumulation promotes cell surface expression of the transmembrane isoform of the macrophage growth factor CSF1 (tmCSF1), which is detected by macrophages through the CSF1 receptor to drive trogocytosis. Insufficient or impaired degradation of protein aggregates in lysosomes results in their transport along microtubules to the plasma membrane where they physically associate with tmCSF1. This enables macrophages to specifically trogocytose protein aggregates. These findings reveal a protein homeostasis system that eliminates protein aggregates in a non-cell-autonomous manner.

## Main text

Protein aggregates are byproducts of protein translation. They form upon the accumulation of misfolded proteins produced by proteostatic stress or errors in protein folding. Protein aggregates cause cellular damage by multiple mechanisms. They interfere with lysosomal and mitochondrial function, disrupt plasma membrane integrity, and induce reactive oxygen species^1^. The protein aggregate-induced decline in cellular fitness leads to tissue pathologies such as neurodegenerative diseases and cardiomyopathies^2,3^. The accumulation of protein aggregates also occurs with aging, contributing to age-associated tissue dysfunction^4^.

All organisms have diverse evolutionarily conserved mechanisms to prevent protein aggregation and remove aggregated proteins. Misfolded proteins are recognized by chaperones to correct folding or are ubiquitinated for proteasome-mediated degradation^5,6^. Proteins exposing the KFERQ amino acid motif upon a folding error are transported to lysosomes for degradation by the chaperon-mediated autophagy^7^. When these systems fail to handle misfolded proteins, autophagosomes incorporate aggregated proteins and fuse with lysosomes to degrade them^8^. Alternatively, vesicle-enclosed aggregates can be released from cells^9^.

While cell-autonomous defense is a general response to stress, cellular homeostasis is also maintained through cell-to-cell communication in multicellular organisms. For example, upon bacterial infection, cells produce antimicrobial proteins as an autonomous defense but also release chemokines to recruit immune cells that remove invading pathogens^10,11^. While cells under hypoxic conditions shift energy production from oxidative phosphorylation to glycolysis, they communicate with endothelial cells via VEGF to initiate angiogenesis and restore oxygen supply^12,13^. These examples illustrate the critical roles of cell communication and division of labor in tissue-level stress responses. Here we explored whether cells undergoing stress can recruit help from ‘professional’ cell types when cell-autonomous stress responses are insufficient.

### Cellular protein aggregates are removed by macrophages via trogocytosis

To model insufficient proteostasis, mouse embryonic fibroblasts (MEFs) were treated with bafilomycin A1 (Baf A1), which suppresses lysosomal acidification and degradation of misfolded proteins, thereby leading to protein aggregate accumulation. Previous seminal studies have shown that macrophages monitor the state of neighboring cells and support their fitness in tissue^14^. We hypothesized that macrophages play a critical role in assisting the maintenance of homeostasis in protein aggregate-accumulating cells. To test this, Baf A1-treated MEFs (MEF^Baf^ ^A1^) were co-cultured with bone marrow-derived macrophages (BMDMs) (Fig. 1a). We used CellTracker Deep Red (CTDR), a reagent that covalently binds to amino acids, to visualize undigested proteins as punctate material inside the cell. Time-lapse confocal imaging revealed that punctate cell materials labelled by CTDR in MEF^Baf^ ^A^^1^ were captured and removed by macrophages (Fig. 1b, c and Supplementary Videos 1 and 2). The incorporated materials were beneath the wheat germ agglutinin (WGA)-labelled plasma membrane and were transported to lysosomes in macrophages (Fig. 1d, e), indicating that MEF^Baf^ ^A^^1^-derived materials were internalized and processed for degradation. To identify the macrophage-engulfed material, correlative light and electron microscopy (CLEM) was performed on MEF^Baf^ ^A1^-BMDM co-culture. We found that MEF^Baf^ ^A1^-derived material in macrophages was heterogeneous but mainly composed of electron-dense clusters, a typical appearance of protein aggregates under electron microscopy (Fig. 1f). Furthermore, a fluorescent chemical proteostat, which binds to cross-β spine structure of protein aggregates, stained more than 90% of MEF^Baf^ ^A1^-derived materials in BMDMs (Fig. 1g, h). CTDR-puncta in MEF^Baf^ ^A1^ were also proteostat-positive (Fig. 1g, h). Consistently, proteostat-positive puncta were co-stained with another dye with high affinity for protein aggregates, thioflavin T, in MEF^Baf^ ^A1^ (Extended Data Fig. 1a). Proteostat-positive materials were detergent-resistant and detected with flow cytometry as particles (Supplementary Fig. 1a). These results demonstrate that the undigested materials that macrophages incorporate from MEF^Baf^ ^A1^ were protein aggregates. These observations were not due to an experimental artifact caused by CTDR labelling, given that protein aggregates were observed in BMDMs co-cultured with CTDR-free MEF^Baf^ ^A1^ (Extended Data Fig. 1b). When MEF^Baf^ ^A1^ and BMDMs were cultured in a transwell system, negligible amount of protein aggregate transfer was detected (Fig. 1i, j), indicating the requirement of cell-to-cell contact. These data suggest that macrophage uptake of protein aggregates occurs via trogocytosis, a process in which cells nibble partial bodies of the target cells^15,16^. Indeed, Focused Ion Beam-Scanning Electron Microscopy (FIB-SEM) revealed an extended portion of MEF^Baf^ ^A1^ cell body containing electron-dense conglomerates in a phagocytic cup-like invagination of BMDMs (Fig. 1k-m and Supplementary Video 3). Thus, cellular protein aggregates are removed by macrophages through trogocytosis.

**Fig. 1.**
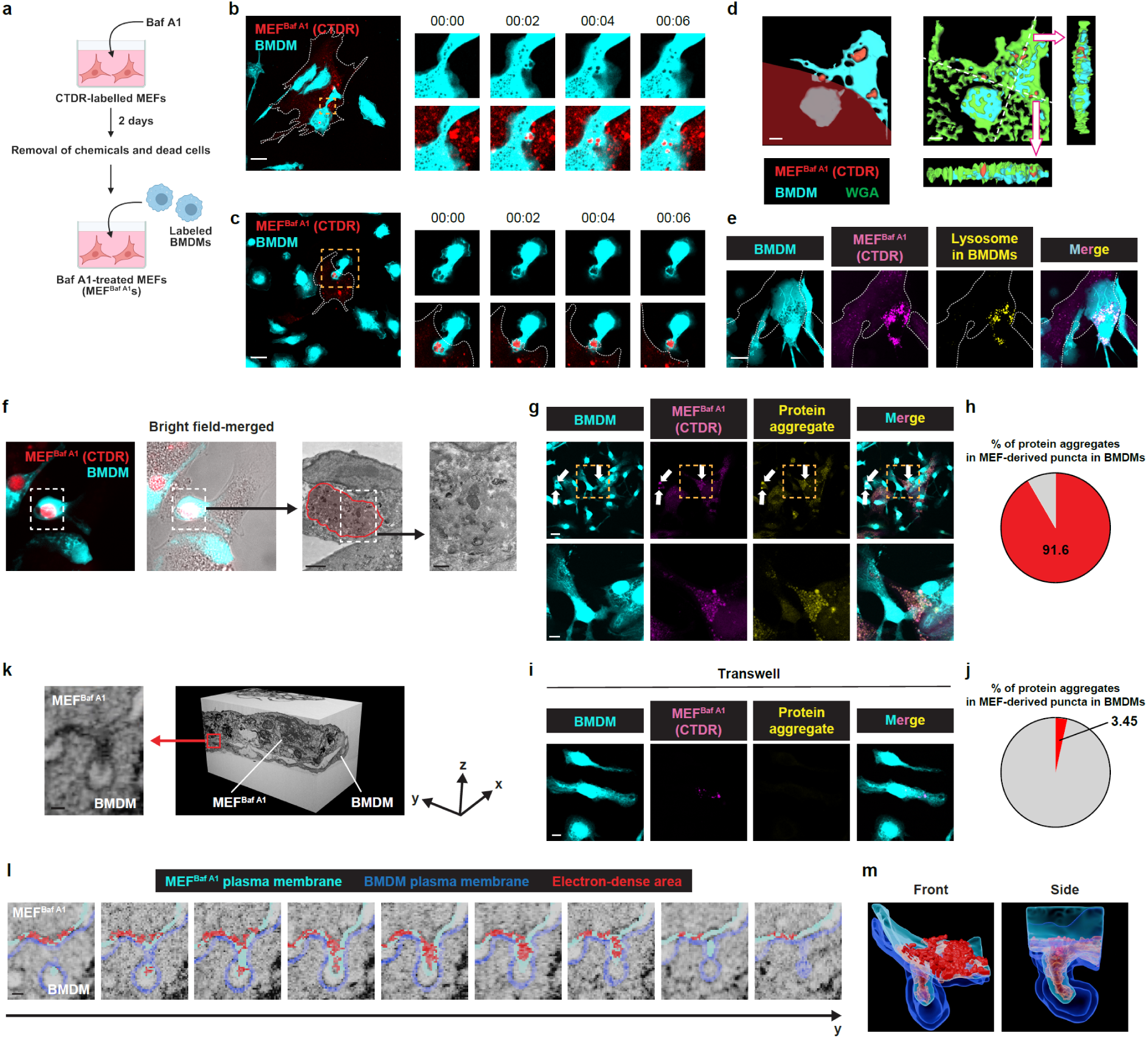
Cellular protein aggregates are engulfed by macrophages via trogocytosis. a, Schematic of the experimental process for co-culture of BMDMs and Baf A1-treated MEFs (MEF^Baf^ ^A1^). CTDR = CellTracker Deep Red. **b, c,** Representative time-lapse confocal images of co-cultured BMDMs and MEF^Baf^ ^A1^. The left panel shows the image at 00:00. The upper-right and lower-right panels show the zoomed images of the white-dotted areas in the left panel. Images taken every 2 min are shown. **d**, Representative 3D-rendered images of co-cultured BMDMs and MEF^Baf^ ^A1^. The red-shaded area in the left panel indicates the cell body of MEF^Baf^ ^A1^. Striped panels show the images sectioned along the white-dotted lines in the right panel. **e,** Representative confocal images of co-cultured BMDMs with pre-labelled lysosomes and MEF^Baf^ ^A1^. **f,** Representative CLEM images of co-cultured BMDMs and MEF^Baf^ ^A1^. The two left panels show fluorescent microscopy images. The second panel from the right displays electron microscopy image of the areas indicated by the white-dotted boxes in the two left panels. The red-shaded area indicates an electron-dense region surrounded by membranes. The far-right panel shows a zoomed image of the area indicated by the white-dotted box in the second panel from the right. **g,** Representative confocal images of co-cultured BMDMs and MEF^Baf^ ^A1^. White arrows indicate sites where MEF-derived dye and protein aggregates are co-localized in BMDMs. The lower panels display zoomed images of the areas indicated by orange-dotted boxes in the upper panels. **h,** Percentage of protein aggregates co-localized with MEF-derived dye puncta (n = 119) in BMDMs co-cultured with MEF^Baf^ ^A1^. **i,** Representative confocal images of BMDMs upon contactless co-culture with MEF^Baf^ ^A1^ in a transwell system. **j,** Percentage of protein aggregates co-localized with MEF-derived dye puncta (n = 58) in BMDMs co-cultured with MEF^Baf^ ^A1^ in a contactless manner with a transwell system. **k-m,** FIB-SEM analysis of co-cultured BMDMs and MEF^Baf^ ^A1^. **k,** The right panel shows the raw 3D image. The left panel displays a representative x-z plane image indicating macrophage uptake of a partial MEF^Baf^ ^A1^ body. **l,** x-z plane images along y-axis indicating macrophage uptake of a partial MEF^Baf^ ^A1^ body. The electron-dense area and plasma membranes of MEF^Baf^ ^A1^ and BMDM are segmented with the indicated colors. **m,** 3D-reconstructed images of the segmentation in (**l**). Scale bars, 20 μm (**b, c**), 5 μm (**d,** lower panel of **g, i**), 10 μm (**e,** upper panel of **g**), 2 μm (**f,** the second from the right), 600 nm (**f,** the far right), 50 nm (**k, l**). **a,** Created with BioRender.com

### Transmembrane isoform of CSF1 induces trogocytosis of protein aggregates

We next investigated the mechanism of trogocytic uptake of protein aggregates. Flow cytometric analysis showed enhanced membrane transfer from MEFs to BMDMs following Baf A1 pre-treatment, in a contact-dependent manner (Extended Data Fig. 1c, d and Supplementary Fig. 1b). Using this system, we screened macrophage receptors responsible for the detection and trogocytosis of protein aggregate-harboring cells. Previous studies indicated that some phagocytic receptors have the potential to perform trogocytosis^17–21^. Phagocytic receptors in macrophages belong to three classes: ITAM-dependent receptors, integrins, and TAM (MerTK, Axl, Tyro-3) family receptor tyrosine kinases (RTKs)^22,23^. While FcγR and TREM2 induce ITAM-mediated activation of Syk and can serve as both phagocytic and trogocytic receptors^17,18^, treatment with a Syk inhibitor R406 and TREM2 deficiency in BMDMs did not significantly change the uptake of MEF^Baf^ ^A1^ materials (Extended Data Fig. 1e, f). CD11b is a component of phagocytic integrins and triggers trogocytosis in some contexts^17,19–21^. However, CD11b-deficient BMDMs exhibited uptake activity similar to that of wild-type (WT) cells (Extended Data Fig. 1g). In contrast, inhibition of PI3K, a signaling molecule activated by RTK, with wortmannin reduced uptake (Extended Data Fig. 1h). Treatment with IPI145 and tenalisib, which inhibit the activity of PI3K-δ/γ selectively expressed in immune cells, yielded similar results (Extended Data Fig. 1h). Phagocytic RTKs such as MerTK, Axl and Tyro-3 recognize phosphatidylserine (PS) on target cells^24^. We tested their involvement by masking PS with annexin V. However, no significant decrease in trogocytosis was observed, unlike phagocytosis of heat-killed MEFs (Extended Data Fig. 1i). Together, these data suggested the involvement of unidentified trogocytic RTKs. RNA-sequencing (RNA-seq) analysis revealed that CSF1R was the most highly expressed RTK in BMDMs (Extended Data Fig. 1j). To determine the contribution of CSF1R, BMDMs were co-cultured with MEF^Baf^ ^A1^ in the presence of a blocking antibody or CSF1R kinase inhibitor BLZ-925. We found that both treatments suppressed the uptake activity of BMDMs (Extended Data Fig. 1k, l). These results indicate that CSF1R and its kinase activity are essential for trogocytosis by macrophages.

CSF1, the ligand of CSF1R, is a macrophage growth factor expressed by a wide range of cell types including epithelial cells, endothelial cells, and fibroblasts^25^. While the major isoform is released as a soluble molecule (sCSF1), a minor alternatively spliced isoform resides on the plasma membrane as a transmembrane protein (tmCSF1)^26^. In light of the contact-dependency of the trogocytosis process, we hypothesized that tmCSF1 on protein aggregate-harboring cells is recognized by macrophages and triggers aggregate removal. To determine whether tmCSF1 induces trogocytosis, we established CSF1-deficient cells, transduced them with tmCSF1 (Extended Data Fig. 2a) and co-cultured with mock-transduced cells and BMDMs. Time-lapse imaging showed that materials from tmCSF1-transduced cells were incorporated by macrophages (Fig. 2a and Supplementary Video 4). Uptake from mock-transduced cells was not detected (Fig. 2a and Supplementary Video 4), indicating that tmCSF1 promotes macrophage uptake of cellular materials with negligible effects on bystander cells. In contrast to tmCSF1, sCSF1 did not enhance macrophage uptake of cellular materials upon transduction (Fig. 2b, c and Extended Data Fig. 2b), illuminating the tmCSF1 isoform-specific engagement in cellular material transfer. Although tmCSF1 was cleaved and released by ADAM17 (Extended Data Fig. 2c-e) as described previously^27^, an ADAM17 inhibitor batimastat did not reduce macrophage uptake of materials from tmCSF1-transduced cells (Extended Data Fig. 2f). Accordingly, material transfer from tmCSF1-transduced cells was not observed upon contactless co-culture with BMDMs (Fig. 2d), demonstrating that uncleaved tmCSF1 is essential.

**Fig. 2.**
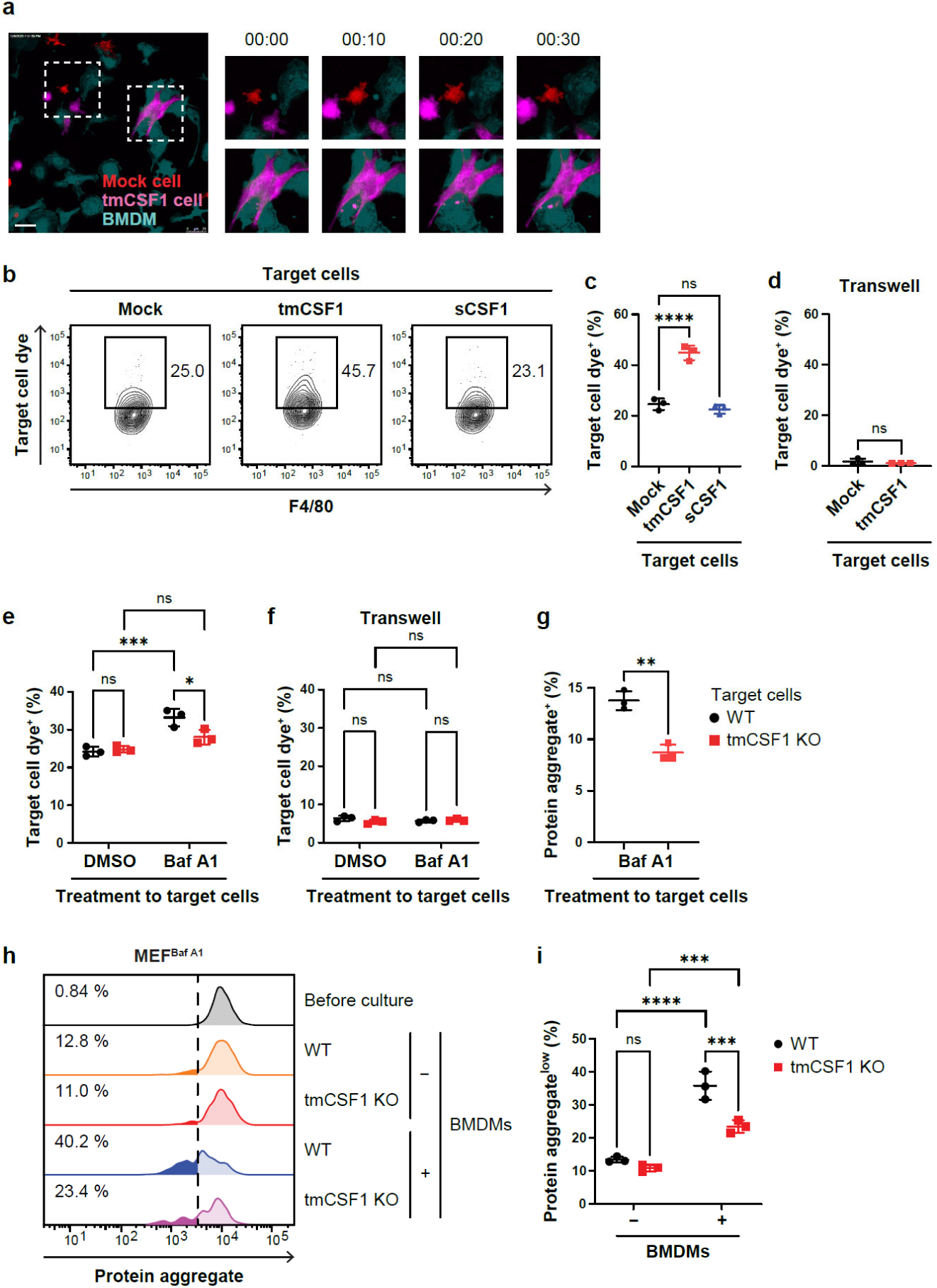
Transmembrane isoform of CSF1 (tmCSF1) on stressed cells induces trogocytosis-mediated removal of protein aggregates by macrophages. a, Representative time-lapse confocal images of co-cultured mock-transduced cells, tmCSF1-transduced cells and BMDMs. The left panel shows the image at 00:00. The upper-right and lower-right panels show the areas indicated by the white-dotted squares in the left panel. Each cell type was stained with a different cytoplasmic dye. Images taken every 10 min are shown. Scale bar, 25 μm. **b, c,** Uptake of materials from mock-, tmCSF1-and sCSF1-transduced cells by BMDMs upon the co-culture with each transduced cell line. Shown are the representative contour plots (**b**) and percentages (**c**) of BMDMs positive for transduced cell-staining cytoplasmic dye. **d,** Uptake of materials from mock-and tmCSF1-transduced cells by BMDMs upon contactless co-culture with each transduced cell line in a transwell system. Shown are the percentages of BMDMs positive for transduced cell-staining cytoplasmic dye. **e, f,** Uptake of materials from WT and tmCSF1 KO MEF^Baf^ ^A1^ by BMDMs upon the co-culture (**e**) or contactless co-culture in a transwell system (**f**). Shown are the percentages of BMDMs positive for MEF-staining membrane dye. **g,** Percentages of BMDMs positive for protein aggregate-staining dye upon the co-culture with WT and tmCSF1 KO MEF^Baf^ ^A1^. **h, i,** Amount of protein aggregates in WT and tmCSF1 KO MEF^Baf^ ^A1^ after culture alone or co-culture with BMDMs. Shown are the representative histograms (**h**) and percentages (**i**) of MEFs with lower intensity of protein aggregate-staining dye than that indicated by the black-dotted line in (**h**). Graphs show mean ± s.d. **c-g, i,** n = 3 per group. **c,** One-way ANOVA with Dunnett’s multiple-comparison test. **d, g,** Two-tailed unpaired Student’s t test. **e, f, i,** Two-way ANOVA with Sidak’s multiple-comparison test. \**P* < 0.05, \*\**P* < 0.01, \*\*\**P* < 0.001, \*\*\*\**P* < 0.0001; ns, not significant.

Macrophage incorporation of cellular materials is likely associated with CSF1R-mediated internalization of tmCSF1 into macrophages, rather than nibbling-independent uptake such as gap junction-mediated material transfer, because tmCSF1 and CSF1R levels on the surface of tmCSF1-transduced cells and BMDMs, respectively, were markedly decreased upon co-culture (Extended Data Fig. 2g-j). Unlike lymphocyte trogocytosis^28–30^, incorporated tmCSF1 was not translocated to the cell surface of macrophages (Extended Data Fig. 2k, l), implicating internalization as the primary fate of this uptake process. Collectively, these data show that tmCSF1 expression on the plasma membrane induces trogocytosis through CSF1R on macrophages.

To assess the requirement of tmCSF1 for trogocytic removal of protein aggregates, MEFs from tmCSF1-knockout (KO) mice^31^ were treated with Baf A1 and co-cultured with BMDMs. The enhancement in macrophage uptake of MEF materials with Baf A1 treatment was abrogated by tmCSF1 deficiency in a cell contact-dependent manner (Fig. 2e, f). In parallel, the amount of protein aggregates detected in BMDMs after co-culture with MEF^Baf^ ^A1^ was significantly lower in the absence of tmCSF1 (Fig. 2g). Consistently, protein aggregate load in WT MEF^Baf^ ^A1^ was significantly decreased by co-culture with BMDMs, whereas the effect was milder in tmCSF1 KO MEF^Baf^ ^A1^ (Fig. 2h, i). Survival of WT MEF^Baf^ ^A1^ was improved by co-culture with BMDMs but not in tmCSF1 KO MEF^Baf^ ^A1^ (Extended Data Fig. 2m). Taken together, tmCSF1 induces macrophage trogocytosis and the associated removal of protein aggregates, thereby contributing to homeostasis maintenance in aggregate-accumulating cells.

Given that sCSF1 binds to CSF1R and induces its internalization into cells via endocytosis^32^, we examined whether sCSF1 suppresses tmCSF1-mediated trogocytosis. CSF1R on BMDMs was mildly but significantly decreased following treatment with culture supernatant from sCSF1-transduced cells (Extended Data Fig. 3a, b). Treatment with recombinant CSF1 at a supraphysiological dose led to a dramatic decrease of cell-surface CSF1R, while residual CSF1R expression was still observed (Extended Data Fig. 3c, d). Nevertheless, tmCSF1-mediated macrophage uptake of co-cultured cell materials was not reduced by culture supernatant from sCSF1-transduced cells and was mitigated but not completely abrogated by supraphysiological dose of recombinant CSF1 (Extended Data Fig. 3e). These results indicate that trogocytosis is not limited by the level of cell-surface CSF1R expression, which may underlie the feasibility of this protein homeostasis system in tissue where sCSF1 is abundant to maintain the macrophage population.

### Lysosomal stress promotes tmCSF1 isoform expression

To determine the mechanisms that control tmCSF1 expression, we designed quantitative polymerase chain reaction (qPCR) primers for *tmCsf1* and *sCsf1* that specifically detected each target isoform in tmCSF1-and sCSF1-transduced cells (Extended Data Fig. 4a). Using these primers, we found that a lysosomal damage inducer L-leucyl-L-leucine methyl ester (LLOMe) upregulated both *tmCsf1* and *sCsf1*, with a greater increase in *tmCsf1*, thereby shifting the *Csf1* isoform balance toward *tmCsf1* (Fig. 3a). We next investigated stress response pathways that govern CSF1 isoform selection. MiT/TFE family members (TFEB, MITF, TFE3 and TFEC) are transcription factors activated in response to lysosomal damage^33^. Knockdown (KD) of MITF, TFE3 and TFEC, but not TFEB, reduced *tmCsf1-*skewing in LLOMe-treated cells (Fig. 3b and Extended Data Fig. 4b). Baf A1 treatment upregulated lysosomal stress-induced genes, particularly at later time points, 48 hours after challenge (Fig. 3c), suggesting that lysosomal damage does not occur immediately after the decline of lysosomal acidification but in association with protein aggregate accumulation. In correlation with lysosomal stress, *tmCsf1-*biased expression was induced by Baf A1 treatment (Fig. 3d). However, this skewing did not result from *tmCsf1* upregulation but rather from *sCsf1* downregulation (Extended Data Fig. 4c). Similar to the result with LLOMe treatment, the *tmCsf1-*biased expression was mediated by the MiT/TFE family members except for TFEB (Fig. 3e and Extended Data Fig. 4d), suggesting that *tmCsf1-*skewing occurs alongside the downregulation of total *Csf1* RNA. Despite the absence of *tmCsf1* upregulation, its protein expression was elevated, and sCSF1 protein release was only modestly changed by Baf A1 treatment (Fig. 3f, g). These results implicate enhanced trafficking of tmCSF1 proteins to cell surface and/or elevation of CSF1 translation upon Baf A1 treatment. Collectively, lysosomal stress responses play crucial roles in balancing CSF1 isoform expression toward tmCSF1.

**Fig. 3.**
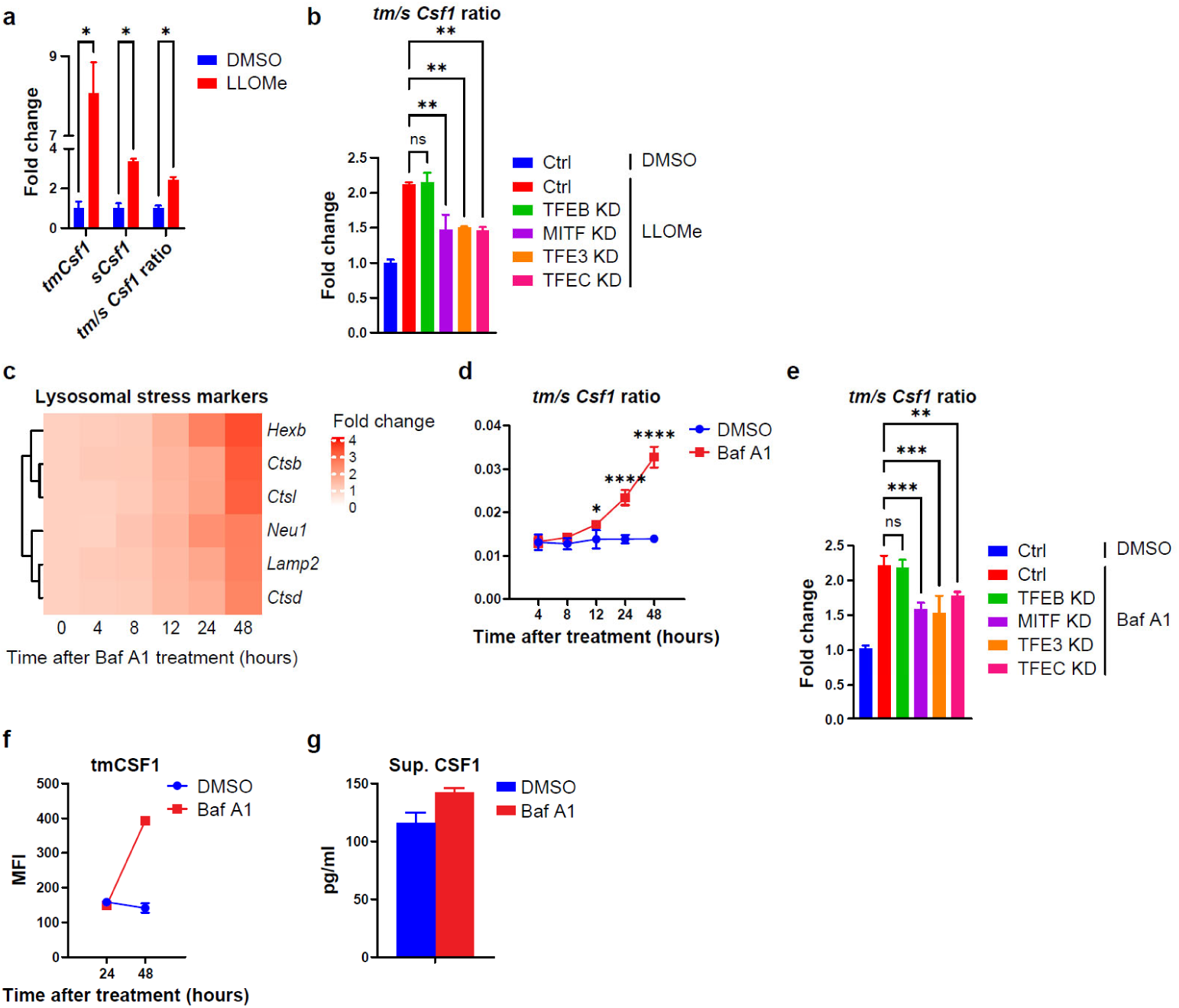
Lysosomal stress induces the skewing of CSF1 isoform balance toward tmCSF1. a, Expression of the indicated genes in MEFs treated with or without L-Leucyl-L-Leucine methyl ester (LLOMe). **b,** *tmCsf1* to *sCsf1* expression ratio in control (Ctrl), TFEB knockdown (KD), MITF KD TFE3 KD or TFEC KD MEFs treated with or without LLOMe. **c,** Expression of lysosomal stress marker genes in MEFs treated with or without Baf A1, analyzed by RNA-seq (n=2 per group). **d,** *tmCsf1* to *sCsf1* expression ratio in MEFs treated with or without Baf A1. **e,** *tmCsf1* to *sCsf1* expression ratio in Ctrl, TFEB KD, MITF KD TFE3 KD or TFEC KD MEFs treated with or without Baf A1. **f, g,** Surface tmCSF1 protein expression (**f**) and CSF1 protein level in the culture supernatant (**g**) of MEFs treated with or without Baf A1. MFI, mean fluorescence intensity. n = 3 per group unless otherwise indicated. Graphs show mean ± SD. **a,** Multiple unpaired Student’s t test. **b, e,** One-way ANOVA with Dunnett’s multiple-comparison test. **d,** Two-way ANOVA with Sidak’s multiple-comparison test. \**P* < 0.05, \*\**P* < 0.01, \*\*\**P* < 0.001, \*\*\*\**P* < 0.0001; ns, not significant.

Based on splicing-mediated control of CSF1 isoform balance, we investigated RNA-binding proteins (RBPs) that might dictate tmCSF1 expression. Van Nostrand *et al.* performed transcriptomics on two human cell lines following KD of 263 RBPs individually^34^. Analysis of the datasets from this report revealed that KD of 34 RBPs decreased *tmCSF1*-to-*sCSF1* ratio by more than 2-fold in both cell lines (Extended Data Fig. 5a). Among these RBPs, PPIL4 and MTPAP bound to CSF1 RNA, based on enhanced crosslinking and immunoprecipitation (eCLIP) sequencing performed in the same report (Extended Data Fig. 5b). Their binding sites were close (within 800 bp) to the alternative splicing acceptor site that produces the *tmCSF1* transcript, implicating PPIL4 and MTPAP in splicing-mediated control of tmCSF1 expression. We tested the roles of PPIL4 and MTPAP under lysosomal stress by knocking them down and found that both were required for stress-induced *tmCsf1*-skewing (Extended Data Fig. 5c). Expression of PPIL4 and MTPAP was upregulated by LLOMe in a MITF-dependent manner (Extended Data Fig. 5d), indicating that they are stress-responsive RBPs that promote *tmCsf1*-skewing.

### tmCSF1 guides protein aggregates to the plasma membrane

We reasoned that protein aggregates should be spatially associated with tmCSF1 at the plasma membrane to be specifically targeted by macrophages for trogocytic removal. To test this, co-localization of protein aggregates and tmCSF1 at the plasma membrane was evaluated by total internal reflection fluorescence (TIRF) microscopy, which detects fluorescent signals within 100–200 nm of the cell surface. MEFs from sCSF1 KO mice^35^ were used to specifically detect tmCSF1 with an anti-CSF1 antibody. We found that tmCSF1 formed clusters that were partially, but significantly, co-localized with protein aggregates (Fig. 4a, b). In tmCSF1 KO MEF^Baf^ ^A1^, however, fewer protein aggregates were detected (Fig. 4a, c), while flow cytometric analysis showed that total amounts of protein aggregates were similar between sCSF1 KO and tmCSF1 KO MEF^Baf^ ^A1^ (Extended Data Fig. 6a). Consistently, under TIRF microscopy, we observed cluster formation of transduced tmCSF1 and its co-localization with protein aggregates upon Baf A1 treatment (Fig. 4d, e). In parallel, tmCSF1 transduction enhanced protein aggregate localization at the plasma membrane (Fig. 4d, f) without increasing total amount of cellular protein aggregates (Extended Data Fig. 6b). To determine whether tmCSF1 is physically associated with protein aggregates, FLAG-tagged tmCSF1 (FLAG-tmCSF1) was expressed in cells and, after Baf A1 treatment, pulled down from the membrane fraction of cell lysate (Fig. 4g, h). Flow cytometric analysis showed that protein aggregates were co-precipitated with FLAG-tmCSF1 (Fig. 4i-k). Although protein aggregates were also detected upon pull-down with the control antibody, the amount was much lower than that obtained with anti-FLAG antibody (Fig. 4k), suggesting the specific association between tmCSF1 and protein aggregates. Swapping the cytoplasmic tail of FLAG-tmCSF1 with that of CD4 resulted in abrogation of protein aggregate co-precipitation with tmCSF1 (Fig. 4h, j, k), indicating that the cytoplasmic tail of tmCSF1 is required for its association with protein aggregates. This association appears to facilitate targeted uptake of protein aggregates by macrophages via trogocytosis.

**Fig. 4.**
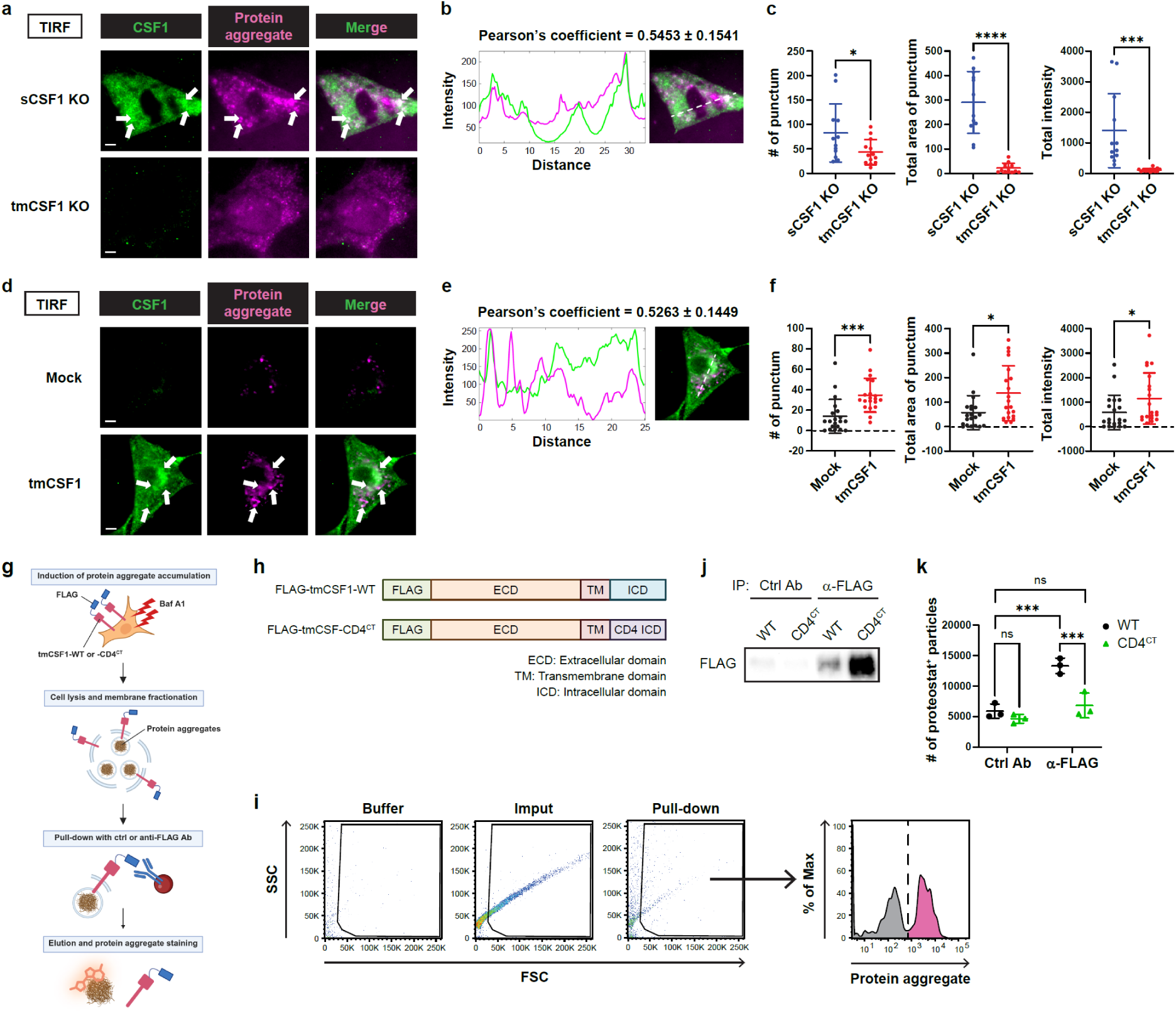
tmCSF1 spatially associates with protein aggregates via its cytosplasmic tail. a-f, TIRF microscopic analysis. **a,** Representative images of sCSF1 KO and tmCSF1 KO MEF^Baf^ ^A1^. White arrows indicate the areas where CSF1 and protein aggregates are co-localized. **b,** Correlation coefficient of CSF1 and protein aggregate signals in sCSF1 KO MEF^Baf^ ^A1^ (n = 12). Plot profile histogram shows the fluorescence intensities of both signals along the white-dotted line in the right panel. **c,** Number, total area and total intensity of protein aggregate puncta in sCSF1 KO (n = 12) and tmCSF1 KO (n = 13) MEF^Baf^ ^A1^. **d,** Representative images of Baf A1-treated mock-and tmCSF1-transduced cells. White arrows indicate the areas where CSF1 and protein aggregates are co-localized. **e,** Correlation coefficient of CSF1 and protein aggregate signals in Baf A1-treated tmCSF1-transduced cells (n = 20). Plot profile histogram shows the fluorescence intensities of both signals along the white-dotted line in the right panel. **f,** Number, total area and total intensity of protein aggregate puncta in Baf A1-treated mock-(n = 20) and tmCSF1-transduced cells (n = 20). **g,** Schematic of the experimental process for immunoprecipitation of FLAG-tagged tmCSF1 (FLAG-tmCSF1) from plasma membrane lysates of protein aggregate-harboring cells. **h,** Domain structures of FLAG-tmCSF1-WT and FLAG-tmCSF1-CD4^CT^. **i,** Flow-cytometric gating strategy to detect protein aggregate particles after the process described in (**g**). **j, k,** Immunoblot images of FLAG (**j**) and numbers of protein aggregate particles (**k**) pulled down with control and anti-FLAG antibody from plasma membrane lysates of Baf A1-treated cells transduced with FLAG-tmCSF1-WT or FLAG-tmCSF1-CD4^CT^ (n = 3 per group in **k**). Scale bars, 5 μm. **g,** Created with BioRender.com. Correlation coefficient and graphs show mean ± s.d. **c, f,** Two-tailed unpaired Student’s t test. **k,** Two-way ANOVA with Sidak’s multiple-comparison test. \**P* < 0.05, \*\*\**P* < 0.001, \*\*\*\**P* < 0.0001; ns, not significant.

### Protein aggregate-laden lysosomal vesicles are transported along microtubule to associate with tmCSF1

Electron microscopic observation of MEF^Baf^ ^A1^ revealed electron-dense conglomerates that appeared encompassed by two membranes (Fig. 5a). In addition, the protein aggregates were surrounded by LC3-positive membranes (Fig. 5b), indicating that autophagosomes encapsulate protein aggregates^36^. However, the aggregates-containing vesicles were also positive for LAMP1 and CD81 (Fig. 5b). These profiles resemble the features of amphisome-derived autophagolysosomes (ADAs, as we refer to them here) and lysosome-related organelles (LROs)^37,38^. Protein aggregates were increasingly associated with ADA/LRO-like vesicles over the course of Baf A1 treatment (Fig. 5c). MEF^Baf^ ^A1^-derived CD81 was co-localized with protein aggregates engulfed by macrophages (Fig. 5d), suggesting the transfer of aggregate-laden ADA/LRO-like vesicles upon trogocytosis.

**Fig. 5.**
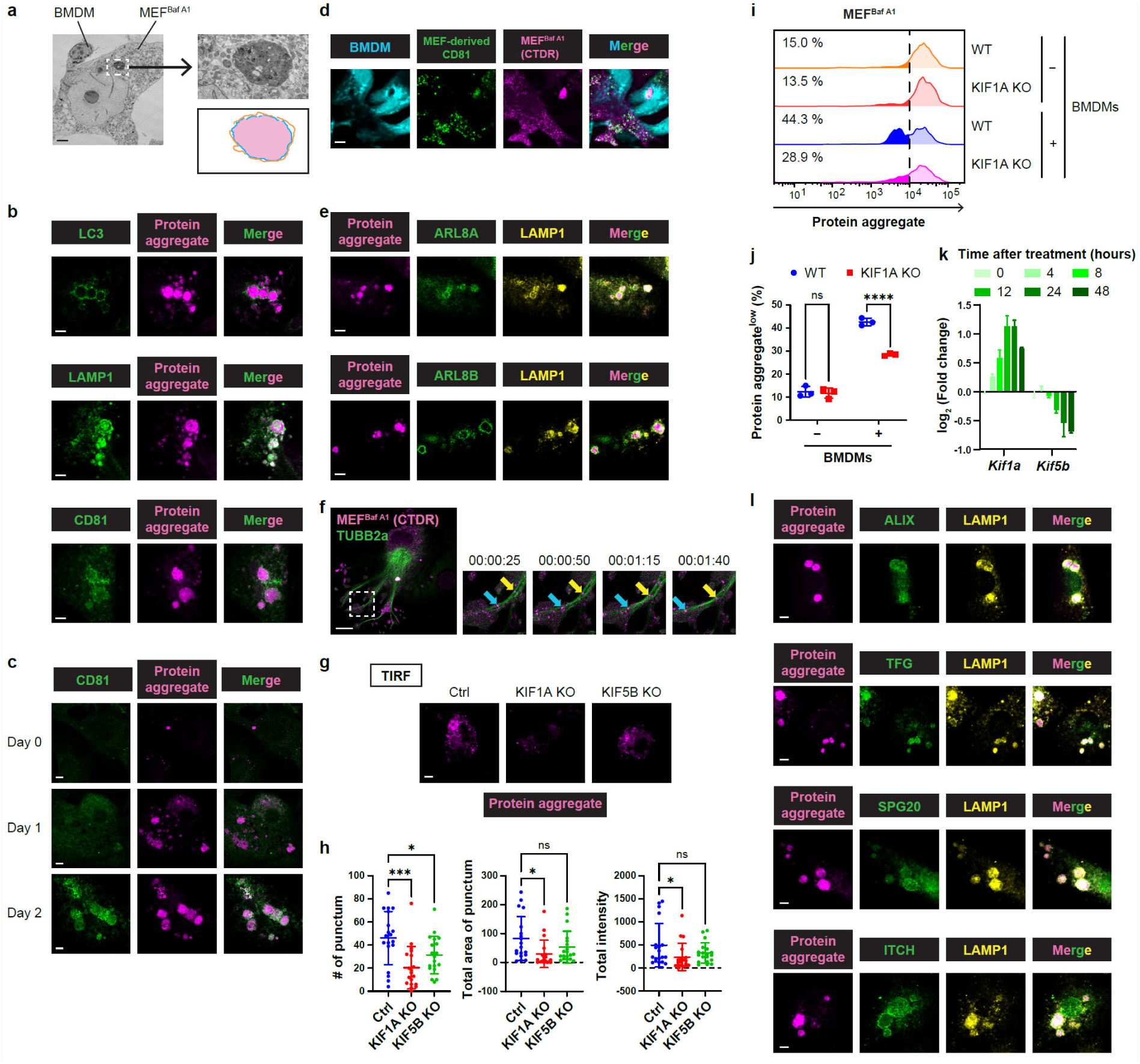
Protein aggregates in amphisome-derived autophagosome/lysosome-related organelle-like vesicles are transported to the plasma membrane along microtubules. a, Representative EM image of co-cultured BMDMs and MEF^Baf^ ^A1^. The upper-right panel displays a zoomed image of the area indicated by the white-dotted box in the left panel. The lower-right panel shows schematics of electron-dense area (red-shaded) and membranes surrounding it (blue and orange lines) in the upper-right panel. **b,** Representative confocal images of MEF^Baf^ ^A1^. Protein aggregates are zoomed in. **c,** Representative confocal images of MEFs treated with Baf A1 for the indicated times. Protein aggregates are zoomed in. **d,** Representative confocal images of co-cultured BMDMs and MEF^Baf^ ^A1^. MEF^Baf^ ^A1^ was transfected with fluorescently labelled CD81 before co-culture. **e,** Representative confocal images of MEF^Baf^ ^A1^. Protein aggregates are zoomed in. **f,** Representative time-lapse confocal images of MEF^Baf^ ^A1^. The left panel shows the image at 00:00:00. The right four panels show zoomed images of white-dotted areas in the left panel. Arrows indicate CTDR puncta that move along microtubule. Images taken every 25 sec are shown. **g,** Representative TIRF images of Ctrl, KIF1A KO and KIF5B KO MEF^Baf^ ^A1^. **h,** Number, total area and total intensity of protein aggregate puncta in (**g**) (n = 19 per group). **i, j,** Amount of protein aggregates in WT or KIF1A KO MEF^Baf^ ^A1^ after culture alone or co-culture with BMDMs (n = 3 per group). Shown are the representative histograms (**i**) and percentages (**j**) of MEFs with lower intensity of protein aggregate-staining dye than that indicated by the black-dotted line in (**j**). **k,** Changes of *Kif1a* and *Kif5b* expression in MEFs after Baf A1 treatment, determined by RNA-seq analysis (n = 2 per group). **l,** Representative confocal images of MEF^Baf^ ^A1^. Protein aggregates are zoomed in. Scale bars, 4 μm (**a**), 7.5 μm (**b, c, e, l**), 20 μm (**d, f**), 5 μm (**g**). Graphs show mean ± s.d. **h,** One-way ANOVA with Dunnett’s multiple-comparison test. **j,** Two-way ANOVA with Sidak’s multiple-comparison test. \**P* < 0.05, \*\*\**P* < 0.001, \*\*\*\**P* < 0.0001; ns, not significant.

Amphisome is a hybrid vesicle produced by the fusion of autophagocome and late endosome, which ultimately fuses with lysosome to form ADA for cargo degradation or releases cargo extracellularly^37^. In contrast, LRO is functionally specialized for storage and secretion of cargo, rather than degradation^39^. It generally contains molecules specifically expressed in a given cell type; for example, melanin, which is transferred from melanocytes to neighboring keratinocytes via trogocytosis-like mechanisms^40^. Overall, the functionality of ADAs and LROs requires regulated transport systems^37,39^. On this basis, we hypothesized that protein aggregates in ADA/LRO-like vesicles are actively transported to the plasma membrane for physical association with tmCSF1. Anterograde transport of lysosomal vesicles is driven by the BLOC-one-related complex (BORC)^41^. Protein aggregates in MEF^Baf^ ^A1^ were positive for ARL8A and ARL8B (Fig. 5e), which are recruited by BORC and tether client vesicles to kinesin-1 and-3^41^. To determine whether the localization of aggregate-laden vesicles was controlled by microtubules, MEFs were transfected with fluorescently tagged tubulin, TUBB2a, and treated with Baf A1. Time-lapse imaging showed multiple CTDR-labelled aggregates moving along TUBB2a-labelled microtubules (Fig. 5f, Supplementary Video 5). In addition, we knocked out KIF1A and KIF5B, ARL8-interacting motor proteins belonging to kinesin-3 and kinesin-1 families, respectively^42^, in MEFs (Extended Data Fig. 6c) and induced protein aggregate accumulation with Baf A1. TIRF microscopy analysis revealed that the amount of protein aggregates at the plasma membrane was significantly decreased in KIF1A KO MEF^Baf^ ^A1^, while only a mild reduction was observed in KIF5B KO cells (Fig. 5g, h). We did not detect significant changes in the total amounts of protein aggregates and CSF1 expression at the plasma membrane upon KIF1A and KIF5B deficiency (Extended Data Fig. 6d, e). Moreover, KIF1A KO cells showed a mild reduction of protein aggregate amount upon co-culture with macrophages (Fig. 5i, j), suggesting the requirement of KIF1A-mediated anterograde transport in trogocytic aggregate removal. Consistent with KIF1A-biased mechanism of protein aggregate transport, *Kif1a* was upregulated whereas *Kif5b* was downregulated upon Baf A1 treatment (Fig. 5k). Thus, the anterograde transport system is reprogrammed upon stress to favor the KIF1A-microtubule machinery for aggregate translocation to the plasma membrane. The vesicles enclosing protein aggregates were positive for ALIX, TFG, SPG20 and ITCH, which have been reported to be recruited to damaged lysosomes (Fig. 5l)^43–45^. These results support the requirement of a shift in intracellular transport program toward the KIF1A-microtubule pathway, given that KIF1A is activated in response to Ca^2+^ ^46^, which is released upon lysosomal damage^43^.

We next hypothesized that the vesicle transport system would need to be directed to tmCSF1 on the plasma membrane, to enable tmCSF1-dependent protein aggregate removal through trogocytosis. We further reasoned that this targeting should require tethering molecules that link the microtubule plus end and the short unstructured cytoplasmic tail of tmCSF1. Sequence analysis revealed a conserved class II PDZ-binding motif (Φ-X-Φ-COOH) at the C-terminus of tmCSF1 cytoplasmic tail (Fig. 6a). MAGUK family proteins, essential molecules for spatial stabilization of plasma membrane proteins and for maintenance of cell polarity, contain PDZ domains and link cytoskeleton to cytoplasmic tail of transmembrane proteins^47^. In our RNA-seq datasets, MPP1 was the most highly expressed among MAGUK proteins and was upregulated upon Baf A1 treatment in MEFs (Fig. 6b). TIRF microscopy revealed that MPP1 was co-localized with tmCSF1 and protein aggregates at plasma membrane (Fig. 6c and d). In the absence of MPP1, the plasma membrane localization of protein aggregates was markedly decreased (Fig. 6c, e and Extended Data Fig. 6f), while total amounts of protein aggregates and plasma membrane tmCSF1 expression were not undermined (Extended Data Fig. 6g, h). Accordingly, MPP1 deficiency blunted macrophage-mediated reduction of protein aggregate amount in MEF^Baf^ ^A1^ (Fig. 6f, g). To test physical interaction of tmCSF1 with MPP1, tmCSF1 was pulled down from the membrane fraction of FLAG-tmCSF1-transduced cell lysate. We detected co-immunoprecipitation of MPP1, which was elevated upon Baf A1 treatment, while deletion of PDZ-binding motif in tmCSF1 abrogated MPP1 pull-down with tmCSF1 (Fig. 6h, i). These results suggest that MPP1 is required for association of protein aggregate-containing vesicles with tmCSF1. Together, these data highlight a stress-induced microtubule system that contributes to anterograde transport of protein aggregates and their specific association with tmCSF1 for trogocytic aggregate removal.

**Fig. 6.**
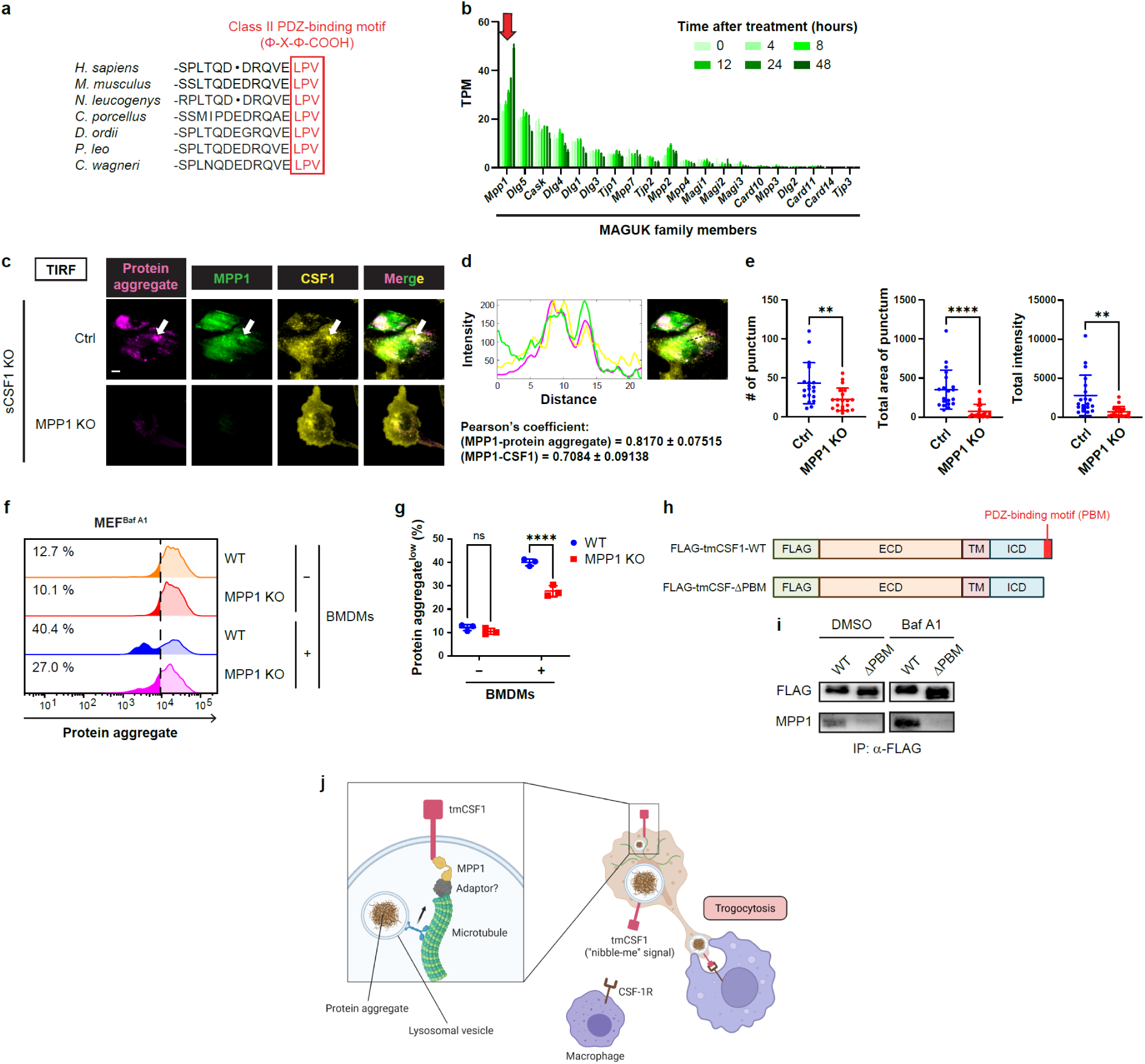
tmCSF1 requires MPP1-mediated scaffolding to spatially associate with protein aggregates. a, C-terminal amino acid sequences of tmCSF1 in the indicated mammals. Red letters indicate PDZ-binding motif. **b,** Expression change of MAGUK family members in MEFs after Baf A1 treatment, determined by RNA-seq analysis (n = 2 per group). The red arrow indicates the expression level of MPP1. **c,** Representative TIRF microscopy images of Ctrl and MPP1 KO sCSF1-deficient MEF^Baf^ ^A1^. White arrows indicate the areas where protein aggregates, MPP1 and CSF1 are co-localized. Scale bars, 5 μm. **d,** Plot profile histogram showing the fluorescence intensities of protein aggregate, MPP1 and CSF1 signals along the black-dotted line in the right panel. Shown below are correlation coefficients of MPP1 and protein aggregate signals and MPP1 and CSF1 signals in Ctrl sCSF1-deficient MEF^Baf^ ^A1^ (n = 20). **e,** Number, total area and total intensity of protein aggregate puncta in (**c**) (n = 20 per group). **f, g,** Amount of protein aggregates in WT or MPP1 KO MEF^Baf^ ^A1^ after culture alone or co-culture with BMDMs. Shown are the representative histograms (**f**) and percentages (**g**) of MEFs with lower intensity of protein aggregate-staining dye than that indicated by the black-dotted line in (**f**) (n = 3 per group). **h,** Domain structures of FLAG-tmCSF1-WT and FLAG-tmCSF1-ΔPBM. **i,** Immunoblot images of FLAG and MPP1 pulled down with anti-FLAG antibody from plasma membrane lysates of DMSO-or Baf A1-treated cells transduced with FLAG-tmCSF1-WT or FLAG-tmCSF1-ΔPBM. **j,** Summary schematic, created with BioRender.com. Graphs show mean ± s.d. **e,** Two-tailed unpaired Student’s t test. **g,** Two-way ANOVA with Sidak’s multiple-comparison test. \*\**P* < 0.01, \*\*\*\**P* < 0.0001; ns, not significant.

### Protein aggregates are sequestered in lysosomal vesicles independently of canonical protein homeostasis programs

We next investigated the mechanisms underlying protein aggregate compartmentalization in ADA/LRO-like vesicles. Mass spectrometry analysis showed that proteostat-positive protein aggregates were enriched with a plethora of cytosolic proteins, particularly cytoskeletal components and cytoskeleton-associated molecules (Extended Data Fig. 7a and Supplementary Fig. 1a). Interestingly, these profiles resemble the patterns of protein aggregates accumulated in aged killifish tissues^48^. Aggregation of cytoskeleton-related proteins has been also reported in diverse proteinopathies^49–52^. Cytosolic proteins are recruited to lysosomes through diverse autophagy mechanisms^7,8^. Association with various combinations of autophagy receptors, chaperones and cofactors enables lysosomal sorting of misfolded and aggregated proteins for degradation^53^. However, none of the molecules that we tested (ubiquitin, p62, CCT2, TAX1BP1 and Hsc70) were specifically co-localized with protein aggregates in MEF^Baf^ ^A1^, except for BAG3 that surrounded protein aggregates (Extended Data Fig. 7b). Although aggresome, a protein complex designed to store protein aggregates, can also drive autophagic degradation^54^, typical aggresome markers (HDAC6, γ-tubulin and vimentin) were minimally associated with Baf A1-induced protein aggregates (Extended Data Fig. 7b). We further tested the involvement of autophagy programs by knocking out autophagy receptors p62 and CCT2, a co-factor BAG3 and a lysosomal misfolded protein importer LAMP2^7^ (Extended Data Fig. 7c).

Compartmentalization of protein aggregates in ADA/LRO-like vesicles was not impaired by any of the manipulations of these molecules, including BAG3 deletion, despite the association of BAG3 with protein aggregates (Extended Data Fig. 7b, d, e). These results suggest that aggregates targeted for trogocytosis are handled independently of previously identified autophagy pathways, although details of the mechanisms and relation to (perhaps noncanonical) autophagy remain to be elucidated.

Because ubiquitin association with protein aggregates was negligible (Extended Data Fig. 7b), a question remained as to whether proteins destined for proteasomal degradation can be targeted by trogocytosis. To address this, cells were treated with proteasome inhibitors bortezomib and carfilzomib. While chronic proteasome inhibition induced ubiquitinated protein aggregation, we did not observe detectable compartmentalization of the aggregates in ADA/LRO-like vesicles (Extended Data Fig. 7f). Proteasome inhibition significantly decreased *tmCsf*1-skewing and tmCSF1 protein expression on the cell surface (Extended Data Fig. 7g, h), suggesting that ubiquitinated protein aggregates are not the dominant targets of trogocytosis. Thus, trogocytic removal of protein aggregates appears to be distinct from ubiquitin-proteasomal system.

### Overproduced protein aggregates are handled by trogocytic protein homeostasis program

Insufficiency of cell-autonomous proteostasis can be caused not only by failed clearance of misfolded and aggregated proteins but also their excessive production. We tested whether tmCSF1-mediated trogocytosis acts on protein aggregate overload without induced impairment of lysosomal function. The polyglutamine huntingtin mutant HTT-Q103, which is widely used to experimentally model Huntington disease, forms aggregates and induces cellular damage. We found that HTT-Q103 aggregates that formed upon HTT-Q103 overexpression in tmCSF1-transduced cells were incorporated by macrophages in a manner dependent on tmCSF1-CSF1R interaction (Extended Data Fig. 8a, b). Moreover, HTT-Q103 aggregates were pulled down with tmCSF1, while a smaller number of aggregates were detected when the cytoplasmic tail of tmCSF1 was swapped with that of CD4 (Extended Data Fig. 8c, d). In addition, multiple HTT-Q103 aggregates were enclosed by CD81^+^ LAMP1^+^ ADA/LRO-like vesicles, as were the aggregates of Tau (K18-P301L/V337M), which carries human pathogenic mutations associated with neurodegenerative tauopathies^55^ (Extended Data Fig. 8e). Given that protein aggregates can trigger lysosomal damage and dysfunction^1^, these results indicate that overproduced protein aggregates can be handled by trogocytosis programs that are induced by lysosomal stress.

### CSF1R possesses phagocytic activity

CSF1R activates PI3K and MAPK signaling and promotes macrophage proliferation, differentiation, survival, and migration^32^. Our data show that macrophages co-opt CSF1R to perform trogocytosis (Extended Data Fig. 1k, l). To determine the mechanisms of uptake downstream of CSF1R, we established an *in vitro* particle-uptake system in which the biotinylated extracellular domain (ECD) of tmCSF1 was conjugated to streptavidin-coated fluorescent beads and incubated with BMDMs. Macrophages engulfed tmCSF1-tethered beads in a manner completely dependent on CSF1R in this system, as shown by abrogated uptake of beads in the presence of a CSF1R-blocking antibody (Extended Data Fig. 9a, b). The bead diameter was 1.24 μm, which was at least ten times larger than the initial size of endosomes formed by endocytosis (60-120 nm)^56^, indicating that uptake of tmCSF1-tethered beads did not simply represent receptor-mediated endocytosis. Soluble CSF1 protein did not enhance uptake of untethered beads (Extended Data Fig. 9c, d). Furthermore, while BMDMs engulfed beads upon incubation with tmCSF1 ECD and beads without pre-tethering, uptake was abrogated in the presence of streptavidin that blocked tmCSF1 ECD-bead conjugation (Extended Data Fig. 9c, d). These results indicate that enhanced bead incorporation upon tmCSF1 ligation is not explained by a possible elevation of general uptake activity in macrophages. We found that the CSF1R-PI3K-actin polymerization pathway was responsible for macrophage incorporation of tmCSF1-tethered beads, based on the findings that uptake was abrogated by the treatment with wortmannin and latrunculin A, an inhibitor of PI3K and actin polymerization, respectively (Extended Data Fig. 9e, f). This observation is consistent with previous studies showing that the PI3K-actin polymerization axis is a conserved mechanism to trigger trogocytosis in protists and mammalian cells^57,58^. CSF1R drives two uptake processes: macropinocytosis and receptor endocytosis^32,59^. Nevertheless, inhibition of macropinocytosis with DMA did not repress uptake of tmCSF1-tethered beads (Extended Data Fig. 9g, h). Although the size of tmCSF1-tethered beads exceeds that of initially formed endosomes, the mechanisms that drive receptor endocytosis may contribute to uptake of the beads. To test this possibility, we dissected the signaling pathways involved in CSF1R endocytosis. Treatment with wortmannin and latrunculin A markedly decreased CSF1R endocytosis, as evaluated by recombinant CSF1 internalization (Extended Data Fig. 9i, j). Whereas inhibition of Arp2/3, an actin nucleator required for actin filament branching^60^, with CK-666 blocked receptor endocytosis, no significant change was observed upon treatment with SMIFH2, an inhibitor of formins that nucleate actin for actin filament elongation^60^ (Extended Data Fig. 9i, j). In contrast, uptake of tmCSF1-tethered beads was not suppressed by CK-666 but was impaired by SMIFH2 (Extended Data Fig. 9k-n). Collectively, these results suggest that CSF1R exerts phagocytic activity via non-canonical but actin-dependent downstream pathways.

Our RNA-seq dataset revealed that BMDMs expressed a wide variety of formin family members (Extended Data Fig. 10a). *Diaph1,* which mediates integrin-dependent phagocytosis^61^, was among the most highly expressed formins (Extended Data Fig. 10a). We leveraged gene editing in ER-Hoxb8 bone marrow-derived cell line^62^ and established Diaph1 KO cells (Extended Data Fig. 10b). Macrophages derived from Diaph1 KO cells exhibited only a slight reduction of cell numbers after 7 days of differentiation with CSF1-containing medium (Extended Data Fig. 10c), suggesting Diaph1 is minimally involved in CSF1R signaling that soluble CSF1 activates to drive macrophage proliferation and differentiation. Nevertheless, Diaph1 KO macrophages showed reduced uptake of tmCSF1-tethered beads (Extended Data Fig. 10d, e) and of materials from MEF^Baf^ ^A^^1^ (Extended Data Fig. 10f,g). These results suggest that CSF1R-Diaph1 pathway engages in trogocytosis.

## Discussion

Protein misfolding and aggregation are inevitable perturbations that cells experience as a consequence of cell stress and errors in protein translation process. While organisms in all kingdoms of life have evolved cell-autonomous homeostasis system to remove structurally abnormal proteins, we speculated that multicellular organisms have another layer of homeostasis mediated by cell-to-cell interactions. In this study, we found that intracellular protein aggregates are removed by macrophage trogocytosis (Fig. 6j). This program requires macrophage recognition of tmCSF1 on protein aggregate-harboring cells, which is upregulated through lysosomal stress response pathways. Protein aggregates are loaded into lysosomal vesicles and transported to the plasma membrane along microtubules to physically associate with the cytoplasmic tail of tmCSF1, facilitating targeted uptake of the aggregates by macrophages. This study sheds light on a protein homeostasis program that operates through cell-to-cell interactions upon insufficient clearance of protein aggregates.

Seminal studies have shown a general requirement of neighboring cell-derived assistance for cellular homeostasis upon stress exposure. Neighboring cells supply miRNAs, proteins and metabolites to stressed cells via extracellular vesicles and tunneling nanotubes, promoting stress adaptation of the recipient cells^63–66^. Recent studies also demonstrated the transfer of functional mitochondria through similar mechanisms to cells undergoing mitochondrial stress^66,67^. Removal of extracellular stress sources is also programmed in non-cell-autonomous manner, exemplified by microbial clearance via immune cells that are recruited by infected cells^10^. In contrast, whether neighboring cells assist the clearance of intracellular stress sources has been understudied. Here, our findings indicate that intracellular protein aggregates can be removed in a non-cell-autonomous manner via macrophage trogocytosis.

Our data show that protein aggregates in ADA/LRO-like vesicles are targeted for trogocytosis-mediated removal. Whereas autophagosomes are double-membrane vesicles, autophagolysosomes, including ADAs, have a single membrane because autophagosome-derived inner membrane is degraded after fusion with lysosomes^6,8,36^. LROs, which are generated from endosomal vesicles and Golgi apparatus, are also generally single-membrane vesicles^39^. Nevertheless, the ADA/LRO-like vesicles containing trogocytosis-targeted aggregates exhibited a double-membrane structure. One possible notion raised by these results is that the amphisome-derived inner membrane persists after fusion with lysosome, due to lysosomal dysfunction and damage caused by protein aggregate accumulation as observed by us and reported previously^1^. However, our data show that vesicular compartmentalization of protein aggregates is independent of canonical autophagy, suggesting that currently uncharacterized molecular pathways recruit protein aggregates into amphisomes. Elucidation of the underlying mechanisms of this process will provide further insight into how cells handle protein aggregates for subsequent trogocytosis.

Our findings suggest that the coordination of cell-autonomous programs and trogocytosis is designed to maximize cellular homeostasis. We found that tmCSF1-skewing and sequestration of aggregates in ADA/LRO-like vesicles became prominent upon chronic treatment with the agent that induces protein aggregate accumulation. In contrast, cell-autonomous protein homeostasis programs are acutely driven by protein misfolding and aggregation^6,8,36^. Upon chronic accumulation, protein aggregates can induce lysosomal damage, compromising cell-autonomous systems for protein degradation. While cells can release protein aggregates upon lysosomal damage^9^, excessive accumulation of extracellular aggregates is likely to be detrimental to tissue homeostasis, as suggested by models of neurodegenerative diseases^68–70^. Thus, we propose that trogocytosis is not merely redundant with cell-autonomous systems; rather, it reflects a temporal shift in the overall protein homeostasis program, becoming increasingly important as cell-autonomous defenses fail under chronic stress. Given that lysosomal damage induces tmCSF1 expression, senescent cells, which undergo a decline in lysosomal function and accumulate misfolded/aggregated proteins, may also require trogocytosis to maintain their fitness. Our study highlights the significance of trogocytosis in protein homeostasis and suggests that further mechanistic understanding of trogocytosis regulation could lead to new therapeutic strategies for proteinopathies — particularly at the later stage, when most diagnoses occur.

## Methods

### Cells

MEFs and BMDMs were prepared and maintained as described previously^71^. One day before BMDMs were used for experiments, the medium was switched to complete RPMI [RPMI 1640 (Corning) supplemented with 10% heat-inactivated FBS (Gibco), 2 mM L-glutamine (Gibco), 1 mM sodium pyruvate (Gibco), 10 mM HEPES (Gibco), and 200 U/ml penicillin/streptomycin (Gibco)]. YUMMER1.7 cells were kindly provided by M. Bosenberg (Yale University) and cultured as described previously^72^. NIH/3T3 cells were maintained in complete Dulbecco’s modified Eagle’s medium (DMEM) [Dulbecco’s modified Eagle’s medium (Sigma Aldrich) supplemented with 10% heat-inactivated FBS, 2 mM L-glutamine, 1 mM sodium pyruvate, 10 mM HEPES, and 200 U/ml penicillin/streptomycin]. ER-Hoxb8 bone marrow-derived cells were kindly provided by J. Cox (University of California, Berkeley) and cultured as described previously^62^. Macrophage differentiation was performed using the same protocol used for BMDM preparation.

YUMMER1.7-derived transduced cells and NIH/3T3-derived transduced cells were seeded at 8 × 10^4^ cells on 24-well plate and cultured overnight before use unless otherwise indicated. MEFs were seeded at 4 × 10^4^ cells on 24-well plate or 4-well chamber slide (Thermo Fisher Scientific, 154917) and cultured overnight before treatment unless otherwise indicated.

### Animals

The following mouse strains were obtained from The Jackson Laboratory: C57BL/6J (#000664), TREM2 KO (#027197) and CD11b KO (#003991). tmCSF1 KO and sCSF1 KO mice were established as described previously^31,35^. All animal procedures were performed in accordance with institutional regulations after protocol review and approval by Yale University’s Institutional Animal Care and Use Committee.

### Stress induction *in vitro*

Cells were treated with 0.1 μM Baf A1 (Cayman Chemical, 11038) for 48 hours unless otherwise stated. Cells were then washed three times with DMEM before harvesting or were subjected to EasySep Dead Cell Removal (Annexin V) Kit (Stemcell Technologies, 17899) to remove dead cells. Treatment with 1 mM LLOMe (Cayman Chemical, 16008) was performed for 6 hours. Treatments with 20 nM bortezomib (Cayman Chemical, 10008822) and 20 nM carfilzomib (Cayman Chemical, 17554) were performed for 24 hours unless otherwise stated.

### RNA-sequencing (RNA-seq)

RNA extraction and sequencing of Baf A1-treated MEFs were performed by Plasmidsaurus. Total RNA in BMDMs was isolated using RNeasy Mini Kit (QIAGEN), followed by library construction with NEBNext Ultra II DNA Library Prep Kit for Illumina (New England Biolabs). Sequencing was performed with NextSeq 500 (Illumina). Sequencing reads were aligned to *Mus musculus* transcriptome GRCm38 obtained from Ensembl (ftp://ftp.ensembl.org/pub/release-90/fasta/mus_musculus/cdna/s), using Kallisto v0.46.1 with k-mer index 25 and 100 bootstrapping ^73^. Transcript expression was quantified as transcripts per million (TPM).

For analysis of the datasets generated by Van Nostrand *et al.*^34^, accession IDs were retrieved from https://www.encodeproject.org/experiments for transcript quantification files with data mapped to the GRCh38 assembly and genome annotation version V29. These files were downloaded to extract TPM values for CSF1 transcript isoforms across all experiments. For *sCSF1*, TPM values of the three transcript variants were summed within each replicate. *tmCSF1*, encoded by a single transcript, required no aggregation. The *tm/s CSF1* TPM ratio was first calculated within each replicate, and then the fold difference in this ratio was calculated between knockdown (KD) and control experiments for each RNA-binding protein (RBP) and cell line. Because the replicates were not paired, the KD replicate values of the *tmCSF1/sCSF1* ratio were divided by the mean of the *tmCSF1/sCSF1* ratio of the control replicates to calculate the fold change. For categorizing upregulated and downregulated genes, mean fold change (mFC) was calculated. log_2_ mFC > 1.5 and log_2_ mFC < −1.5 were considered as upregulated and downregulated, respectively.

### Quantitative polymerase chain reaction (qPCR)

Total RNA from cells was purified using RNeasy Mini Kit (Qiagen) or RNA-Bee (Tel-Test). qPCR was performed as described previously^74^. Transcript levels of target genes were normalized to *Rpl13a*. Primers used for qPCR are listed in Supplementary Table 1.

### Enzyme-linked immunosorbent assay (ELISA)

Culture supernatants were centrifuged at 10,000 rpm for 10 min. CSF1 in the supernatant was quantified with Mouse M-CSF Quantikine ELISA Kit (R & D Systems, MMC00). For ADAM17 inhibition, cells were cultured in the presence of 10 μM batimastat (Cayman Chemical, 14742) for 24 hours before supernatant collection. When CSF1 from Baf A1-treated MEFs was measured, concentrations were normalized to the area under the curve of cell growth curves generated from cell numbers counted at 0, 24, and 48 hours after treatment.

### Flow cytometry

Cells were incubated with anti-CD16/CD32 antibody (clone 93, eBioscience, 14-0161-86) to block Fc receptor and subsequently stained with the antibodies listed in Supplementary Table 2. Dead cell staining was performed with propidium iodide (Thermo Fisher Scientific, 00-6990-50). Cell numbers were determined with 123count eBead Counting Beads (Thermo Fisher Scientific, 01-1234-42). Flow cytometry was performed on BD LSR II. Data were analyzed with FlowJo software (FlowJo LLC).

### Plasmid cloning

tmCSF1, sCSF1, FLAG-tmCSF1-WT, FLAG-mCSF1^CD4CT^ and FLAG-tmCSF1-ΔPBM were cloned into pMSCVpuro-IRES-GFP (pMIG)^75^. Empty pMIG was used for mock transduction. mCherry-HTT-Q103 were cloned into pTetOne-blast generated by replacing puromycin resistance cassette in pLVX-TetOne-puro (TaKaRa) with blasticidin-resistance cassette. All DNA ligations into plasmids were performed with NEBuilder HiFi DNA Assembly Master Mix (New England Biolabs). All constructs were sequenced before use. Vector maps are available on request. Detailed methods are described in Supplementary Methods.

### Gene transduction

Retroviral transduction of CSF1-deficient YUMMER1.7 cells and CSF1-deficient NIH/3T3 cells with pMIG-backbone vectors was performed as described previously^75^. For lentivirus production, HEK293T cells were co-transfected with pTetOne-blast backbone vectors and packaging vectors psPAX2 and VSV-G, the latter two of which were kindly provided by C. Wilen (Yale University). Culture medium was replaced the next day. Two days later, culture supernatant was collected and passed through 0.45 μm filter. After three-fold concentration with Lenti-X Concentrator (TaKaRa), viral supernatant was added to cells with 8 μg/ml polybrene (Sigma Aldrich, TR-1003). Culture plates were centrifuged at 1,200 × g for 90 min at room temperature, followed by replacement of the viral supernatant with fresh medium. Two days later, selection of transduced cells was initiated with 50 μg/ml blasticidin and continued for 7 days. Cells transduced with pTetOne-blast-based vectors were maintained in medium containing tetracycline-free FBS (Thermo Fisher Scientific, A4736101). Transfection of MEFs was performed with Andwin Scientific JETPRIME DNA/SIRNA RGNT (Fisher Scientific, NC0291070). Plasmids used are listed in Supplementary Table 3.

### Gene knockdown

MEFs were transfected with 12.5 pmol siRNAs using Lipofectamine RNAiMAX Transfection Reagent (Thermo Fisher Scientific). After 24 hours, cells were treated with chemicals and subjected to qPCR analysis. siRNAs used are listed in Supplementary Table 4.

### Gene deletion

DNA sequences encoding *Csf1*-targeting guide RNAs (5′-CACCGTCTTCGGTAAGCTGCAACCG-3′ and 5′-AAACCGGTTGCAGCTTACCGAAGAC-3′) were designed with CHOPCHOP (https://chopchop.cbu.uib.no/). These fragments were annealed and cloned into pSpCas9(BB)-2A-GFP (PX458) (Addgene). Vector was transfected into YUMMER1.7 or NIH/3T3 cells with FuGENE HD Transfection Reagent (Promega) or Lipofectamine 2000 Transfection Reagent, respectively. After an overnight incubation, single-cell sorting was performed to collect GFP-expressing transfected cells with FACSAria (BD Biosciences). CSF1 deficiency of clones was determined by quantifying of CSF1 in the culture supernatant and tmCSF1 on the cell surface.

A pair of single-guide RNAs listed in Supplementary Table 5 were used for gene deletion in MEFs, following the protocols described previously^76^.

For Diaph1 deletion, guide RNA-encoding oligonucleotides were designed (gRNA#3172) and cloned into pLV lentivirus vector with U6 promoter and TagBFP by VectorBuilder. Scramble guide RNA (gRNA#1) was cloned into the same vector as a control. Lentiviral transduction into ER-Hoxb8 cells was performed as described above. Two days after transduction, BFP-positive cells were sorted using Bigfoot Cell Sorter (Thermo Fisher Scientific) with 100 μm nozzle.

### eCLIP data analysis

eCLIP narrowpeak files (GRCh38 genome version) were downloaded from the ENCODE project website (www.encodeproject.org) for PPIL4 and MTPAP (experiments: ENCSR197INS and ENCSR200DKE, respectively). Narrowpeak files were converted to bed files using awk and the pValue for each peak was retained in the bed score column. Peak positions on genome in each replicate experiment were displayed with UCSC Genome Browser (https://genome.ucsc.edu/). Peaks appearing in both replicates were determined using intersectBed (Bedtools suite^77^) and the amount of overlap was reported using the “-wo” flag. Replicate peaks with at least 4 nucleotides of overlap were selected using awk (awk ‘$13 >= 4’). Peaks were associated with genes using intersectBed with Gencode genes downloaded from UCSC table browser (GRCh38/hg38 Gencode V46). Ensembl transcript IDs were converted to Ensembl gene IDs using Python.

### Confocal live imaging

YUMMER1.7-derived mock-and tmCSF1-transduced cells were labelled with CellTracker Orange CMRA (Thermo Fisher Scientific, C34551) and CellTracker Deep Red (CTDR; Thermo Fisher Scientific, C34565), respectively. In 35-mm glass-bottom dish (Mattek, P35G-1.5-14-C), 1 × 105 mock-transduced cells and 1 × 10^5^ tmCSF1-transduced cells were mixed and co-cultured overnight. BMDMs were added at 5 × 10^5^ cells after labelled with CellTrace Violet (CTV; Thermo Fisher Scientific, C34557).

MEFs were labelled with CTDR and seeded at 1.2 × 10^5^ cells on 35-mm glass-bottom dish. Two days after Baf A1 treatment, MEFs were co-cultured with CTV-labelled 5 × 10^5^ BMDMs.

Time-lapse imaging was initiated 2 hours after co-culture of BMDMs and transduced cells/MEFs was started. Images were taken every 2 min using Leica TCS SP5 or Leica Stellaris 8 confocal microscope.

### Cellular material uptake assay

Transduced cells or MEFs were labelled with CTV or Vibrant DiD (Thermo Fisher Scientific, V22887). In non-tissue culture-treated (non-TC) 96-well plate, 5 × 10^4^ BMDMs were co-cultured with 1 × 10^4^ CTV-labelled cells or 0.625 × 10^3^ DiD-labelled cells for 2 hours. For quantification of protein aggregates in macrophages, 5 × 10^4^ unlabelled MEFs were co-cultured. For contactless co-culture, BMDMs were seeded in the bottom well of HTS Transwell-96 Permeable Support with 8.0 µm Pore Polyester Membrane (Corning, 3374). One hour later, labelled transduced cells or MEFs were added on the upper well and co-cultured for 2 hours. To block CSF1R ligation, BMDMs were incubated with anti-CD16/CD32 antibody at 1/20 dilution for 5 min and subsequently with 250 μg/ml isotype control antibody (clone 2A3, Bio X Cell, BE0089) or anti-CSF1R antibody (clone AFS98, Bio X Cell, BE0213) for 30 min at 4°C. Cell suspensions were then diluted with RPMI such that the final concentration of the antibodies during co-culture was 10 μg/ml. For competitive inhibition of CSF1R ligation, BMDMs were incubated with 50 ng/ml recombinant CSF1 or the conditioned medium from YUMMER1.7-derived sCSF1-transduced cells for 2 hours prior to co-culture with labelled transduced cells. Conditioned medium was prepared by collecting culture supernatant 24 hours after plating sCSF1-transduced cells. For inhibitor treatment, BMDMs were seeded in 96-well plates and pretreated with 5 μM R406 (Cayman Chemical, 11422), 5 μM IPI-145 (Cayman Chemical, 16800), 5 μM tenalisib (Cayman Chemical, 30949), 1 μM wortmannin (Cayman Chemical, 10010591) or 6 μM BLZ-945 (Cayman Chemical, 19626) for 30 min. Co-culture with labelled MEFs was subsequently performed in the continued presence of the same concentrations of inhibitors. To mask phosphatidylserine, Baf A1-treated MEFs were incubated with 0.125 mg/ml recombinant Annexin V (BD Biosciences, 556416) in annexin-binding buffer (10 mM HEPES, 140 mM NaCl and 2.5 mM CaCl_2_, pH 7.4) for 20 min at 4°C. Subsequently, cells were diluted with DMEM such that the final concentration of Annexin V was 5 μg/ml during co-culture with BMDMs. After target cells were removed by three washes with medium, macrophages were harvested with PBS containing 5 mM EDTA. Percentages of CTV-or DiD-positive cells among live CD11b^+^ F4/80^+^ macrophages were determined by flow cytometry. The upper boundary of the gate was defined based on the fluorescence intensity of labelled target cells to exclude macrophage-target cell doublets. Comparable numbers of macrophages and target cells across experimental conditions were confirmed using 123count eBead Counting Beads.

### Co-culture system

For confocal microscopic analysis, MEFs were treated with Baf A1 in 4-well chamber and co-cultured with 2 × 10^5^ CTV-labelled BMDMs for 4–6 hours. Pre-staining of lysosomes in BMDMs was performed with 50 nM LysoTracker Red DND-99 (Thermo Fisher Scientific, L7528). For flow cytometric quantification of protein aggregate in MEFs, 3 × 10^4^ MEFs were cultured overnight in 12-well plate. After Baf A1 treatment, 4.5 × 10^5^ BMDMs were added. Cells were collected 6 hours later. For the assessment of cellular fitness upon co-culture, 1.2 × 10^5^ MEFs were seeded in 6-well plate and cultured overnight. Following Baf A1 treatment, cells were cultured with or without 6 × 10^5^ BMDMs for 24 hours. Cells were harvested and subjected to EasySep Mouse CD11b Positive Selection Kit II (Stemcell Technologies, 18970). Magnetic separation was performed three times in total. Negatively selected MEFs were seeded at 3 × 10^4^ cells in 24-well plate and cultured for 24 hours. Dead cell proportion was quantified with flow cytometry. For analysis of tmCSF1 and CSF1R expression upon co-culture, 1.5 × 10^5^ BMDMs and 3 × 10^4^ YUMMER1.7-derived mock-or tmCSF1-transduced cells were mixed and cultured in 24-well plate. Cells were collected 4 hours later and analyzed with flow cytometry. Cells transduced with HTT-Q103 were seeded at 5 × 10^4^ cells in 4-well chamber slides and treated with 0.1 μg/ml doxycycline (Sigma Aldrich, D9891) for 24 hours to induce mutant protein expression. After cells were rinsed to remove doxycycline, 1 × 10^5^ CTV-labelled BMDMs were added and incubated for 6 hours in medium containing tetracycline-free FBS for confocal microscopy.

### Focused Ion Beam-Scanning Electron Microscopy (FIB-SEM)

Baf A1-treated MEFs on No. 1.5 thickness 35 mm^2^ glass-bottom dishes were co-cultured with BMDMs for 30 minutes. Cells were prepared for enhanced FIB-SEM (eFIB-SEM) imaging essentially as previously described^78^. Briefly, the regions of interest (ROIs) were trimmed following coating and polymerization of the surface with Durcupan ACM resin (Sigma Aldrich). The embedded sample was scanned by X-ray microtomography on a Zeiss Xradia VersaXRM 730 to locate the ROIs. The block was trimmed to the ROI on a UC7 ultramicrotome (Leica Microsystems). The specimen was imaged on Zeiss Merlin FIB-SEM system. Data was acquired at a scan rate of ∼500 kHz (pixel dwell time ∼2 µs) with electron beam at 1.2kV landing energy with 8 nm xy resolution. Milling was performed with gallium ion column operated at 30 kV. Material was removed in 8 nm z-steps to generate a final isotropic voxel size of 8 × 8 × 8 nm. Raw data frames were converted to HDF5 and imported into the Janelia Render web service (in-house software, HHMI Janelia Research Campus https://github.com/saalfeldlab/render) for alignment with affine transformations, spectral de-streaking with spatially localized blending, and shading correction by a fourth-order model estimated from aligned tissue, followed by export to N5 for downstream analysis. Amira (Thermo Fisher Scientific) tools were used to analyze and segment the 3D volumes. Plasma membranes were segmented manually, with limited use of interpolation and smoothing. Electron-dense material near the membrane of invaginations was segmented using the magic wand. Masking limited thresholding to the structure of interest. No smoothing was applied to the resulting segmentations. Segmentation of the plasma membrane and electron-dense materials extending into the invaginations were then exported from Amira. Meshes of the MEF membrane, macrophage membrane, and electron-dense materials were extracted from the segmented ROI stack using Fiji’s 3D viewer plugin (http://fiji.sc/). The SEM volumes were converted into AVI videos. For visualization and modeling, the STL files and videos were imported to and rendered in Blender (Blender Foundation, http://www.blender.org). Using Boolean subtraction operations, we separated the electron-dense internalized material into the cytoplasmic portion and the membrane-embedded/associated portion. Small mesh fragments that appeared to be exterior to the internalized MEF membrane were excluded from the model.

### Immunofluorescence staining

Cells were fixed with 4% paraformaldehyde (PFA) in PBS for 20 min. When treated with Baf A1, cells were washed three times with DMEM before fixation. For 3D imaging, cells were stained with Alexa Fluor 488-conjugated wheat germ agglutinin (Thermo Fisher Scientific, W11261) at 5 μg/ml in HBSS without calcium, magnesium and phenol red (Thermo Fisher Scientific) supplemented with 2 mM L-glutamine for 10 min at 37°C and were washed three times with medium before fixation. Following permeabilization with 0.1% Triton X-100 and blocking with 5% bovine serum albumin (BSA), cells were stained with antibodies listed in Supplementary Table 6. Cells were mounted with ProLong Gold Antifade Mountant (Thermo Fisher Scientific) and observed with Leica Stellaris 8. Three-dimensional images were reconstructed using AIVIA (Leica) from z-stack images taken every 0.8 μm from the bottom to the top of the cells.

### tmCSF1-tethered bead uptake assay

Recombinant protein of the tmCSF1 extracellular domain (tmCSF1-ECD) with AviTag and 6 × histidine was cloned, purified and biotinylated by GenScript. The amino acid sequence is as follows: MGWSCIILFLVATATGVHSKEVSEHCSHMIGNGHLKVLQQLIDSQMETSCQIAFEFVDQEQLDDPVCYLKKAFFLVQDIIDETMRFKDNTPNANATERLQELSNNLNSCFTKDYEEQNK ACVRTFHETPLQLLEKIKNFFNETKNLLEKDWNIFTKNCNNSFAKCSSRGHVEQHEGSSDPQIPEGSGLNDIFEAQKIEWHEHHHHHH. Streptavidin-coated fluorescent beads (Spherotech, SVFP-1068-5) were washed with PBS containing 2% FBS and 1 mM EDTA (PFE) before use. The concentration of beads was determined by flow cytometric analysis. Biotinylated tmCSF1-ECD was incubated with beads at 2.2 mg/10^6^ beads in PFE for 20 min at 4°C. After washing, CTV-labelled 1.5 × 10^5^ BMDMs in 4-well chamber slides were incubated with 3 × 10^5^ tmCSF1-coated beads. Pre-treatment of BMDMs with isotype control or anti-CSF1R antibody was performed as described above. Inhibitor treatment started 30 min before incubation with tmCSF1-coated beads. Antibody and inhibitor treatment continued during culture with beads. Inhibitors and their concentrations were as follows: 1 μM wortmannin (Sigma Aldrich, 681675), 3 μM latrunculin A (Cayman Chemical, 10010630), 200 μM DMA (Cayman Chemical, 19100), 200 μM CK666 (Sigma Aldrich, SML0006), 10 μM SMIFH2 (Cayman Chemical, 31517). Positive-control experiment of DMA treatment was performed by incubating BMDMs with 100 ng/ml recombinant CSF1 (Cell Signaling Technologies, 5228) and Texas Red-labelled neutral dextran of 70,000 MW (Thermo Fisher Scientific, D1830) for 15 min in the presence of DMA After 2-hour incubation of BMDMs with beads, cells were washed three times with RPMI, fixed, mounted and observed under confocal microscope. The number of beads in each BMDM was counted using ImageJ.

### CSF1R endocytosis assay

In chamber slides, 1.5 × 10^5^ BMDMs were incubated with 50 ng/ml recombinant CSF1 for 5 min. The number of CSF1 puncta in individual BMDMs was analyzed with ImageJ. Background intensity of Alexa 488 in latrunculin A-treated BMDMs was reduced so that total Alexa488 intensity per image became similar to that of other experimental groups.

### Immunoblotting

Electrophoresis, membrane transfer and blotting of proteins in lysates were performed as described previously^75^. Blocking of membrane was performed with 5% BSA in TBST (20 mM Tris, 150 mM NaCl, 0.05% Tween 20). Antibodies used are listed in Supplementary Table 7. Blotted bands were visualized with Bio-Rad Image Lab system.

### Correlative light and electron microscopy (CLEM)

CTDR-labelled MEFs were seeded at 1.2 × 10^5^ cells on 35-mm gridded glass-bottom dish (Mattek, P35G-1.5-14-C-GRD) and cultured overnight. Following Baf A1 treatment, CTV-labelled BMDMs were added at 4 × 10^5^ cells. After 6 hours of co-culture, cells were fixed and observed under fluorescence microscope (Zeiss Axio Observer with Colibri7 light source) to determine the ROIs. Cells were further fixed with 2.5% glutaraldehyde in 0.1 M sodium cacodylate (pH = 7.4) for 2 hours and with 0.5% osmium tetroxide and 0.8% potassium ferrocyanide in 0.1M sodium cacodylate for 1 hour. Following rinse with 0.1 M sodium cacodylate and water, 2% uranyl acetate treatment was performed for 45 min. After wash with water, cells were dehydrated with 30%, 50%, 70%, 90% and 100% ethanol. The dish was incubated with 1:1 (v/v) mixture of EPON and ethanol for 20 min and subsequently with EPON for 20 min, followed by incubation at 60°C overnight for resin polymerization. Following resin separation, the ROIs were identified under the ultramicrotome, cut and mounted on dummy blocks. Sections of 70 nm were collected and treated with chloroform to stretch the cells. The samples were observed with FEI Tecnai Biotwin Transmission Electron Microscope at 80kV and AMT nanosprint 15 camera.

### Staining of protein aggregates

After fixation, cells were treated with 0.5% Triton X-100 and 3 mM EDTA in PBS for 15 min and subsequently stained with proteostat (Enzo, ENZ-51023) at 1/10,000 dilution in PBS for 30 min. Staining with thioflavin T was performed at 50 μM in PBS for 30 min. All reactions were performed at room temperature. Antibody staining was performed before proteostat staining when needed. For microscopic analysis, cells were mounted after being washed three times with PBS following proteostat staining. For flow cytometric analysis, cells were analyzed without washing after proteostat staining.

### Total internal reflection fluorescence (TIRF) microscopy

MEFs or NIH/3T3-derived transduced cells were seeded in glass-bottom 24-well plate (Sigma Aldrich, EP0030741021) and treated with Baf A1 next day. After fixation and staining with antibodies and proteostat, cells in PBS were subjected to TIRF microscopy analysis using a Nikon Ti2-E inverted motorized microscope stand with a motorized stage with a stage-top Piezo, Nikon H-TIRF, Agilent laser combiner with four lines (405, 488, 561, and 640 nm), scientific CMOS camera Photometrics Prime 95B, and 100× TIRF objective with a NA of 1.49. Images were analyzed with ImageJ.

### Immunoprecipitation

FLAG-tagged tmCSF1-transduced cells, derived from NIH/3T3, were seeded at 2.5 × 10^6^ cells in 15-cm dish and cultured overnight. After Baf A1 treatment, membrane proteins were extracted using Mem-PER Plus Membrane Protein Extraction Kit (Thermo Fisher Scientific, 89842) Centrifugation after membrane solubilization was performed at 3,000 × *g* for 10 min. Membrane protein lysates (500 μl) prepared from 5 × 10^6^ cells were mixed with 0.75 μg of control monoclonal rabbit IgG antibody (clone DA1E, Cell Signaling Technologies, 3900) or anti-FLAG antibody (clone D6W5B, Cell Signaling Technologies, 14793) overnight at 4°C, followed by incubation with 25 μl Dynabead Protein G for Immunoprecipitation (Thermo Fisher Scientific, 10003D) for 4 hours at 4°C. Beads were washed three times with IP lysis buffer (Thermo Fisher Scientific, 87787) and incubated with 1 mg/ml FLAG peptides (Thermo Fisher Scientific, A36805) in 50 μl IP lysis buffer for 30 min at room temperature. After magnetic bead separation, the eluted solution was either stained with proteostat at 1/5,000 dilution in 500 μl PBS for 15 min for flow cytometry analysis or subjected to immunoblotting.

### Mass spectrometry

Baf A1-treated MEFs were lysed with RIPA buffer (ThermoFisher Scientific, 89900) and stained with proteostat. Proteostat-positive and-negative particles were sorted into RIPA buffer using flow cytometry and collected with centrifugation at 16,000 × g for 30 min. Mass spectrometry of both particles (1.5 × 10^6^) was performed with MSB-17 (4h LC-MS/MS) in MS Bioworks. Spectral counts of identified proteins were generated after exclusion of possible contaminants and reverse hits by MS Bioworks. Normalized spectral abundance factor (NSAF) was calculated as below.

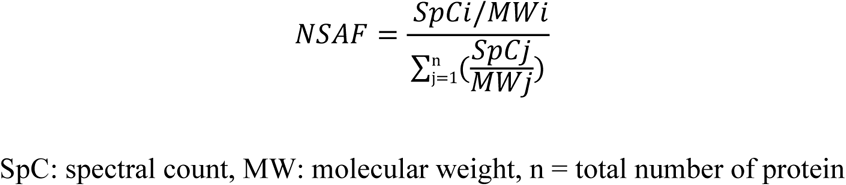

SpC: spectral count, MW: molecular weight, n = total number of protein

Enrichment of protein abundance in proteostat-positive particles was calculated by dividing NSAF of proteostat-positive particles by that of proteostat-negative particles.

## Statistical analysis

Two-tailed unpaired t test and an analysis of variance (ANOVA) test with multiple comparisons were used where appropriate to determine statistical significance between groups (*P* < 0.05 was considered significantly different) using Prism (GraphPad Software, Inc). All data are representatives of 2-3 independent experiments and shown as mean ± s.d. \**P* < 0.05, \*\**P* < 0.01, \*\*\**P* < 0.001, \*\*\*\**P* < 0.0001; ns, not significant.

## Data availability

All data are available in the main text or the supplementary materials. Code used for analyses is available at https://github.com/archavan/csf1. RNA-seq data are available in GEO with accession number GSE349575. The mass spectrometry proteomics data have been deposited to the ProteomeXchange Consortium via the PRIDE partner repository with the dataset identifier PXD085176.

## Acknowledgments

Cells and reagents were generously provided by M. Bosenberg (Yale University), P. Satpute-Krishnan (Uniformed Services University), J. Cox (University of California, Berkeley) and C. Wilen (Yale University). We thank L. Cantley (Yale University) and S. Ishibe (Yale University) for their technical advice on experiments using mouse kidney and S. Yu, C. Zhang and J. Cullen in R. Medzhitov lab for their assistance with serum collection from mice and animal care. We also thank Zachary Wright (Janelia Research Campus) for technical assistance with confocal microscopy and FIB-SEM and invaluable discussion. We would like to thank all members of the R. Medzhitov lab for their invaluable input and support throughout this work. Flow cytometry analysis was performed at Yale Flow Cytometry Facility. Time-lapse imaging was performed at Yale Stem Cell Center for some experiments. CLEM images were acquired at Center for Cellular and Molecular Imaging (CCMI) at Yale University. We thank Marley Bryant and Grace Park in Janelia’s Project Technical Resources (PTR)-BioImage Analysis team led by Andy Cole for annotation and rendering of FIB-SEM images. R. Medzhitov lab is supported by the Howard Hughes Medical Institute, the Blavatnik Family Foundation, the Food Allergy Science Initiative, and grants from the National Institutes of Health (R01AI144152, R01AI182061, P01AI179273). Y. Kimura was supported by The Uehara Memorial Foundation and Japan Society for the Promotion of Science (JSPS).

## Contributions

Conceptualization: Y.K., R.M., Methodology: Y.K., K.S., A.R.C., W.K., Z.W., M.S., W.Q., N.I., M.I., I.S., S.D.P., G.Y., A.R., C.B., E.T., K.I., X.S., J.L., R.M., Investigation: Y.K., K.S., A.R.C., W.K., Z.W., M.S., W.Q., N.I., M.I., I.S., S.D.P., A.R., Formal analysis: Y.K., K.S., A.R.C., W.K., Z.W., M.S., W.Q., N.I., M.I., I.S., S.D.P., A.R., Visualization: Y.K., K.S., A.R.C., W.K., Z.W., M.S., W.Q., N.I., M.I., I.S., S.D.P., Supervision: K.I., X.S., J.L., R.M., Resources: K.S., W.K., Z.W., M.S., W.Q., N.I., M.I., I.S., G.Y., K.I., C.B., E.T., X.S., J.L., Validation: Y.K., K.S., A.R.C., W.K., Z.W., M.S., W.Q., N.I., M.I., I.S., S.D.P., A.R., Writing—original draft: Y.K., K.S., A.R.C., W.K., I.S., S.D.P., R.M., Writing—review & editing:

## Corresponding author

Correspondence to R. Medzhitov.

---

**Extended Data Fig. 1.**
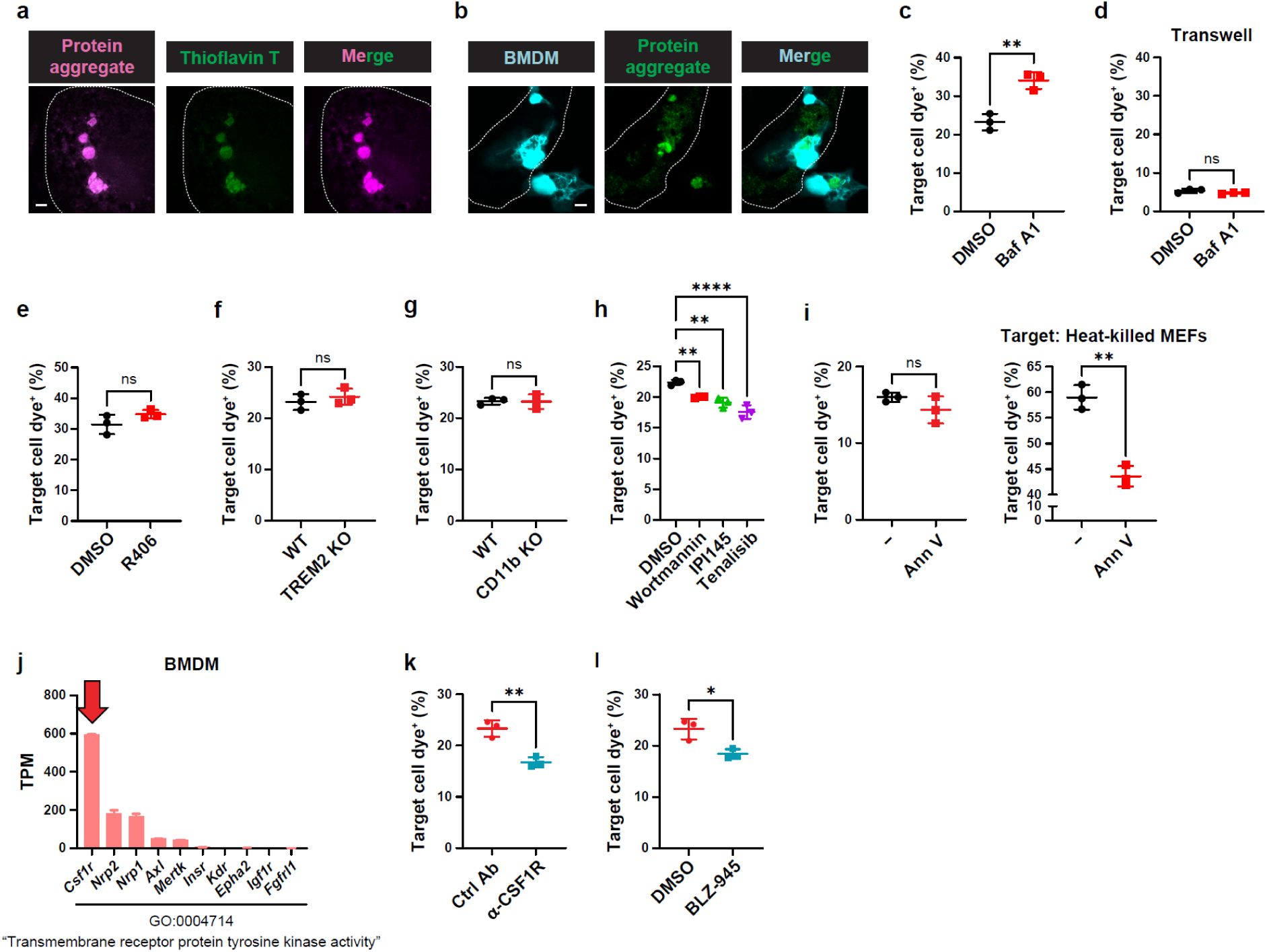
CSF1R on macrophages engages in the trogocytosis of protein aggregate-accumulating cells. a, Representative confocal images of MEF^Baf^ ^A1^. White dotted lines indicate the cell body of MEF^Baf^ ^A1^. **b,** Representative confocal images of co-cultured BMDMs and cytoplasmic labelling-free MEF^Baf^ ^A1^. White dotted lines indicate the cell body of MEF^Baf^ ^A1^. **c, d,** Uptake of materials from Baf A1-treated or untreated MEFs by BMDMs upon co-culture (**c**) or contactless co-culture in a transwell system (**d**) with each MEF. Shown are the percentages of BMDMs positive for MEF-staining membrane dye. **e-i,** Uptake of MEF^Baf^ ^A1^ materials by BMDMs under the indicated conditions. **e,** Uptake in the presence or absence of Syk inhibitor R406. **f,** Uptake by WT or TREM2 KO BMDMs. **g,** Uptake by WT or CD11b KO BMDMs. **h,** Uptake in the presence or absence of pan-PI3K inhibitor wortmannin or PI3Kγ and δ-selective inhibitors IPI145 and tenalisib. **i,** Uptake in the presence or absence of phosphatidylserine masking with annexin V. Uptake of heat-killed MEF materials in the presence or absence of annexin V is shown on the right. Shown are the percentages of BMDMs positive for MEF-staining membrane dye. **j,** Expression of receptor tyrosine kinase genes annotated to GO:0004714 (transmembrane receptor protein tyrosine kinase activity) in BMDMs, determined by RNA-seq analysis (n = 2). The 10 most highly expressed genes are shown. Expression of *Csf1r* is indicated by the red arrow. **k, l,** Uptake of tmCSF1-transduced cell materials by BMDMs in the presence of control or CSF1R-blocking antibody (**k**) and in the presence or absence of CSF1R kinase inhibitor BLZ-945 (**l**). Shown are the percentages of BMDMs positive for transduced cell-staining cytoplasmic dye. Graphs show mean ± s.d. **c-i, k, l,** n = 3 per group. **c-g, i, k, l,** Two-tailed unpaired Student’s t test. **h,** One-way ANOVA with Dunnett’s multiple-comparison test. \**P* < 0.05, \*\**P* < 0.01, \*\*\*\**P* < 0.0001; ns, not significant.

**Extended Data Fig. 2.**
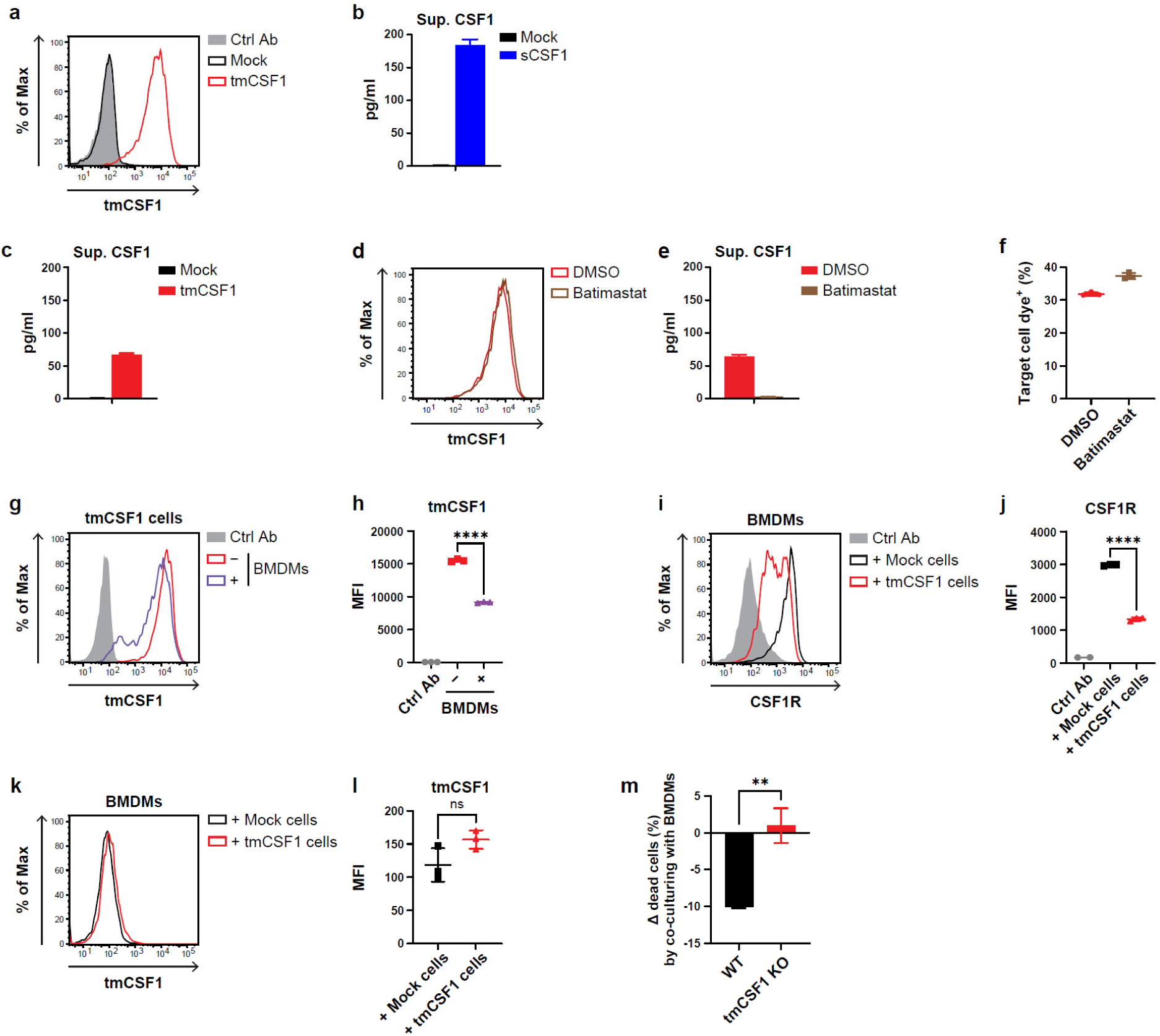
tmCSF1, but not cleaved tmCSF1, induces trogocytosis and enhances the fitness of protein aggregate-harboring cells. a, Representative histograms of tmCSF1 protein expression on mock-and tmCSF1-transduced cells. **b, c,** CSF1 protein level in the culture supernatant of mock-and sCSF1-transduced cells (**b**) and mock-and tmCSF1-transduced cells (**c**). **d, e,** Representative histograms of tmCSF1 protein expression (**d**) and CSF1 protein level in the culture supernatant (**e**) of tmCSF1-transduced cells treated with or without ADAM17 inhibitor batimastat. **f,** Uptake of tmCSF1-transduced cell materials by BMDMs in the presence or absence of batimastat. Shown are the percentages of BMDMs positive for transduced cell-staining cytoplasmic. **g, h,** Representative histograms (**g**) and quantification (**h**) of tmCSF1 expression on tmCSF1-transduced cells co-cultured with or without BMDMs. **i, j,** Representative histograms (**i**) and quantification (**j**) of CSF1R expression on BMDMs co-cultured with mock-or tmCSF1-transduced cells. **k, l,** Representative histograms (**k**) and quantification (**l**) of tmCSF1 expression on BMDMs co-cultured with mock-or tmCSF1-transduced cells. **m,** Changes in the proportion of dead cells among WT and tmCSF1 KO MEF^Baf^ ^A1^ sorted after co-culture with BMDMs and subsequently incubated, compared with cells cultured alone. **b, c, e, f, h, j, l, m,** n = 3 per group. Graphs show mean ± s.d. **h, j, l, m,** Two-tailed unpaired Student’s t test. \*\**P* < 0.01, \*\*\*\**P* < 0.0001; ns, not significant.

**Extended Data Fig. 3.**
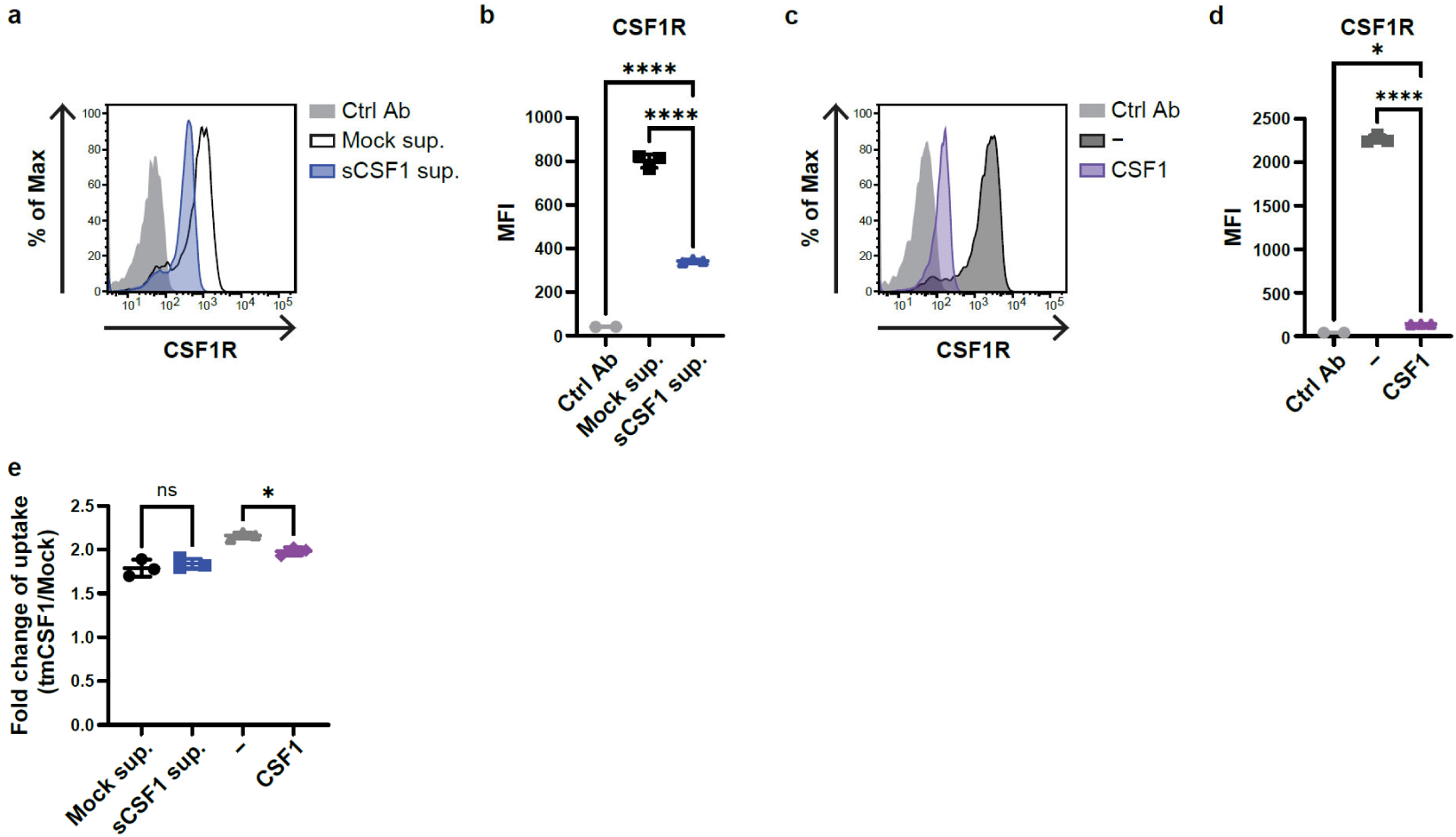
sCSF1 minimally blocks tmCSF1-mediated trogocytosis. a, b, Representative histograms (**a**) and quantification (**b**) of CSF1R expression on BMDMs treated with the culture supernatant from mock-or sCSF1-transduced cells. **c, d,** Representative histograms (**c**) and quantification (**d**) of CSF1R expression on BMDMs treated with or without recombinant CSF1 protein. **e,** Fold change in uptake of materials from tmCSF1-transduced cells compared with mock-transduced cells by BMDMs in the presence of the culture supernatant from mock-or sCSF1-transduced cells, with or without recombinant CSF1 treatment. The percentage of BMDMs positive for tmCSF1-transduced cell-labelling dye was divided by the percentage of BMDMs positive for mock-transduced cell-labelling dye in each condition. **b, d, e,** n =3 per group. Graphs show mean ± s.d. **b, d,** One-way ANOVA with Dunnett’s multiple-comparison test. **e,** Two-way ANOVA with Sidak’s multiple-comparison test. \**P* < 0.05, \*\*\*\**P* < 0.0001; ns, not significant.

**Extended Data Fig. 4.**
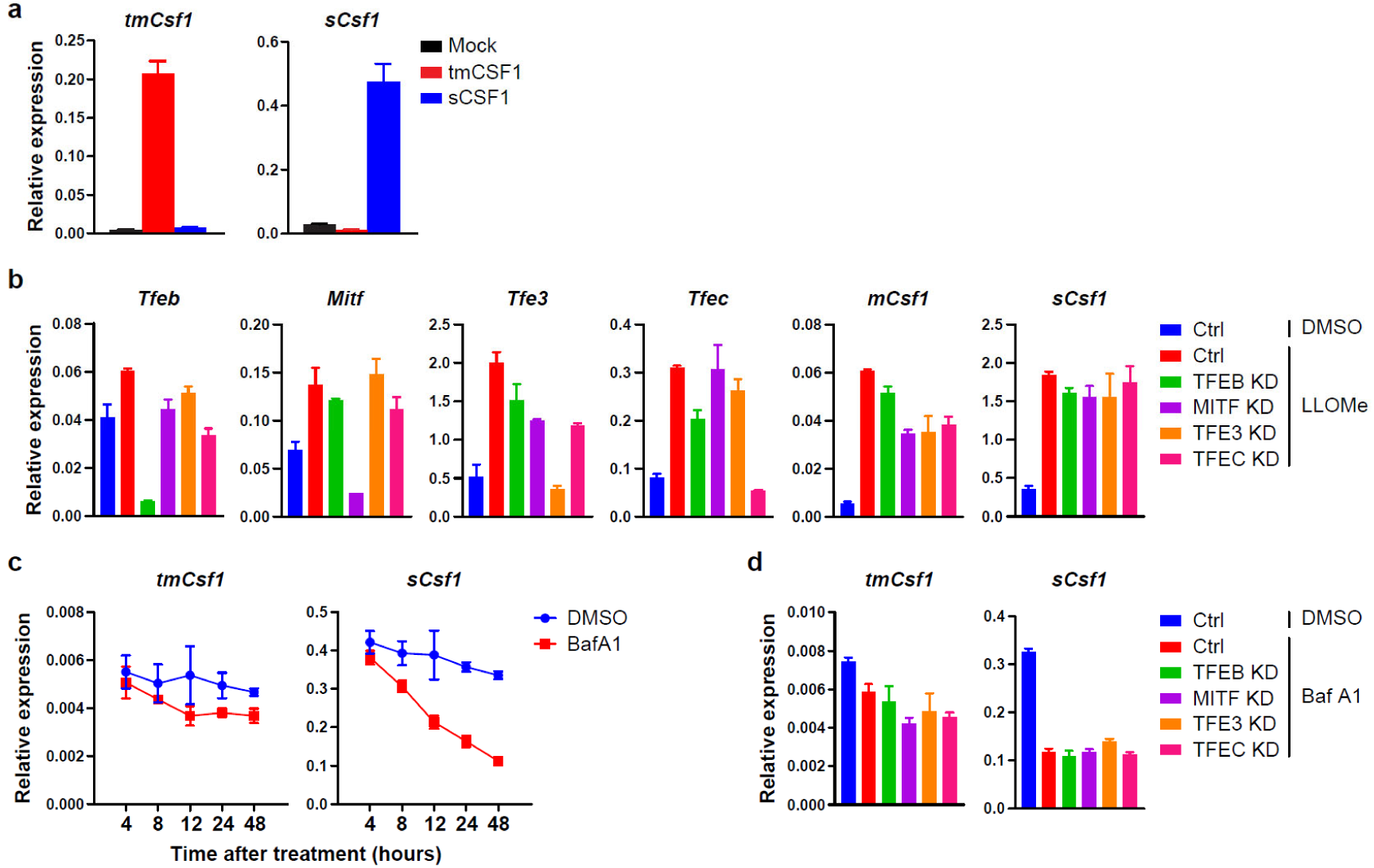
Lysosomal stress responses skew the balance of CSF1 isoforms toward tmCSF1. a, Specificity of quantitative polymerase chain reaction (qPCR) primers for *tmCsf1* and *sCsf1* detection. Shown are *tmCsf1* and *sCsf1* expression levels in mock-, tmCSF1-and sCSF1-transduced cells measured using the designed *tmCsf1* and *sCsf1* primers. **b,** Expression of the indicated genes in control (Ctrl), TFEB knockdown (KD), MITF KD TFE3 KD or TFEC KD MEFs treated with or without LLOMe. **c,** *tmCsf1* and *sCsf1* expression in MEFs treated with or without Baf A1. **d,** *tmCsf1* and *sCsf1* expression in Ctrl, TFEB KD, MITF KD TFE3 KD or TFEC KD MEFs treated with or without Baf A1. n = 3 per group. Graphs show mean ± SD.

**Extended Data Fig. 5.**
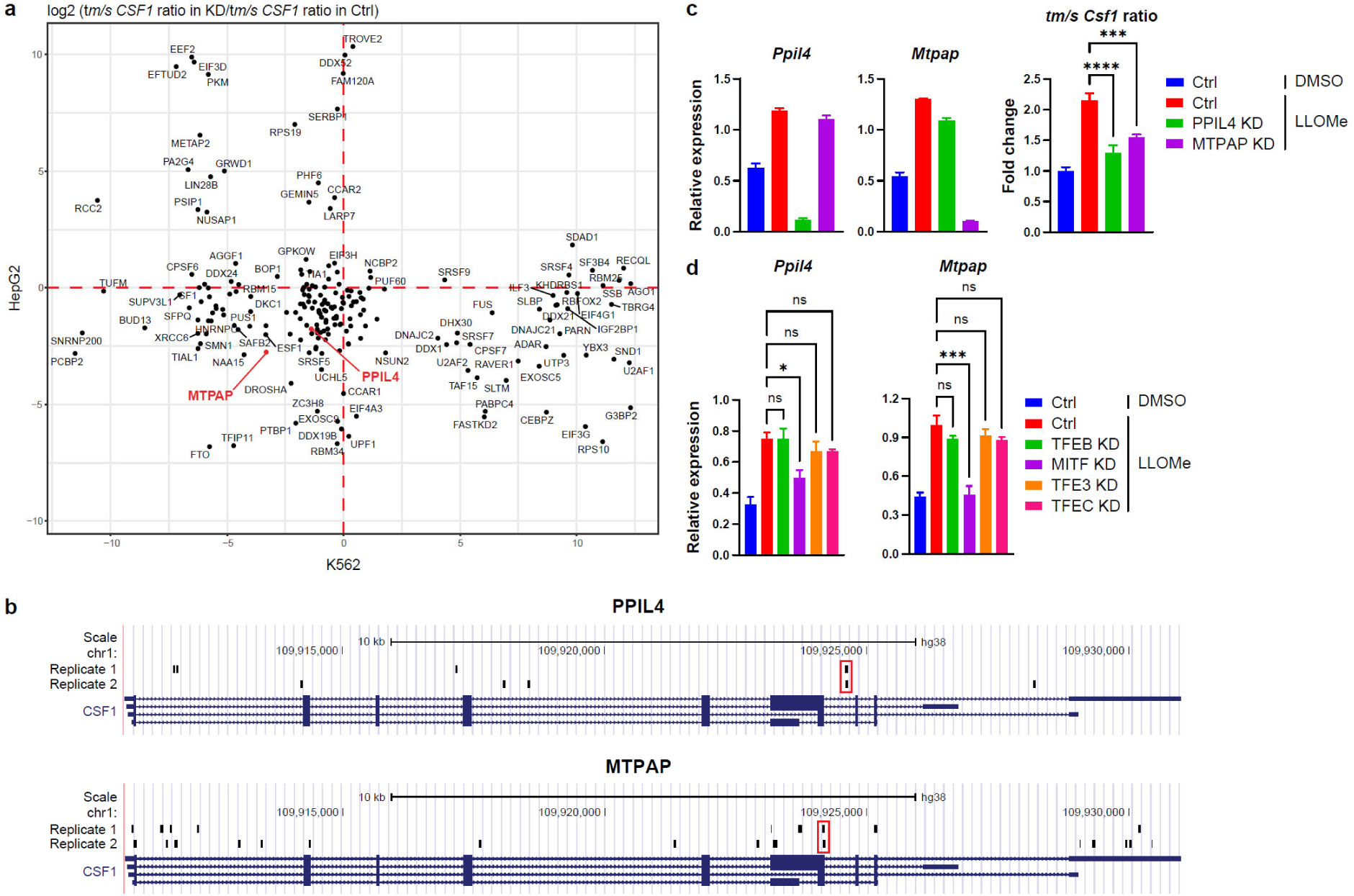
PPIL4 and MTPAP bind to CSF1 RNA and promote tmCSF1-skewing. a, RNA-binding proteins (RBPs) that control the expression of *CSF1* isoforms. Plots show the fold change of *tmCSF1* to *sCSF1* expression ratio following KD of the indicated RBPs in HepG2 (x-axis) and K562 (y-axis) cells. Data are from RNA-seq analysis reported by Van Nostrand *et al*^34^. **b,** Binding sites of PPIL4 and MTPAP on *CSF1* pre-mRNA identified by eCLIP. Black-filled boxes indicate all detected binding sites, and red open boxes indicate sites detected in both biological replicates. Data are from Van Nostrand *et al*^34^. **c,** Expression of the indicated genes in Ctrl, PPIL4 KD or MTPAP KD MEFs treated with or without LLOMe. **d, h,** *Ppil4* and *Mtpap* expression in Ctrl, TFEB KD, MITF KD TFE3 KD or TFEC KD MEFs treated with or without LLOMe.. **c, d,** n = 3 per group. Graphs show mean ± SD. **c, d,** One-way ANOVA with Dunnett’s multiple-comparison test. \**P* < 0.05, \*\*\**P* < 0.001, \*\*\*\**P* < 0.0001; ns, not significant.

**Extended Data Fig. 6.**
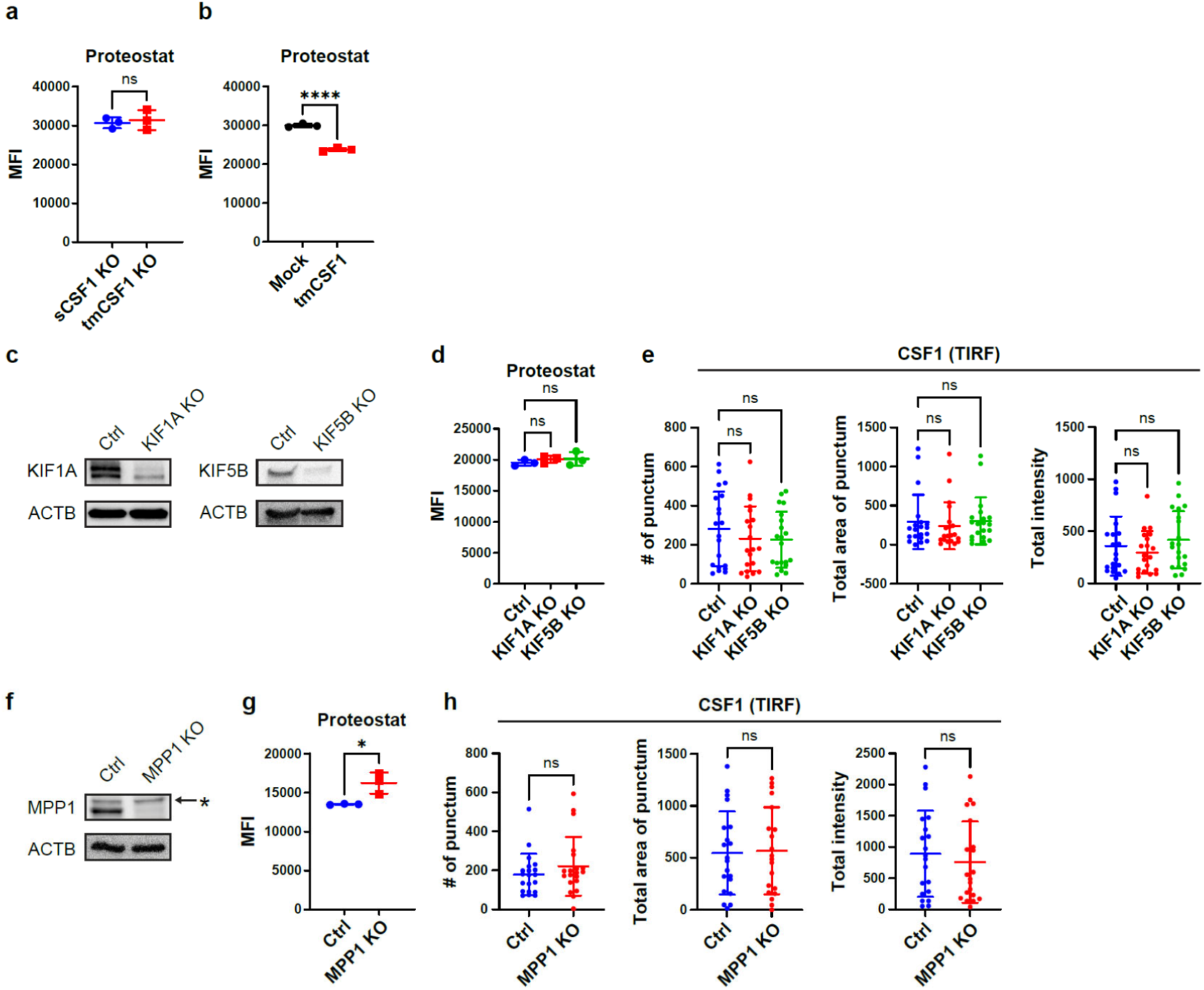
Total amount of cellular protein aggregates and surface tmCSF1 expression are not enhanced by the program to associate protein aggregate-laden lysosomes with tmCSF1. a, b, Total amount of protein aggregates in sCSF1 KO and tmCSF1 KO MEF^Baf^ ^A1^ (**a**) and Baf A1-treated mock-and tmCSF1-transduced cells (**b**). **c,** Representative confocal images of MEFs treated with Baf A1 for the indicated times. Protein aggregates are zoomed in. **c,** Immunoblot images of Ctrl, KIF1A KO and KIF5B KO MEFs. **d,** Total amount of protein aggregates in Ctrl, KIF1A KO and KIF5B KO MEF^Baf^ ^A1^. **e,** Number, total area and total intensity of CSF1 puncta in TIRF microscopic images of Ctrl, KIF1A KO and KIF5B KO MEF^Baf^ ^A1^. **f,** Immunoblot images of Ctrl and MPP1 KO MEFs. Asterisk indicates non-specifically detected bands. **g,** Total amount of protein aggregates in Ctrl and MPP1 KO sCSF1-deficient MEF^Baf^ ^A1^. **h,** Number, total area and total intensity of CSF1 puncta in TIRF microscopic images of Ctrl and MPP1 KO sCSF1-deficient MEF^Baf^ ^A1^. **a, b, d, g,** n = 3 per group. **e, h,** n = 20 per group. Graphs show mean ± SD. **a, b, g, h,** Two-tailed unpaired Student’s t test. **d, e,** One-way ANOVA with Dunnett’s multiple-comparison test. \**P* < 0.05, \*\*\*\**P* < 0.0001; ns, not significant.

**Extended Data Fig. 7.**
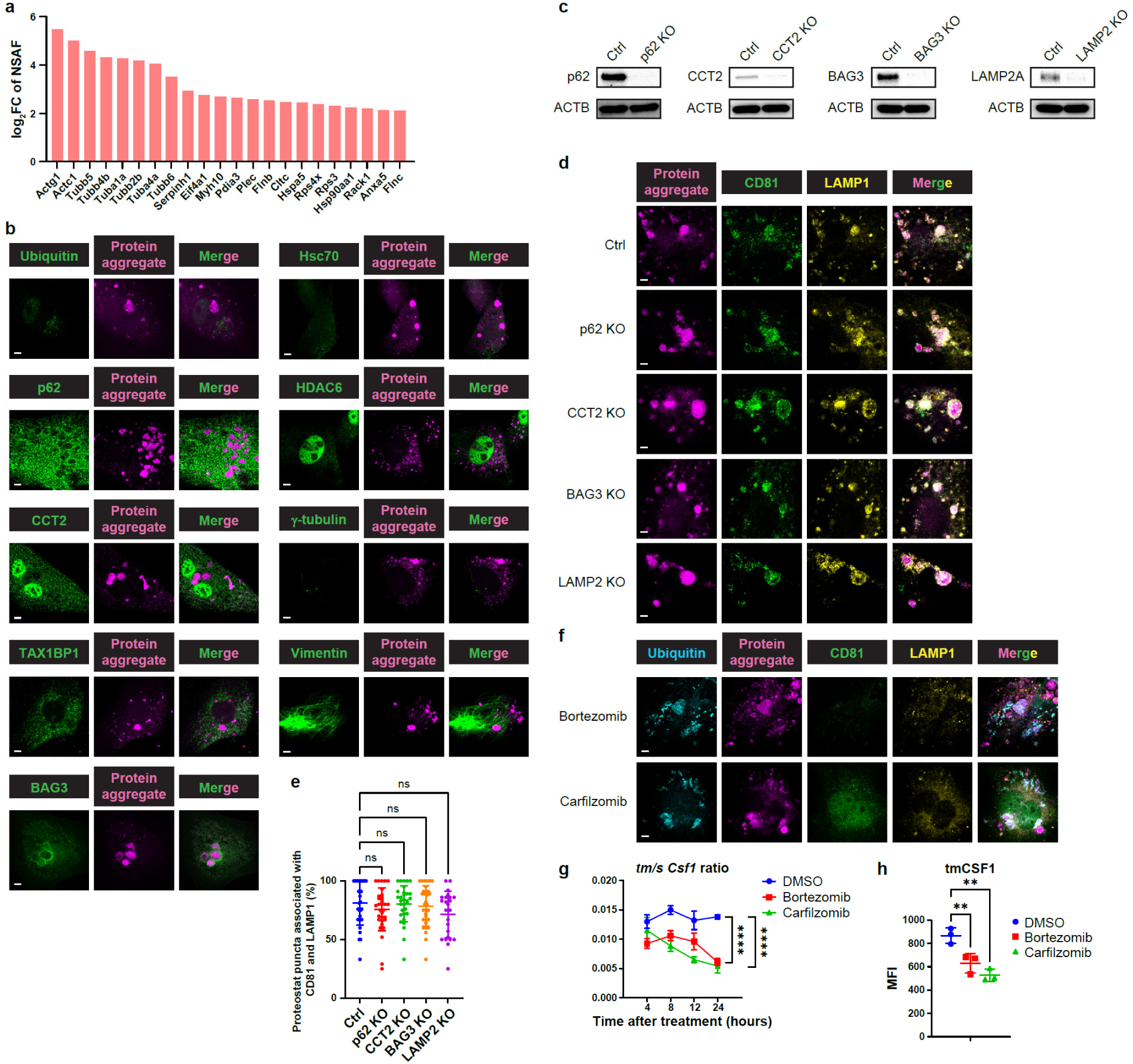
Trogocytosis target aggregates are compartmentalized into lysosomal vesicles independently of canonical protein homeostasis programs. a, Enrichment of protein abundance in proteostat-positive insoluble particles relative to proteostat-negative insoluble particles sorted from MEF^Baf^ ^A1^ lysate. More than 4-fold-enriched proteins are shown. **b,** Representative confocal images of MEF^Baf^ ^A1^. Protein aggregates are zoomed in. **c,** Immunoblot images of Ctrl, p62 KO, CCT2 KO, BAG3 KO MEFs and LAMP2 KO MEFs. **d, e,** Representative confocal images of (**d**) and percentages of protein aggregate puncta associated with both CD81 and LAMP1 in (**e**) Ctrl, p62 KO, CCT2 KO, BAG3 KO and LAMP2 KO MEF^Baf^ ^A1^. Zoomed protein aggregates are shown in (**d**). **f,** Representative confocal images of MEFs treated with proteasome inhibitors bortezomib or carfilzomib. Protein aggregates are zoomed in. **g, h,** *tmCsf1* to *sCsf1* expression ratio in (**g**) and tmCSF1 protein expression on (**h**) MEFs treated with DMSO, bortezomib or carfilzomib. Scale bars, 5 μm. Graphs show mean ± s.d. **e, g, h,** One-way ANOVA with Dunnett’s multiple-comparison test. \*\**P* < 0.01, \*\*\*\**P* < 0.0001; ns, not significant.

**Extended Data Fig. 8.**
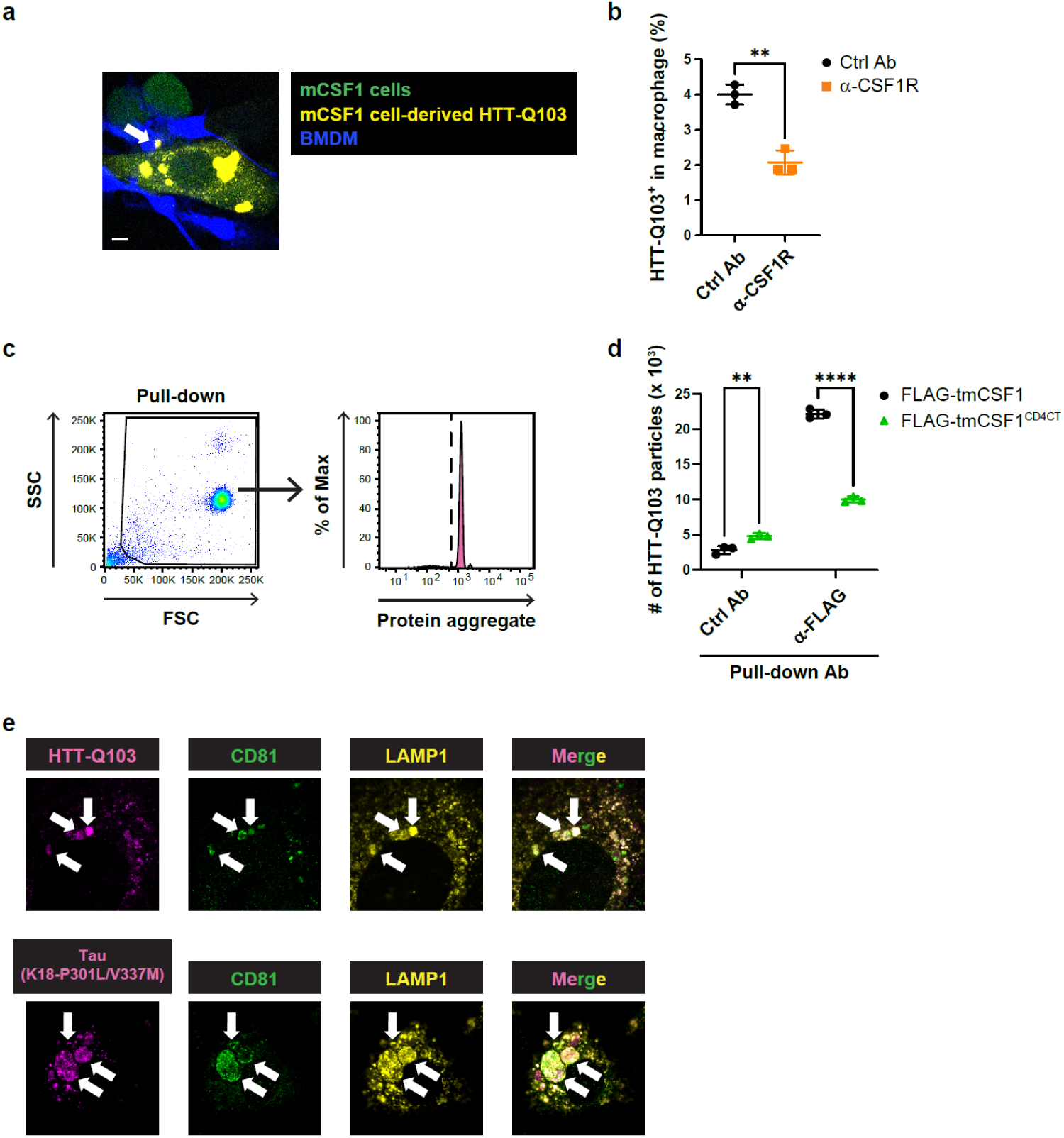
Aggregates of pathogenic model proteins are incorporated by macrophages via tmCSF1-dependent trogocytic program. a, Representative confocal images of co-cultured BMDMs and tmCSF1-transduced cells overexpressing mCherry-conjugated HTT-Q103. GFP is expressed by tmCSF1-transduction vector. White arrows indicate HTT-Q103 incorporated into BMDMs. **b,** Uptake of HTT-Q103 from tmCSF1-transduced cells by BMDMs in the presence of control or CSF1R-blocking antibody. Shown are percentages of BMDMs positive for mCherry-conjugated HTT-Q103. **c, d,** mCherry-positive protein aggregate particles pulled down with control or anti-FLAG antibody from plasma membrane lysates of FLAG-tmCSF1-or FLAG-tmCSF1^CD4CT^-transduced cells that overexpress mCherry-conjugated HTT-Q103. Shown are flow-cytometric gating strategy to detect the particles (**c**) and quantification of mCherry-positive particle numbers (**d**)**. e,** Representative confocal images of MEFs transduced with mCherry-conjugated HTT-Q103 or mScarletI-conjugated Tau.K18(P301L/V337M). **b, d,** n = 3 per group. Scale bars, 10 μm. Graphs show mean ± s.d. **b,** Two-tailed unpaired Student’s t test. **d,** Two-way ANOVA with Sidak’s multiple-comparison test. \*\**P* < 0.01, \*\*\*\**P* < 0.0001.

**Extended Data Fig. 9.**
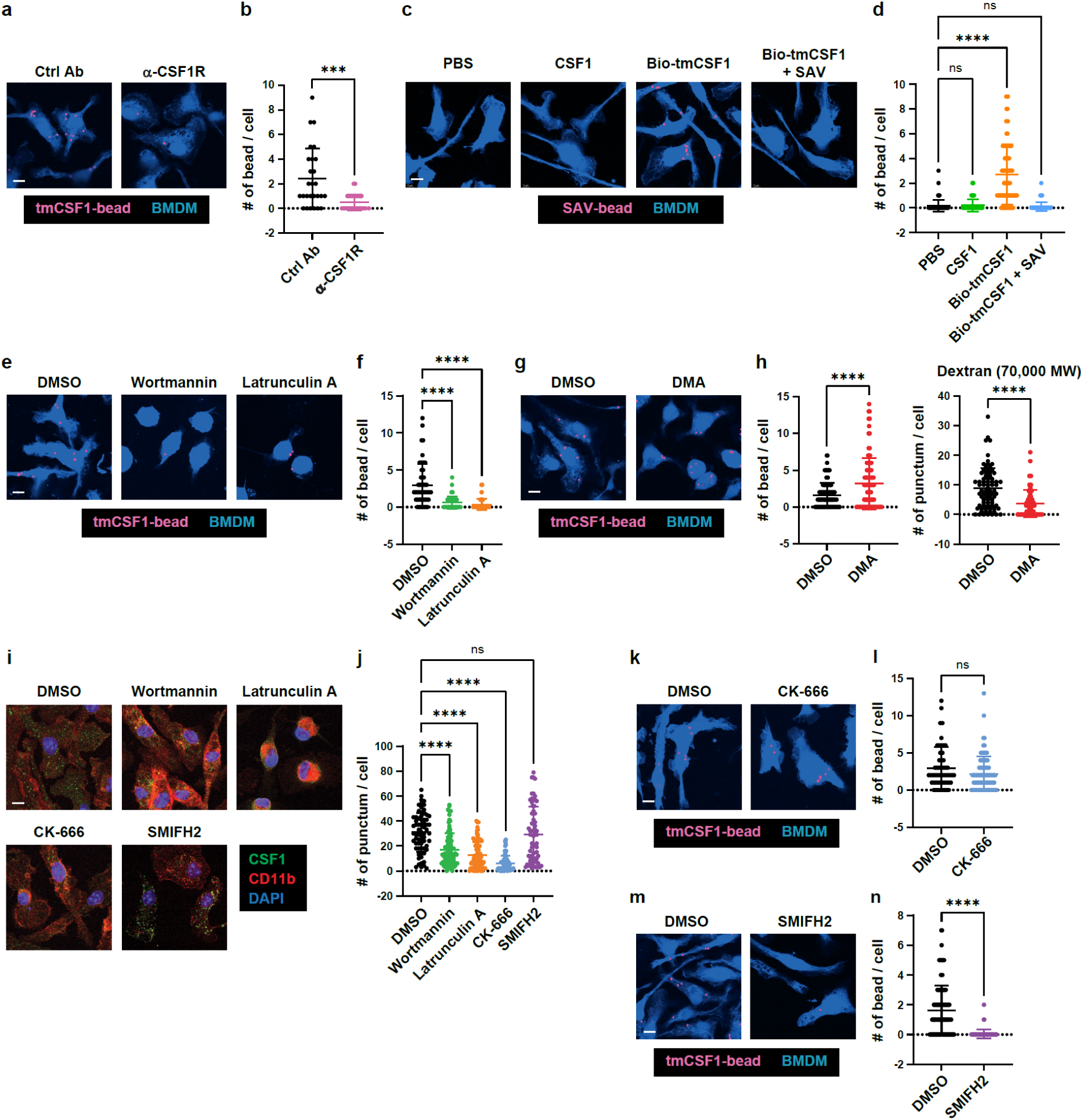
CSF1R exhibits phagocytic activity on tmCSF1-tethered particles. a, b, Uptake of tmCSF1-tethered beads by BMDMs in the presence of control or CSF1R-blocking antibody. Shown are representative confocal images (**a**) and numbers (**b**) of tmCSF1-tethered beads in each BMDM (n =27 Ctrl Ab, 29 α-CSF1R). **c, d,** Uptake of streptavidin (SAV)-coated beads by BMDMs upon the treatment with PBS, recombinant CSF1, biotinylated tmCSF1 (Bio-tmCSF1) or Bio-tmCSF1 + SAV. Shown are representative confocal images (**c**) and numbers (**d**) of tmCSF1-tethered beads in each BMDM (n =84 PBS, 77 CSF1, 87 Bio-tmCSF1, 85 Bio-tmCSF1 + SAV). **e, f,** Uptake of tmCSF1-tethered beads by BMDMs treated with DMSO, wortmannin or latrunculin A. Shown are representative confocal images (**e**) and numbers (**f**) of tmCSF1-tethered beads in each BMDM (n =53 DMSO, 76 Wortmannin, 63 Latrunculin A). **g, h,** Uptake of tmCSF1-tethered beads or dextran by BMDMs treated with DMSO or DMA. Shown are representative confocal images (**g**), numbers (**h,** *left*) of tmCSF1-tethered beads in each BMDM (n =115 DMSO, 94 DMA) and numbers of dextran in each BMDM (**h,** *right*) (n =77 DMSO, 57 DMA). **i, j,** Uptake of recombinant CSF1 by BMDMs treated with DMSO, wortmannin, latrunculin A, CK-666 or SMIFH2. Shown are representative confocal images (**i**) and numbers (**j**) of CSF1 puncta in each BMDM (n =67 DMSO, 88 Wortmannin, 85 Latrunculin A, 68 CK-666, 65 SMIFH2). **k, l,** Uptake of tmCSF1-tethered beads by BMDMs treated with DMSO or CK-666. Shown are representative confocal images (**k**) and numbers (**l**) of tmCSF1-tethered beads in each BMDM (n =53 DMSO, 88 CK-666). **m, n,** Uptake of tmCSF1-tethered beads by BMDMs treated with DMSO or SMIFH2. Shown are representative confocal images (**m**) and numbers (**n**) of tmCSF1-tethered beads in each BMDM (n =115 DMSO, 66 SMIFH2). **a, c, e, g, i, k, m,** Representative confocal images. **b, d, f, h, j, l, n,** Numbers of the beads (**b, d, f, h, l, n**) and CSF1 puncta (**j**) in individual cells. Scale bar, 5 μm. Graphs show mean ± s.d. **b, h, i, n,** Two-tailed unpaired Student’s t test. **d, f, j,** One-way ANOVA with Dunnett’s multiple-comparison test. \*\*\**P* < 0.001, \*\*\*\**P* < 0.0001; ns, not significant.

**Extended Data Fig. 10.**
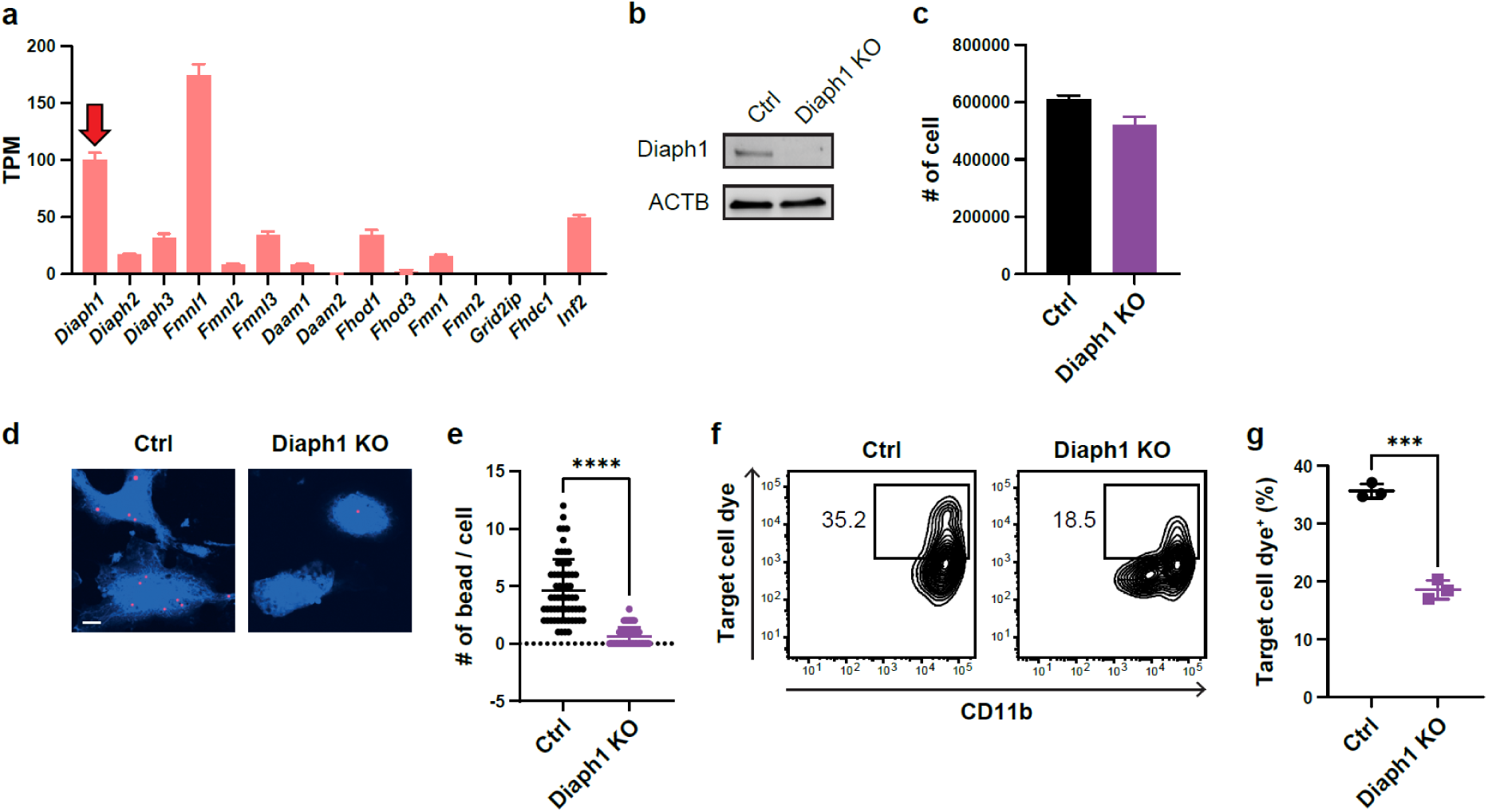
Diaph1 is required for CSF1R-mediated uptake of target materials. a, Expression of formin family member genes in BMDMs, analyzed by RNA-seq analysis (n=2). Expression of *Diaph1* is indicated by the red arrow. **b,** Immunoblot images of Ctrl and Diaph1 KO macrophages derived from ER-Hoxb8 cells. **c,** Numbers of Ctrl and Diaph1 KO ER-Hoxb8 cells upon differentiation to macrophages (n = 3 per group). **d, e,** Uptake of tmCSF1-tethered beads by Ctrl and Diaph1 KO ER-Hoxb8-derived macrophages. Shown are representative confocal images (**d**) and numbers (**e**) of tmCSF1-tethered beads in each macrophage (n =63 Ctrl, 57 Diaph1 KO). Scale bar, 5 μm. **f, g,** Uptake of MEF^Baf^ ^A1^ materials by Ctrl and Diaph1 KO macrophages derived from ER-Hoxb8 cells. Shown are the representative contour plots (**f**) and percentages (**g**) of macrophages positive for MEF-staining membrane dye (n = 3 per group). Graphs show mean ± s.d. **e, f,** Two-tailed unpaired Student’s t test. \*\*\**P* < 0.001, \*\*\*\**P* < 0.0001.

## References

1 Gonzalez-Garcia, M., Fusco, G. & De Simone, A. Membrane Interactions and Toxicity by Misfolded Protein Oligomers. Front Cell Dev Biol 9, 642623 (2021). 10.3389/fcell.2021.642623

2 Ross, C. A. & Poirier, M. A. Opinion: What is the role of protein aggregation in neurodegeneration? Nat Rev Mol Cell Biol 6, 891–898 (2005). 10.1038/nrm1742

3 Henning, R. H. & Brundel, B. Proteostasis in cardiac health and disease. Nat Rev Cardiol 14, 637–653 (2017). 10.1038/nrcardio.2017.89

4 Lopez-Otin, C., Blasco, M. A., Partridge, L., Serrano, M. & Kroemer, G. Hallmarks of aging: An expanding universe. Cell 186, 243–278 (2023). 10.1016/j.cell.2022.11.001

5 Hartl, F. U., Bracher, A. & Hayer-Hartl, M. Molecular chaperones in protein folding and proteostasis. Nature 475, 324–332 (2011). 10.1038/nature10317

6 Pohl, C. & Dikic, I. Cellular quality control by the ubiquitin-proteasome system and autophagy. Science 366, 818–822 (2019). 10.1126/science.aax3769

7 Kaushik, S. & Cuervo, A. M. The coming of age of chaperone-mediated autophagy. Nat Rev Mol Cell Biol 19, 365–381 (2018). 10.1038/s41580-018-0001-6

8 Mizushima, N., Yoshimori, T. & Ohsumi, Y. The role of Atg proteins in autophagosome formation. Annu Rev Cell Dev Biol 27, 107–132 (2011). 10.1146/annurev-cellbio-092910-154005

9 Ye, Y. Regulation of protein homeostasis by unconventional protein secretion in mammalian cells. Semin Cell Dev Biol 83, 29–35 (2018). 10.1016/j.semcdb.2018.03.006

10 Mackay, C. R. Chemokines: immunology’s high impact factors. Nat Immunol 2, 95–101 (2001). 10.1038/84298

11 Mookherjee, N., Anderson, M. A., Haagsman, H. P. & Davidson, D. J. Antimicrobial host defence peptides: functions and clinical potential. Nat Rev Drug Discov 19, 311–332 (2020). 10.1038/s41573-019-0058-8

12 Semenza, G. L. Regulation of oxygen homeostasis by hypoxia-inducible factor 1. Physiology (Bethesda) 24, 97–106 (2009). 10.1152/physiol.00045.2008

13 Perez-Gutierrez, L. & Ferrara, N. Biology and therapeutic targeting of vascular endothelial growth factor A. Nat Rev Mol Cell Biol 24, 816–834 (2023). 10.1038/s41580-023-00631-w

14 Park, M. D., Silvin, A., Ginhoux, F. & Merad, M. Macrophages in health and disease. Cell 185, 4259–4279 (2022). 10.1016/j.cell.2022.10.007

15 Brown, T. Observations by immunofluorescence microscopy and electron microscopy on the cytopathogenicity of Naegleria fowleri in mouse embryo-cell cultures. J Med Microbiol 12, 363–371 (1979). 10.1099/00222615-12-3-363

16 Joly, E. & Hudrisier, D. What is trogocytosis and what is its purpose? Nat Immunol 4, 815 (2003). 10.1038/ni0903-815

17 Faust, T. E., Gunner, G. & Schafer, D. P. Mechanisms governing activity-dependent synaptic pruning in the developing mammalian CNS. Nat Rev Neurosci 22, 657–673 (2021). 10.1038/s41583-021-00507-y

18 Velmurugan, R., Challa, D. K., Ram, S., Ober, R. J. & Ward, E. S. Macrophage-Mediated Trogocytosis Leads to Death of Antibody-Opsonized Tumor Cells. Mol Cancer Ther 15, 1879–1889 (2016). 10.1158/1535-7163.MCT-15-0335

19 Matlung, H. L. et al. Neutrophils Kill Antibody-Opsonized Cancer Cells by Trogoptosis. Cell Rep 23, 3946–3959 e3946 (2018). 10.1016/j.celrep.2018.05.082

20 Fan, A. C. et al. Submicrometre sampling of living cells by macrophages. Nature 654, 495–503 (2026). 10.1038/s41586-026-10435-5

21 Mattei, F. et al. Trogocytosis in innate immunity to cancer is an intimate relationship with unexpected outcomes. iScience 25, 105110 (2022). 10.1016/j.isci.2022.105110

22 Underhill, D. M. & Goodridge, H. S. Information processing during phagocytosis. Nat Rev Immunol 12, 492–502 (2012). 10.1038/nri3244

23 Arandjelovic, S. & Ravichandran, K. S. Phagocytosis of apoptotic cells in homeostasis. Nat Immunol 16, 907–917 (2015). 10.1038/ni.3253

24 Rothlin, C. V., Carrera-Silva, E. A., Bosurgi, L. & Ghosh, S. TAM receptor signaling in immune homeostasis. Annu Rev Immunol 33, 355–391 (2015). 10.1146/annurev-immunol-032414-112103

25 Sehgal, A., Irvine, K. M. & Hume, D. A. Functions of macrophage colony-stimulating factor (CSF1) in development, homeostasis, and tissue repair. Semin Immunol 54, 101509 (2021). 10.1016/j.smim.2021.101509

26 Pixley, F. J. & Stanley, E. R. CSF-1 regulation of the wandering macrophage: complexity in action. Trends Cell Biol 14, 628–638 (2004). 10.1016/j.tcb.2004.09.016

27 Horiuchi, K. et al. Cell surface colony-stimulating factor 1 can be cleaved by TNF-alpha converting enzyme or endocytosed in a clathrin-dependent manner. J Immunol 179, 6715–6724 (2007). 10.4049/jimmunol.179.10.6715

28 Tekguc, M., Wing, J. B., Osaki, M., Long, J. & Sakaguchi, S. Treg-expressed CTLA-4 depletes CD80/CD86 by trogocytosis, releasing free PD-L1 on antigen-presenting cells. Proc Natl Acad Sci U S A 118 (2021). 10.1073/pnas.2023739118

29 Schriek, P. et al. Marginal zone B cells acquire dendritic cell functions by trogocytosis. Science 375, eabf7470 (2022). 10.1126/science.abf7470

30 Keane, C. et al. Macrophages tune NK cell IFNgamma production through direct lipid transfer. Nat Commun 17 (2026). 10.1038/s41467-026-76444-0

31 Yao, G. Q. et al. Selective deletion of the membrane-bound colony stimulating factor 1 isoform leads to high bone mass but does not protect against estrogen-deficiency bone loss. J Bone Miner Metab 30, 408–418 (2012). 10.1007/s00774-011-0336-y

32 Stanley, E. R. & Chitu, V. CSF-1 receptor signaling in myeloid cells. Cold Spring Harb Perspect Biol 6 (2014). 10.1101/cshperspect.a021857

33 Perera, R. M., Di Malta, C. & Ballabio, A. MiT/TFE Family of Transcription Factors, Lysosomes, and Cancer. Annu Rev Cancer Biol 3, 203–222 (2019). 10.1146/annurev-cancerbio-030518-055835

34 Van Nostrand, E. L. et al. A large-scale binding and functional map of human RNA-binding proteins. Nature 583, 711–719 (2020). 10.1038/s41586-020-2077-3

35 Yao, G. Q., Troiano, N., Simpson, C. A. & Insogna, K. L. Selective deletion of the soluble Colony-Stimulating Factor 1 isoform in vivo prevents estrogen-deficiency bone loss in mice. Bone Res 5, 17022 (2017). 10.1038/boneres.2017.22

36 Nakatogawa, H. Mechanisms governing autophagosome biogenesis. Nat Rev Mol Cell Biol 21, 439–458 (2020). 10.1038/s41580-020-0241-0

37 Ganesan, D. & Cai, Q. Understanding amphisomes. Biochem J 478, 1959–1976 (2021). 10.1042/BCJ20200917

38 Marks, M. S., Heijnen, H. F. & Raposo, G. Lysosome-related organelles: unusual compartments become mainstream. Curr Opin Cell Biol 25, 495–505 (2013). 10.1016/j.ceb.2013.04.008

39 Banushi, B. & Simpson, F. Overlapping Machinery in Lysosome-Related Organelle Trafficking: A Lesson from Rare Multisystem Disorders. Cells 11 (2022). 10.3390/cells11223702

40 Bento-Lopes, L. et al. Melanin’s Journey from Melanocytes to Keratinocytes: Uncovering the Molecular Mechanisms of Melanin Transfer and Processing. Int J Mol Sci 24 (2023). 10.3390/ijms241411289

41 De Pace, R., Ghosh, S., Williamson, C. D. & Bonifacino, J. S. BLOC-1 and BORC: Complex regulators of endolysosomal dynamics. Cell Chem Biol 32, 1106–1124 (2025). 10.1016/j.chembiol.2025.08.001

42 Guardia, C. M., Farias, G. G., Jia, R., Pu, J. & Bonifacino, J. S. BORC Functions Upstream of Kinesins 1 and 3 to Coordinate Regional Movement of Lysosomes along Different Microtubule Tracks. Cell Rep 17, 1950–1961 (2016). 10.1016/j.celrep.2016.10.062

43 Jia, J. Global cellular responses to lysosomal damage. Curr Biol 36, R807–R820 (2026). 10.1016/j.cub.2026.05.050

44 Gahlot, P. et al. Lysosomal damage sensing and lysophagy initiation by SPG20-ITCH. Mol Cell 84, 1556–1569 e1510 (2024). 10.1016/j.molcel.2024.02.029

45 Goul, C. S. et al. LASER couples damage sensing to ESCRT assembly for lysosome repair. Nature 656, 216–226 (2026). 10.1038/s41586-026-10604-6

46 Stucchi, R. et al. Regulation of KIF1A-Driven Dense Core Vesicle Transport: Ca(2+)/CaM Controls DCV Binding and Liprin-alpha/TANC2 Recruits DCVs to Postsynaptic Sites. Cell Rep 24, 685–700 (2018). 10.1016/j.celrep.2018.06.071

47 Zhu, J., Shang, Y. & Zhang, M. Mechanistic basis of MAGUK-organized complexes in synaptic development and signalling. Nat Rev Neurosci 17, 209–223 (2016). 10.1038/nrn.2016.18

48 Chen, Y. R. et al. Tissue-specific landscape of protein aggregation and quality control in an aging vertebrate. Dev Cell 59, 1892–1911 e1813 (2024). 10.1016/j.devcel.2024.04.014

49 Iqbal, K., Liu, F. & Gong, C. X. Tau and neurodegenerative disease: the story so far. Nat Rev Neurol 12, 15–27 (2016). 10.1038/nrneurol.2015.225

50 Furst, D. O. et al. Filamin C-related myopathies: pathology and mechanisms. Acta Neuropathol 125, 33–46 (2013). 10.1007/s00401-012-1054-9

51 McLendon, P. M. & Robbins, J. Desmin-related cardiomyopathy: an unfolding story. Am J Physiol Heart Circ Physiol 301, H1220–1228 (2011). 10.1152/ajpheart.00601.2011

52 Kotaich, F., Caillol, D. & Bomont, P. Neurofilaments in health and Charcot-Marie-Tooth disease. Front Cell Dev Biol 11, 1275155 (2023). 10.3389/fcell.2023.1275155

53 Mauthe, M., Kampinga, H. H., Hipp, M. S. & Reggiori, F. Digest it all: the lysosomal turnover of cytoplasmic aggregates. Trends Biochem Sci 48, 216–228 (2023). 10.1016/j.tibs.2022.09.012

54 Johnston, H. E. & Samant, R. S. Alternative systems for misfolded protein clearance: life beyond the proteasome. FEBS J 288, 4464–4487 (2021). 10.1111/febs.15617

55 Chen, J. J. et al. Compromised function of the ESCRT pathway promotes endolysosomal escape of tau seeds and propagation of tau aggregation. J Biol Chem 294, 18952–18966 (2019). 10.1074/jbc.RA119.009432

56 Klumperman, J. & Raposo, G. The complex ultrastructure of the endolysosomal system. Cold Spring Harb Perspect Biol 6, a016857 (2014). 10.1101/cshperspect.a016857

57 Ralston, K. S. Taking a bite: Amoebic trogocytosis in Entamoeba histolytica and beyond. Curr Opin Microbiol 28, 26–35 (2015). 10.1016/j.mib.2015.07.009

58 Martinez-Martin, N. et al. T cell receptor internalization from the immunological synapse is mediated by TC21 and RhoG GTPase-dependent phagocytosis. Immunity 35, 208–222 (2011). 10.1016/j.immuni.2011.06.003

59 Lou, J., Low-Nam, S. T., Kerkvliet, J. G. & Hoppe, A. D. Delivery of CSF-1R to the lumen of macropinosomes promotes its destruction in macrophages. J Cell Sci 127, 5228–5239 (2014). 10.1242/jcs.154393

60 Campellone, K. G. & Welch, M. D. A nucleator arms race: cellular control of actin assembly. Nat Rev Mol Cell Biol 11, 237–251 (2010). 10.1038/nrm2867

61 Colucci-Guyon, E. et al. A role for mammalian diaphanous-related formins in complement receptor (CR3)-mediated phagocytosis in macrophages. Curr Biol 15, 2007–2012 (2005). 10.1016/j.cub.2005.09.051

62 Roberts, A. W. et al. Cas9(+) conditionally-immortalized macrophages as a tool for bacterial pathogenesis and beyond. Elife 8 (2019). 10.7554/eLife.45957

63 Alharbi, M. G., Lee, S. H., Abdelazim, A. M., Saadeldin, I. M. & Abomughaid, M. M. Role of Extracellular Vesicles in Compromising Cellular Resilience to Environmental Stressors. Biomed Res Int 2021, 9912281 (2021). 10.1155/2021/9912281

64 Crewe, C. Energetic Stress-Induced Metabolic Regulation by Extracellular Vesicles. Compr Physiol 13, 5051–5068 (2023). 10.1002/cphy.c230001

65 Qadeer, A. et al. Extracellular Vesicles Associated Metabolites as Intercellular Signalling Mediators in Disease and Therapy. Metabolites 16 (2026). 10.3390/metabo16030207

66 Dong, X. et al. Tunneling nanotubes: a new dimension of intercellular communication and recent progress. Cell Commun Signal 24 (2026). 10.1186/s12964-026-02980-3

67 Wang, B. et al. Context-Dependent Functional Outcomes of Mitochondrial Transfer: A Donor-Recipient Perspective. Adv Biol (Weinh) 10, e70137 (2026). 10.1002/adbi.70137

68 Epelbaum, S. et al. Acute amnestic encephalopathy in amyloid-beta oligomer-injected mice is due to their widespread diffusion in vivo. Neurobiol Aging 36, 2043–2052 (2015). 10.1016/j.neurobiolaging.2015.03.005

69 Manassero, G. et al. Dual Mechanism of Toxicity for Extracellular Injection of Tau Oligomers versus Monomers in Human Tau Mice. J Alzheimers Dis 59, 743–751 (2017). 10.3233/JAD-170298

70 Sacino, A. N. et al. Brain injection of alpha-synuclein induces multiple proteinopathies, gliosis, and a neuronal injury marker. J Neurosci 34, 12368–12378 (2014). 10.1523/JNEUROSCI.2102-14.2014

71 Zhou, X. et al. Circuit Design Features of a Stable Two-Cell System. Cell 172, 744–757 e717 (2018). 10.1016/j.cell.2018.01.015

72 Wang, J. et al. UV-induced somatic mutations elicit a functional T cell response in the YUMMER1.7 mouse melanoma model. Pigment Cell Melanoma Res 30, 428–435 (2017). 10.1111/pcmr.12591

73 Bray, N. L., Pimentel, H., Melsted, P. & Pachter, L. Near-optimal probabilistic RNA-seq quantification. Nat Biotechnol 34, 525–527 (2016). 10.1038/nbt.3519

74 Florsheim, E. B. et al. Immune sensing of food allergens promotes avoidance behaviour. Nature 620, 643–650 (2023). 10.1038/s41586-023-06362-4

75 Ip, W. K. E., Hoshi, N., Shouval, D. S., Snapper, S. & Medzhitov, R. Anti-inflammatory effect of IL-10 mediated by metabolic reprogramming of macrophages. Science 356, 513–519 (2017). 10.1126/science.aal3535

76 Freund, E. C. et al. Efficient gene knockout in primary human and murine myeloid cells by non-viral delivery of CRISPR-Cas9. J Exp Med 217 (2020). 10.1084/jem.20191692

77 Quinlan, A. R. & Hall, I. M. BEDTools: a flexible suite of utilities for comparing genomic features. Bioinformatics 26, 841–842 (2010). 10.1093/bioinformatics/btq033

78 Obara, C. J. et al. Motion of VAPB molecules reveals ER-mitochondria contact site subdomains. Nature 626, 169–176 (2024). 10.1038/s41586-023-06956-y

