## Supplementary Fig. 1 for "Non-cell-autonomous removal of protein aggregates via trogocytosis"

**Supplementary Information**

**Contents:**

I. **Supplementary Methods**
II. **Supplementary Figures**
III. **Supplementary Tables**
IV. **Supplementary Videos**

**Supplementary Methods**

**Plasmid cloning**

tmCSF1 and sCSF1 cDNAs were amplified by PCR, using cDNA templates synthesized from total RNA extract of YUMMER1.7 cells. Primers used are as follows: 5′-TTCTCTAGGCGCCGGAATTAATGACCGCGCGGGGCGCC-3′ and 5′-TAGGGGGGGGGGCGGAATTCGTTCTATACTGGCAGTTCCACCTGTCTGTCCTCATCCTGG-3′. Amplified tmCSF1 and sCSF1 cDNAs were separated with agarose gel electrophoresis and cloned into BglII-HpaI site of pMSCVpuro-IRES-GFP (pMIG)^1^. For cloning FLAG-tagged tmCSF1 (FLAG-tmCSF1), the signal sequences and the downstream sequences of tmCSF1 were separately amplified by PCR from pMIG-cloned tmCSF1, using the primers: 5′-TATAGGGAGACCCAAGCTGGATGACCGCGCGGGGCGCC-3′ and 5′-CCTTATAATCCTTGTCGTCATCGTCTTTGTAGTCGATATCATGATCCTTATAATCACCATCATGATCCTTATAATCCACCTC-3′ for the signal sequences and 5′-TGACGACAAGAAGGAGGTGTCAGAACAC-3′ and 5′-CGGCCGCCACTGTGCTGGATCTATACTGGCAGTTCCAC-3′ for the downstream sequences. PCR products were ligated with the oligonucleotides encoding 3 × FLAG (produced by annealing 5′-TGACGACAAGGATTATAAGGATCATGATGGTGATTATAAGGATCATGATATCGACTACAAAGACGATGACGACAAG-3′ and 5′-ACACCTCCTTCTTGTCGTCATCGTCTTTGTAGTCGATATCATGATCCTTATAATCACCATCATGATCCTTATAATC-3′) and pcDNA3.1^2^ that was cut by EcoRV and NheI. Subsequently, FLAG-tmCSF1 sequences were amplified by PCR with the primers: 5′-TTCTCTAGGCGCCGGAATTAATGACCGCGCGGGGCGCC-3′ and 5′-TAGGGGGGGGGGCGGAATTCGTTCTATACTGGCAGTTCCACCTGTCTGTCCTCATCCTGG-3′. PCR products were cloned into BglII-HpaI site of pMIG. To swap the cytoplasmic tail of FLAG-tmCSF1 with that of CD4, the region from the signal sequences to the transmembrane domain sequences of FLAG-tmCSF1 was amplified by PCR using pMIG-cloned FLAG-tmCSF1. Primers used are as follows: 5′-TTCTCTAGGCGCCGGAATTAATGACCGCGCGGGGCGCC-3′ and 5′-TGACACAGCAGTAGAACAGGAGGCCCCCAACAGTCAGC-3′. Sequences encoding the cytoplasmic tail of CD4 were amplified using cDNAs synthesized from total RNA extract of mouse spleen. Primers used are as follows: 5′-CCTGTTCTACTGCTGTGTCAGGTGCCGG-3′ and 5′-GGGGGGGGGCGGAATTCGTTCTAGATGAGATTATGGCTCTTCTGCATCC-3′. DNA fragments were cloned together into BglII-HpaI site of pMIG. To delete the PDZ-binding domain in FLAG-tmCSF1, sequences encoding FLAG-tmCSF1 without 3 amino acids at the C-terminus were amplified by PCR with the primers: 5′-TTCTCTAGGCGCCGGAATTAATGACCGCGCGGGGCGCC-3′and 5′-GGGGGGGGGCGGAATTCGTTCTATTCCACCTGTCTGTCCTCATCCTGGGTCAGG-3′. PCR products were cloned into BglII-HpaI site of pMIG.

pLVX-TetOne-puro (TaKaRa) was cut by AvrII and MruI and ligated with blasticidin-resistant gene-encoding DNA fragments amplified by PCR from pMSCV-Blasticidin (Addgene, 75085). Primers used are as follows: 5′-GAGGAGGCTTTTTTGGAGGCCTAGGCTTTTGCAAAACGCGACCATGGCCAAGCCTTTGTCTC-3′ and 5′-CAGAGGTTGATTGTTCCAGACGCGTTTAGCCCTCCCACACATAAC-3′. The resultant vector (pTetOne-blast) was used for the cloning of HTT-Q103. Sequences encoding mCherry-tagged HTT-Q103 were amplified by PCR with the primers: 5′-CACTTCCTACCCTCGTAAAGAAACCGCCATGGCGACCCTG-3′ and 5′-CTCGCAGGGGAGGTGGTCTGGATCCTCATTACTTATACAGCTCGTCCATG -3′. PCR products were cloned into EcoRI-BamHI site of pTetOne-blast.

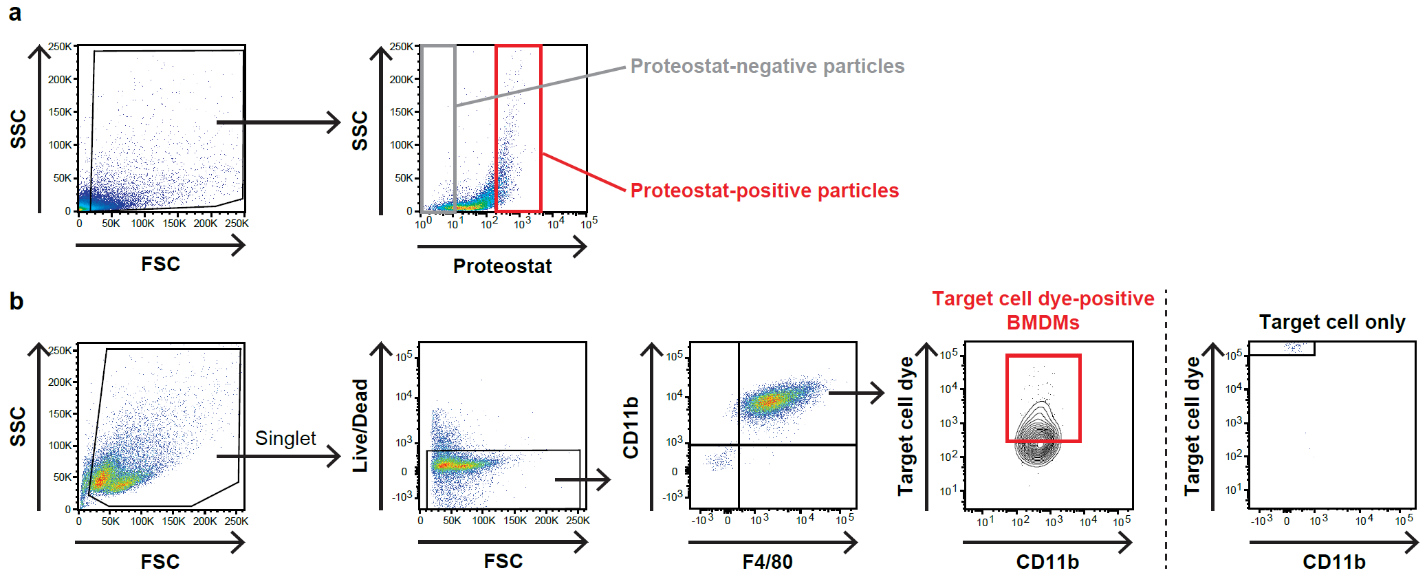

**Supplementary Fig. 1. Gating strategy in flow cytometry analysis**

**a,** Gating strategy for sorting proteostat-positive and negative particles from MEF^Baf A1^ lysate. **b,** Gating strategy for analyzing the uptake of materials from target cells by BMDMs.

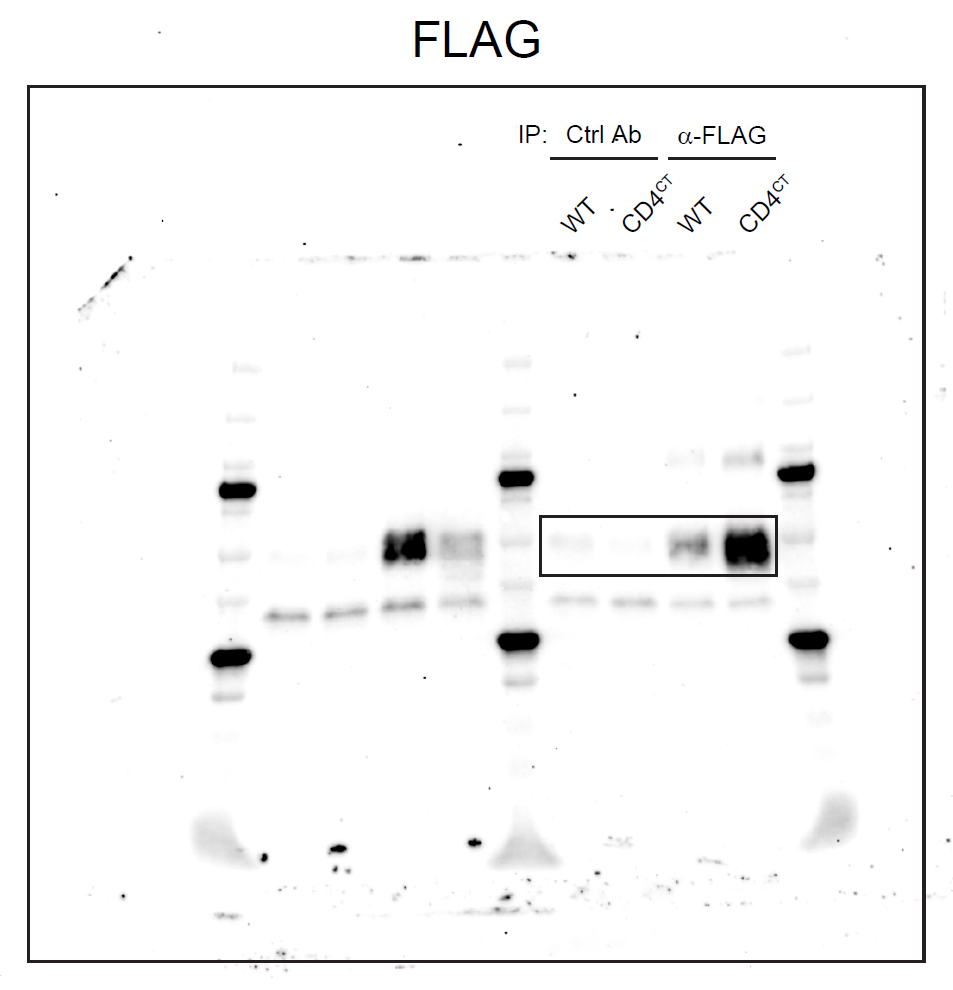

**Supplementary Fig. 2. Uncropped images of immunoblotting for Fig. 4j.**

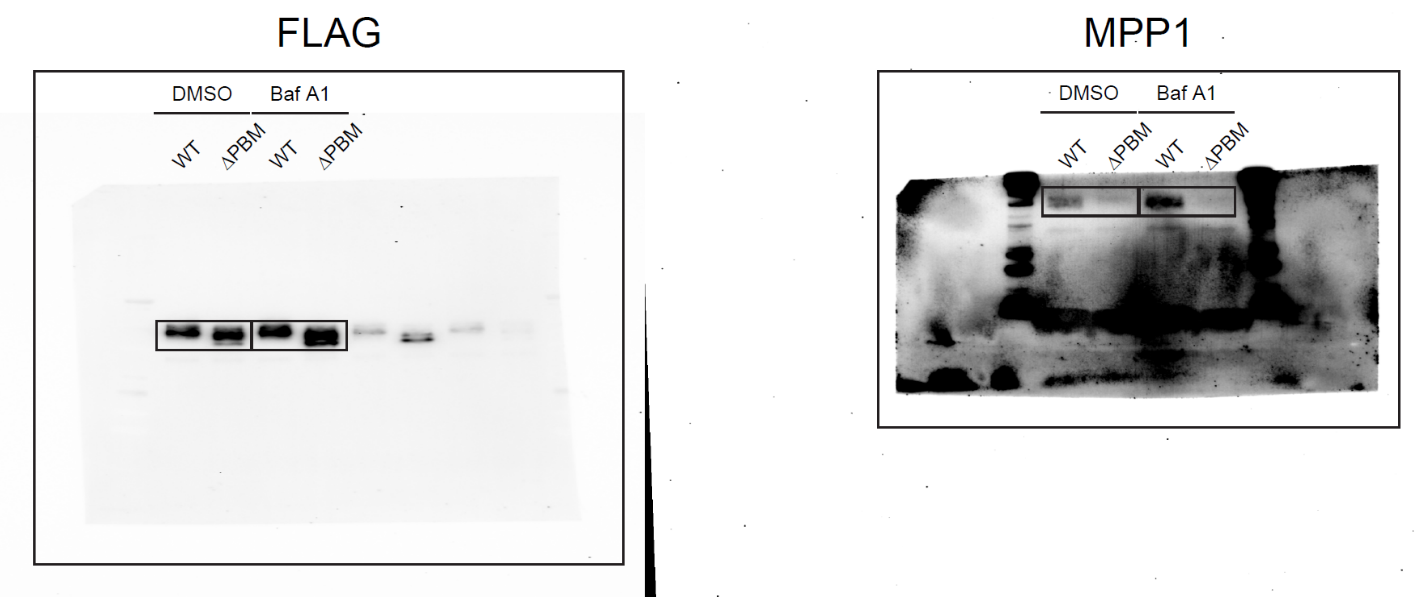

**Supplementary Fig. 3. Uncropped images of immunoblotting for Fig. 6i.**

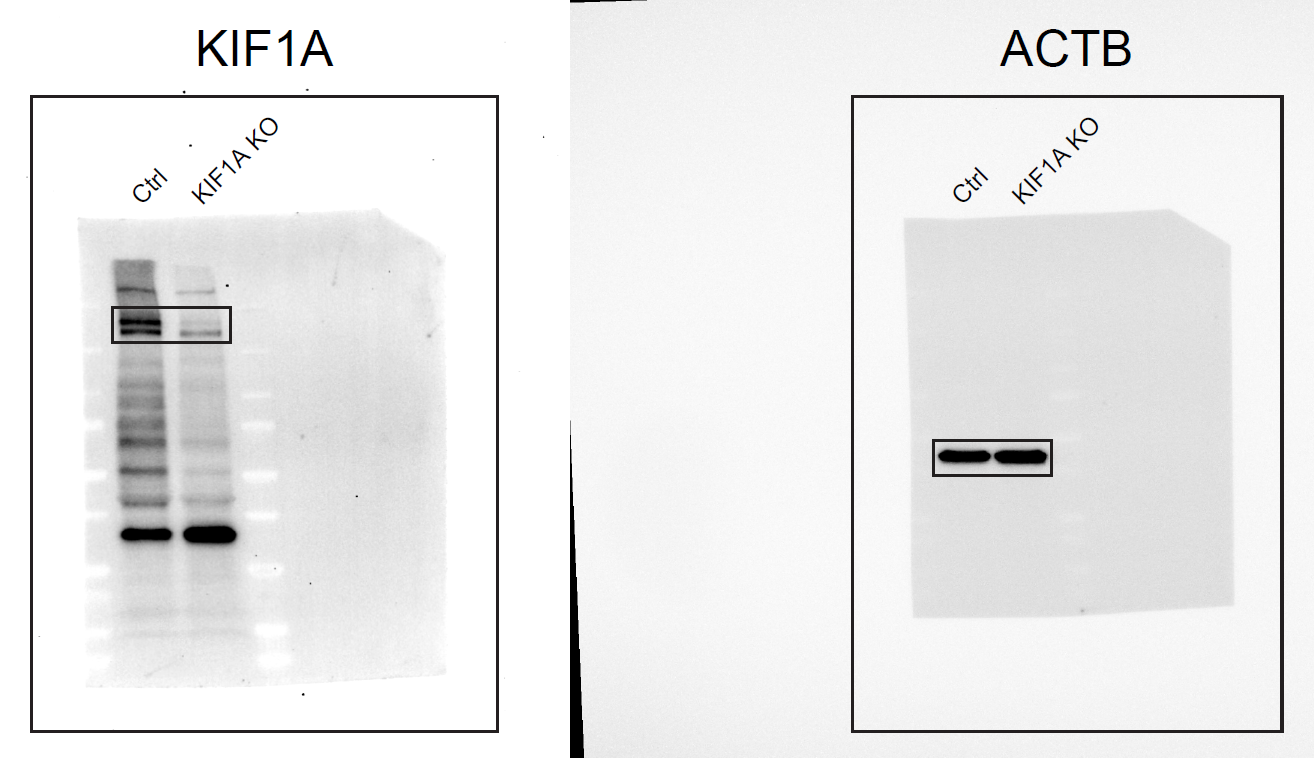

**Supplementary Fig. 4. Uncropped images of immunoblotting for Extended Data Fig. 6c (left panel).**

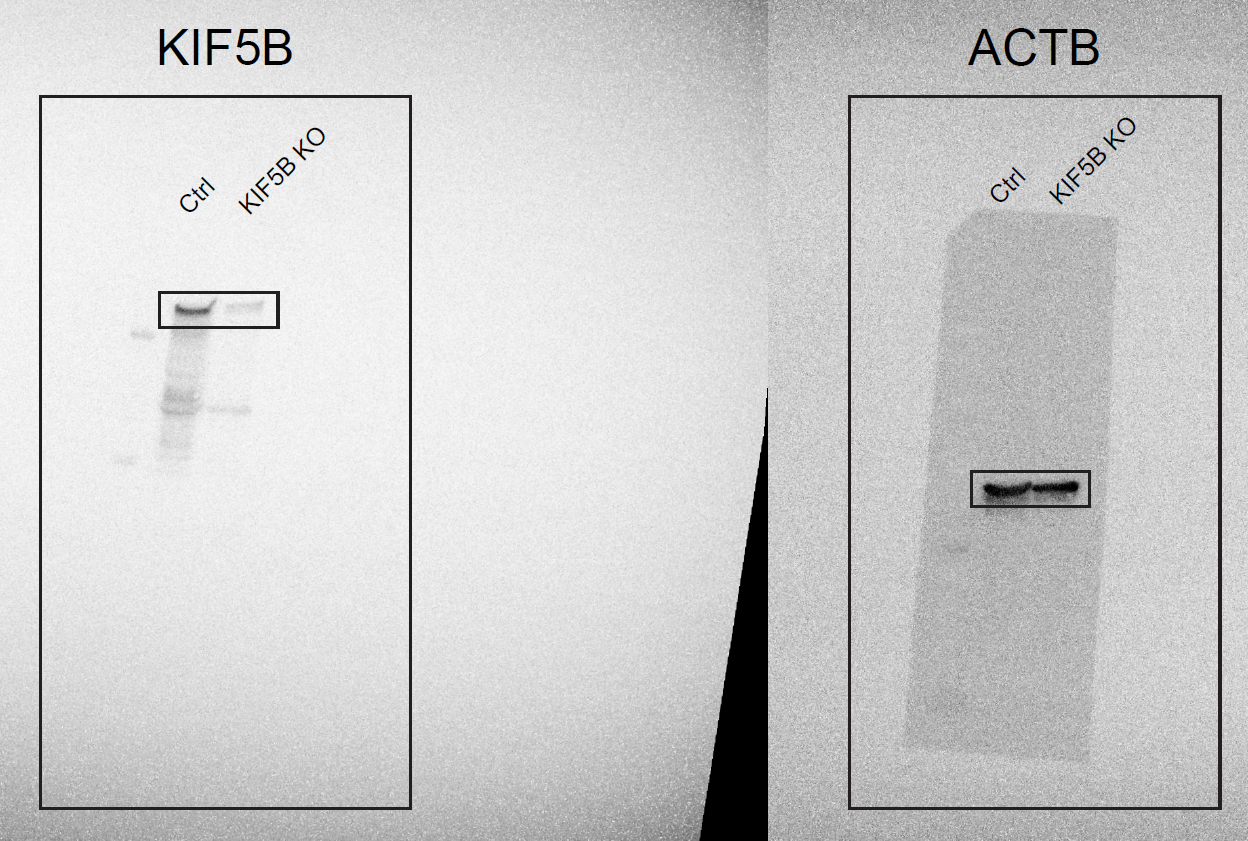

**Supplementary Fig. 5. Uncropped images of immunoblotting for Extended Data Fig. 6c (right panel).**

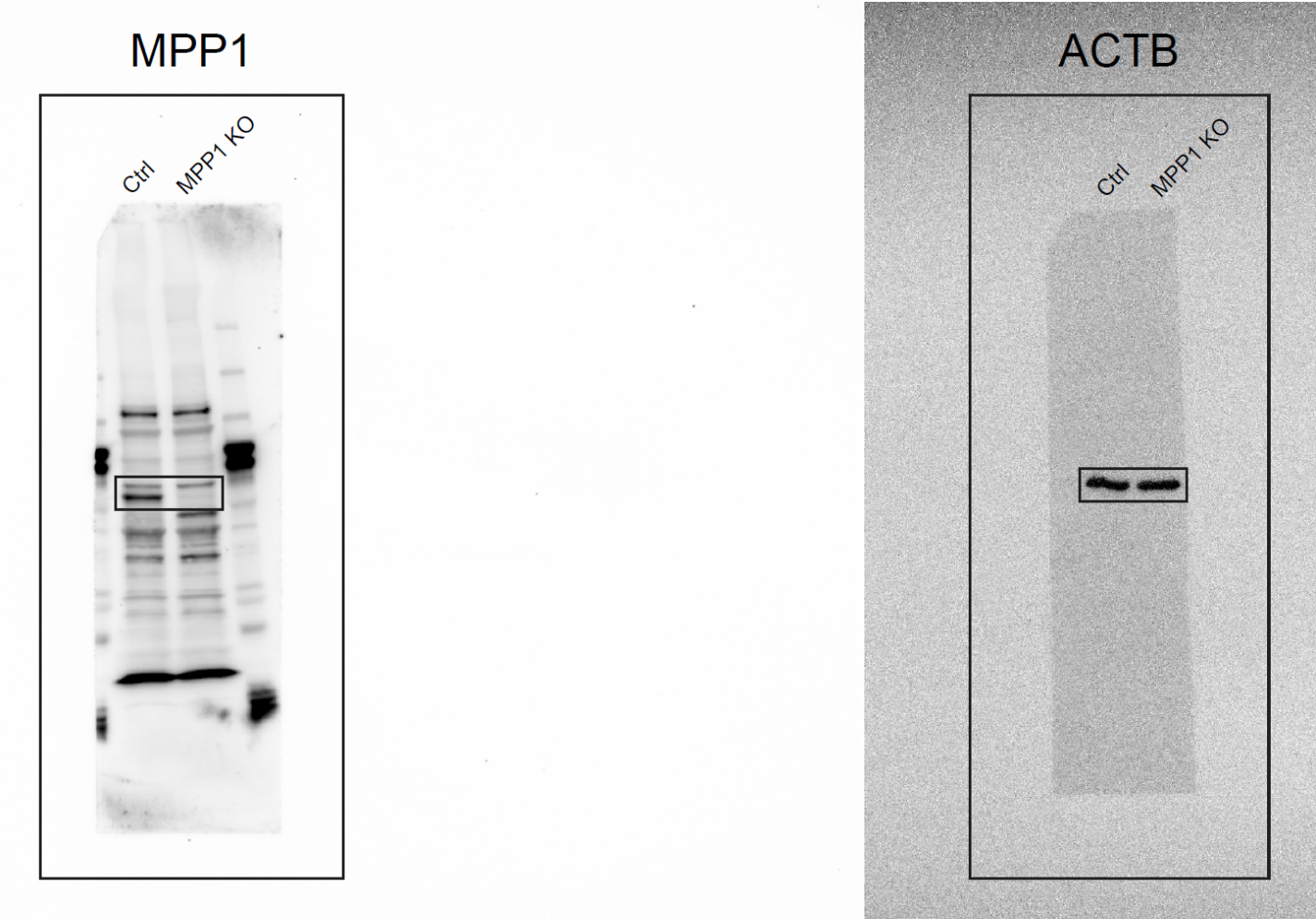

**Supplementary Fig. 6. Uncropped images of immunoblotting for Extended Data Fig. 6f.**

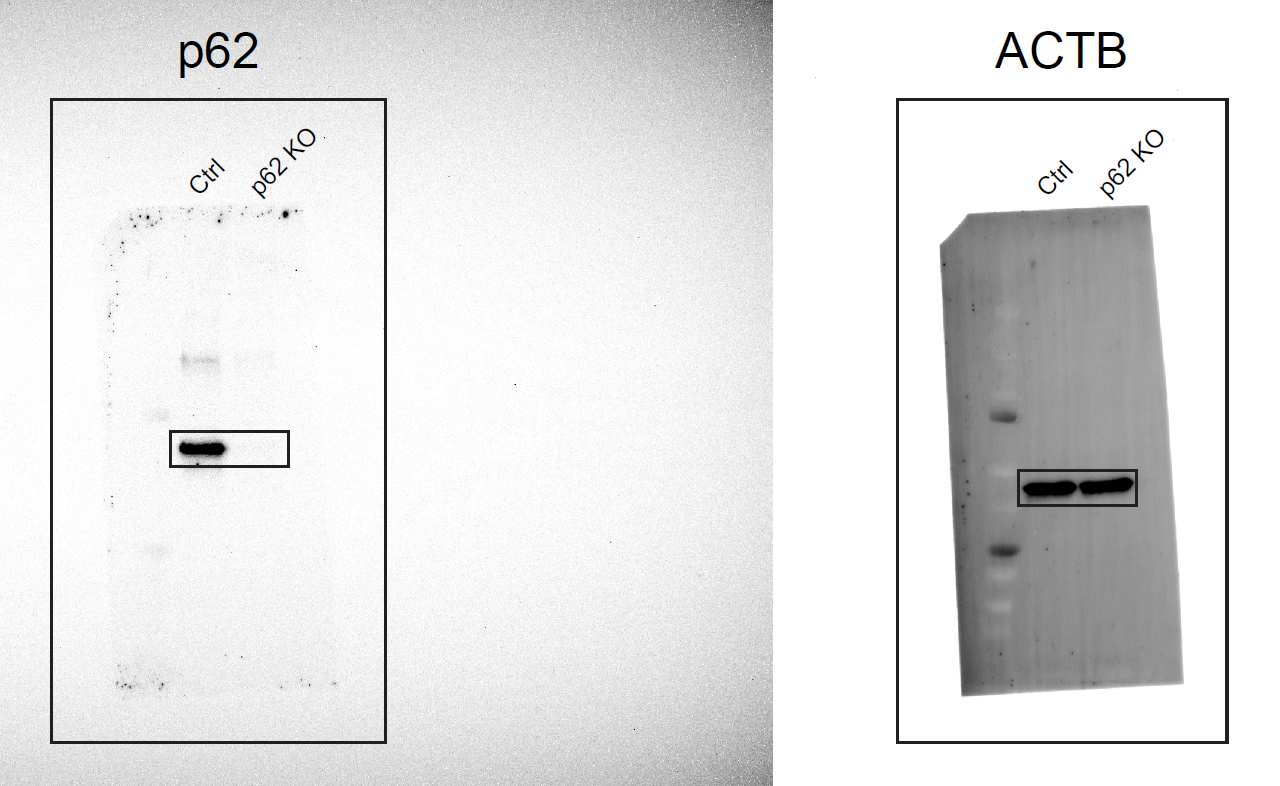

**Supplementary Fig. 7. Uncropped images of immunoblotting for Extended Data Fig. 7c (left panel).**

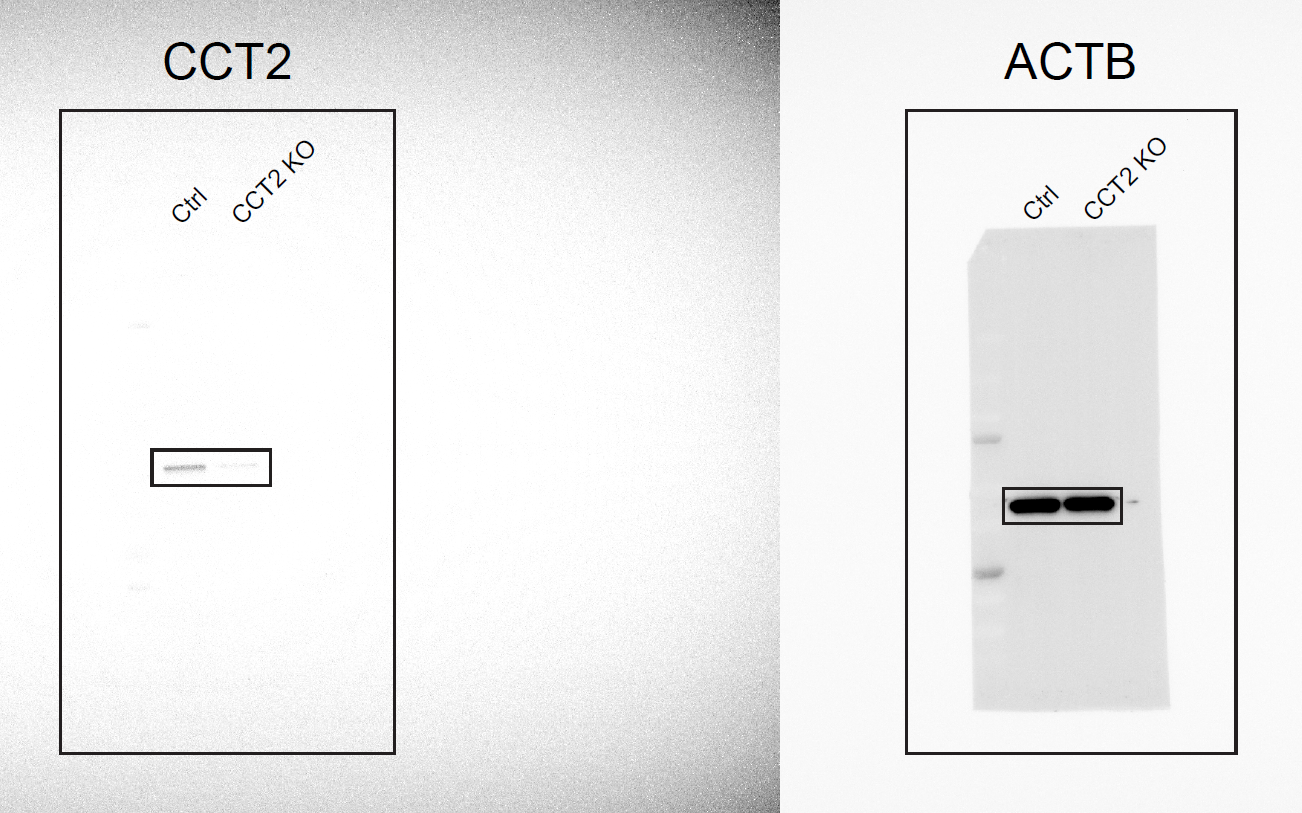

**Supplementary Fig. 8. Uncropped images of immunoblotting for Extended Data Fig. 7c (second panel from the left).**

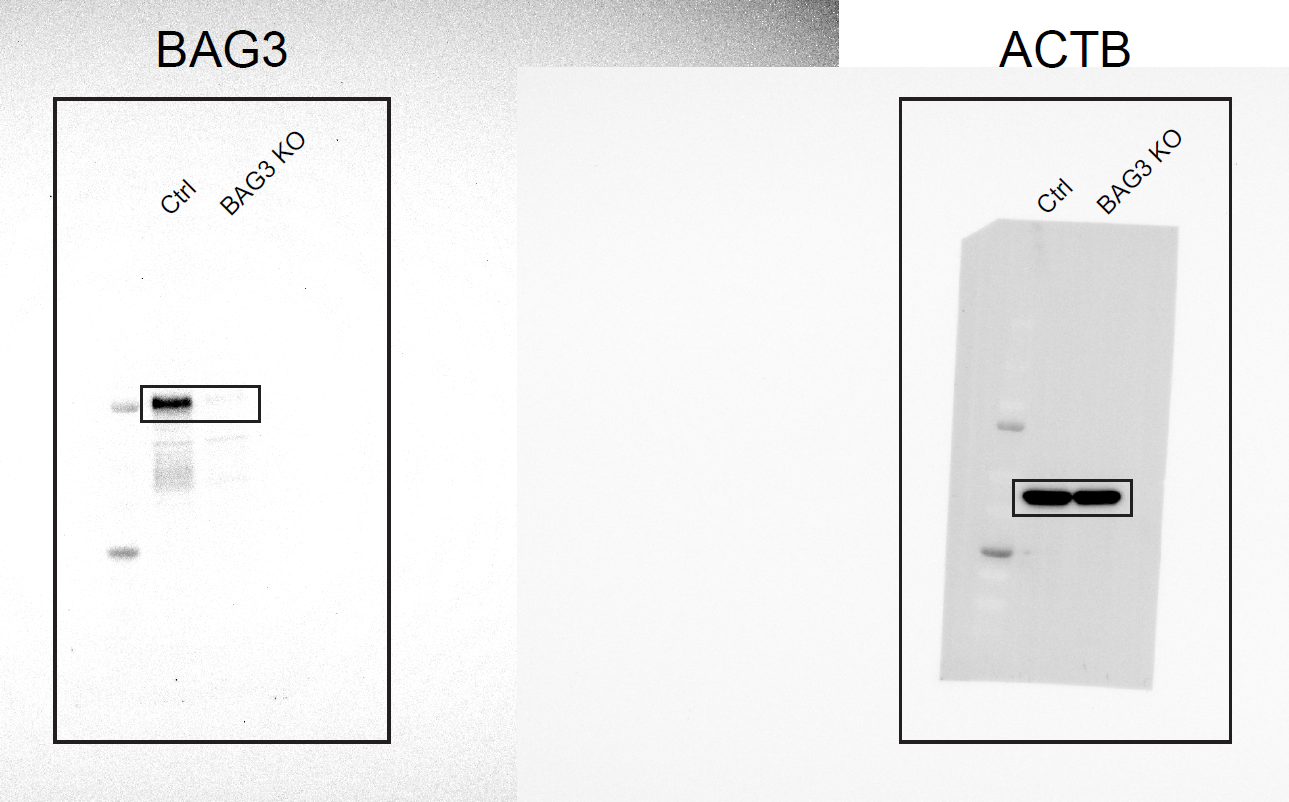

**Supplementary Fig. 9. Uncropped images of immunoblotting for Extended Data Fig. 7c (second panel from the right).**

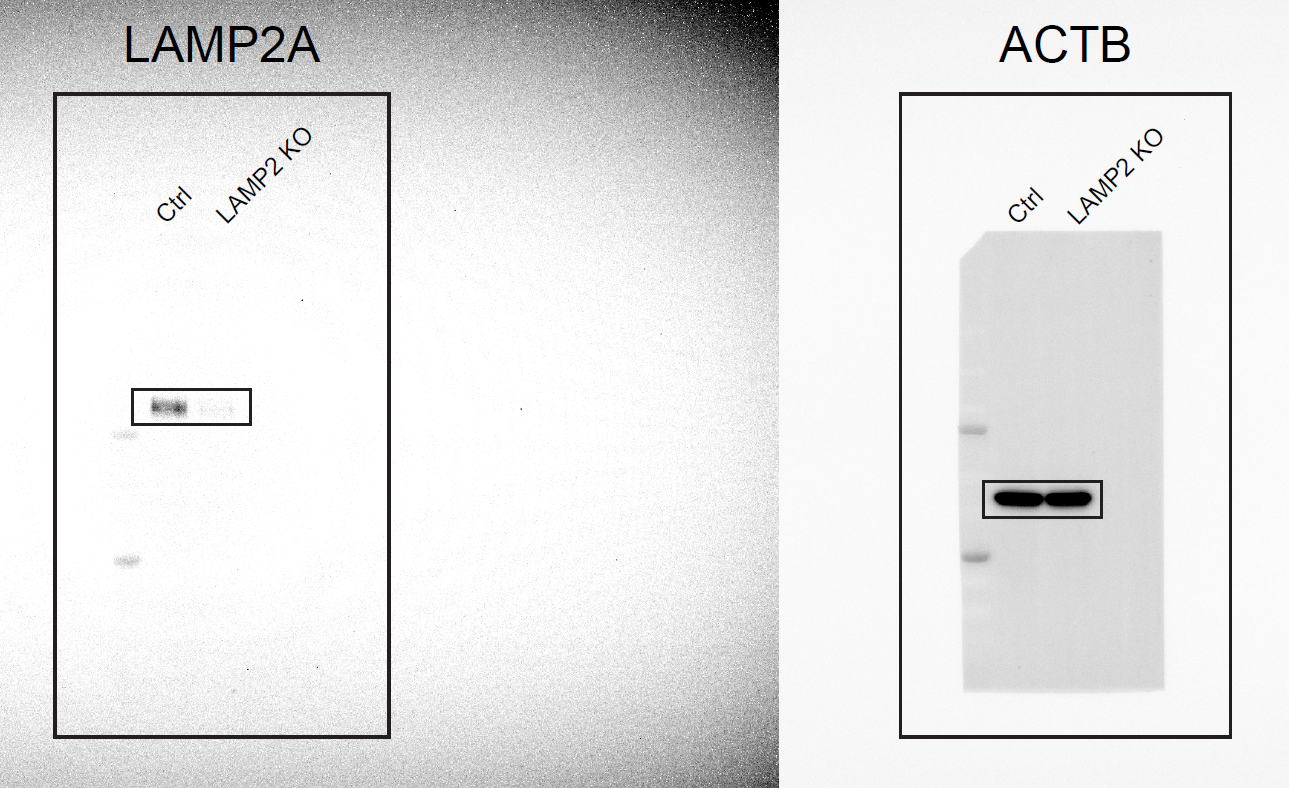

**Supplementary Fig. 10. Uncropped images of immunoblotting for Extended Data Fig. 7c (right panel).**

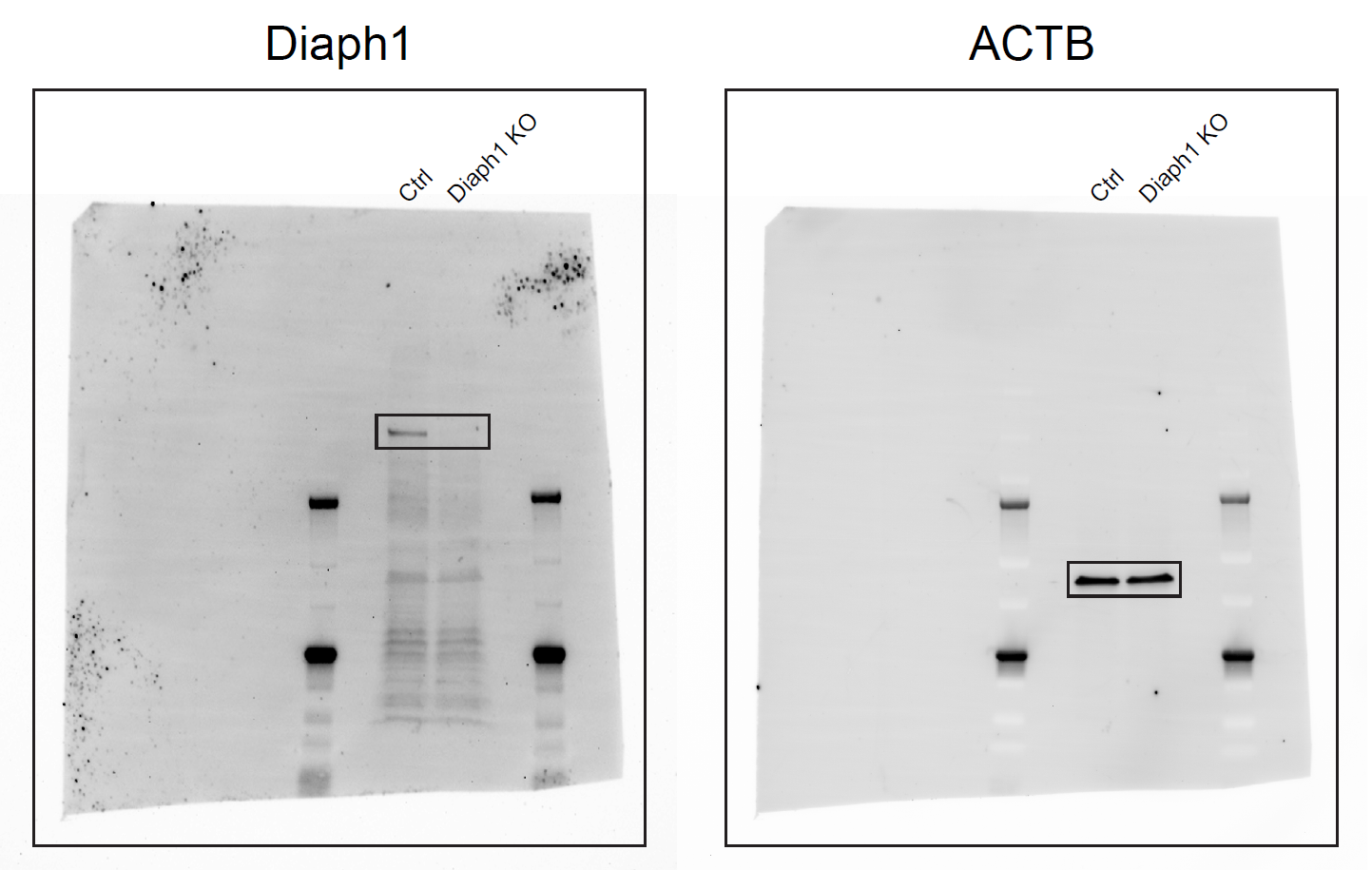

**Supplementary Fig. 11. Uncropped images of immunoblotting for Extended Data Fig. 10b.**

| *tmCsf1* | Forward | CCAAGAACTGCAACAACAGC |
| --- | --- | --- |
|  | Reverse | CCAACAGTCAGCAAGACTAGGA |
| *sCsf1* | Forward | GGGATTTTTGACCCAGGAAG |
|  | Reverse | GGAATGGAGATGCTGAGAGG |
| *Ctsb* | Forward | TGCGTTCGGTGAGGACATAG |
|  | Reverse | CCAAATGCCCAACAAGAGCC |
| *Tfeb* | Forward | CTCCAACCCCGAGAAAGAGT |
|  | Reverse | GTTCAGGTGGCTGCTAGACA |
| *Mitf* | Forward | TTTCCCTTATCCCATCCACCG |
|  | Reverse | TCGTACATGTCAGGTCTGCC |
| *Tfe3* | Forward | GTGTACAGCAGTCAGGGAGT |
|  | Reverse | TGTCTTTCTTCTGTCGCTCCT |
| *Tfec* | Forward | TACCCACATCACCAAAGCAGC |
|  | Reverse | GCTCATAGAACTCCGGGCTT |
| *Ppil4* | Forward | GGTCGAATAGGAGCAGATGAA |
|  | Reverse | CATCAGGTAAGTCTCCCACCA |
| *Mtpap* | Forward | TGGTGTTCAGTGTGCGATG |
|  | Reverse | GTTGGAAGAATGGGAGGTGA |
| *Rpl13a* | Forward | GAAGGAAAAGGCCAAGATGCAC |
|  | Reverse | TGAGGACCTCTGTGAACTTGC |

**Supplementary Table 1. Primers used in qPCR analysis**

| Primary antibodies | | | | |
| --- | --- | --- | --- | --- |
| Antibody target | Conjugation | Clone | Catalog # | Source |
| CD11b | BUV395 | M1/70 | 363-0112-82 | Thermo Fisher Scientific |
| CD11b | APC | M1/70 | 17-0112-82 | Thermo Fisher Scientific |
| CD11b | BV421 | M1/70 | 562605 | BD Biosciences |
| F4/80 | PE/Cy7 | BM8 | 123113 | BioLegend |
| CD115 | PE | AFS98 | 14-1152-82 | Thermo Fisher Scientific |
| Isotype control | Biotin | N/A  (polyclonal goat IgG) | BAF108 | R & D Systems |
| CSF1 | Biotin | N/A  (polyclonal goat IgG) | BAF416 | R & D Systems |
| Streptavidin | | | | |
| Streptavidin | Alexa Fluor 647 | N/A | S32357 | Thermo Fisher Scientific |

**Supplementary Table 2. Antibodies used in flow cytometry**

| Plasmid name | Catalog # | Source |
| --- | --- | --- |
| mEmerald-CD81-10 | 54031 | Addgene |
| pBetaActin-TUBB2a-mScarlet | 191324 |  |
| pJC49 | 133060 |  |

**Supplementary Table 3. Commercially available plasmids used for transfection**

| siRNA target | siRNA ID | Source |
| --- | --- | --- |
| *Tfeb* | SASI_Mm01_00082196 | Sigma Aldrich |
| *Mitf* | SASI_Mm02_00307204 |  |
| *Tfe3* | SASI_Mm01_00085148 |  |
| *Tfec* | SASI_Mm01_00198518 |  |
| *Ppil4* | SASI_Mm02_00329749 |  |
| *Mtpap* | SASI_Mm01_00079345 |  |
| Universal Negative Control #1 | N/A |  |

**Supplementary Table 4. siRNAs used for gene knockdown**

| Guide RNA target |  | Sequence (5' to 3') |
| --- | --- | --- |
| *Kif1a* | 1 | CAAAGGCAACCUGCGUGUGA |
|  | 2 | GAGGUCCUGGAUGUCAUUGU |
| *Kif5b* | 1 | AUCUGGGAAGACCCACACGA |
|  | 2 | AAGAUAAGGGACUUGUUAGA |
| *Mpp1* | 1 | UCAGCUUCAAAGUGAUCCCC |
|  | 2 | GAUCUCAUCCCCAACAUGAA |
| *p62* | 1 | UCGCCCACCAUGUGCUCAGG |
|  | 2 | CGGCCAUCCCCUGCACGAGG |
| *Cct2* | 1 | GUCGUUGGUCACCAUCAGAG |
|  | 2 | GUUGAACGUAUUCUUAAGCA |
| *Bag3* | 1 | ACAACAGCCGCACCACGACG |
|  | 2 | CCUGUAUACCACAAGAUCCA |
| *Lamp2* | 1 | GAUCACGAUGUGCCUCUCUC |
|  | 2 | ACACGAUGGAAGCAGUUGUG |

**Supplementary Table 5. Single-guide RNAs used for gene knockout**

| Primary antibodies | | | |
| --- | --- | --- | --- |
| Antibody target | Clone | Catalog # | Source |
| LC3 | N/A (polyclonal rabbit IgG) | 14600-1-AP | Proteintech |
| LAMP1 | 1D4B | 65050-1-Ig | Proteintech |
| CD81 | D5O2Q | 10037S | Cell Signaling Technologies |
| CSF1 | N/A (polyclonal goat IgG) | BAF416 | R & D Systems |
| MPP1 | N/A (polyclonal rabbit IgG) | 14715-1-AP | Proteintech |
| ALIX | N/A (polyclonal rabbit IgG) | 12422-1-AP | Proteintech |
| TFG | N/A (polyclonal rabbit IgG) | 11571-1-AP | Proteintech |
| SPG20 | N/A (polyclonal rabbit IgG) | 13791-1-AP | Proteintech |
| ITCH | N/A (polyclonal rabbit IgG) | 20920-1-AP | Proteintech |
| Ubiquitin | EPR8830 | ab134953 | Abcam |
| Ubiquitin | GT7811 | MA5-42345 | Thermo Fisher Scientific |
| p62 | EPR4844 | ab109012 | Abcam |
| CCT2 | EPR4084 | ab92746 | Abcam |
| TAX1BP1 | EPR13287(B) | ab176572 | Abcam |
| BAG3 | E7Y9T | 23842T | Cell Signaling Technologies |
| Hsc70 | EP1531Y | ab51052 | Abcam |
| HDAC6 | N/A (polyclonal rabbit IgG) | PA5-86320 | Thermo Fisher Scientific |
| γ-tubulin | EPR16793 | ab179503 | Abcam |
| Vimentin | D21H3 | 5741S | Cell Signaling Technologies |
| CD11b | M1/70 | 612800 | BD Biosciences |
| Secondary antibodies and streptavidin | | | |
| Antibody target | Conjugation | Catalog # | Source |
| Mouse IgG | Alexa Fluor Plus 405 | A48257 | Thermo Fisher Scientific |
| Rabbit IgG | Alexa Fluor Plus 488 | A32731 | Thermo Fisher Scientific |
| Mouse IgG | Alexa Fluor Plus 488 | A32723 | Thermo Fisher Scientific |
| Rat IgG | Alexa Fluor 647 | A21472 | Thermo Fisher Scientific |
| Streptavidin | Alexa Fluor 488 | S32354 | Thermo Fisher Scientific |
| Streptavidin | Alexa Fluor 647 | S32357 | Thermo Fisher Scientific |

**Supplementary Table 6. Antibodies used in immunofluorescence staining**

| Antibody target | Clone | Catalog # | Source |
| --- | --- | --- | --- |
| DYKDDDDK (FLAG) | D6W5B | 14793S | Cell Signaling Technologies |
| KIF1A | EPR24329-6 | ab271047 | Abcam |
| KIF5B | N/A (polyclonal rabbit IgG) | 21632-1-AP | Proteintech |
| MPP1 | N/A (polyclonal rabbit IgG) | 14715-1-AP | Proteintech |
| ACTB | 8H10D10 | 3700 | Cell Signaling Technologies |
| p62 | EPR4844 | ab109012 | Abcam |
| CCT2 | EPR4084 | ab92746 | Abcam |
| LAMP2A | F1A5A | 81197T | Cell Signaling Technologies |
| Diaph1 | D6W5B | 14793S | Cell Signaling Technologies |
| Secondary antibodies | | | |
| Antibody target | Fluorophore | Catalog # | Source |
| Rabbit IgG | Alexa Fluor Plus 800 | A32735 | Thermo Fisher Scientific |
| Mouse IgG | Alexa Fluor Plus 488 | A32723 | Thermo Fisher Scientific |

**Supplementary Table 7. Antibodies used in immunoblotting**

**Supplementary Video 1. Materials in protein aggregate-accumulating cells are captured and engulfed by macrophages.**

Representative time-lapse confocal images of co-cultured BMDMs and CTDR-labelled MEF^Baf A1^. White arrow indicates MEF^Baf A1^-derived materials incorporated by BMDMs. Images were taken every 2 min.

**Supplementary Video 2. Materials in protein aggregate-accumulating cells are pulled away and removed by macrophages.**

Representative time-lapse confocal images of co-cultured BMDMs and CTDR-labelled MEF^Baf A1^. White arrow indicates MEF^Baf A1^-derived materials incorporated by BMDMs. Images were taken every 2 min.

**Supplementary Video 3. Cell bodies containing electron-dense materials are trogocytosed by macrophages from protein aggregate-accumulating cells.**

FIB-SEM images of co-cultured MEF^Baf A1^ and BMDMs. The top and bottom cells show MEF^Baf A1^ and BMDM, respectively. Electron-dense area (red) and plasma membranes of MEF^Baf A1^ (light blue) and BMDM (dark blue) are segmented and reconstructed as a 3D image.

**Supplementary Video 4. Materials of tmCSF1-expressing cells are incorporated into macrophages.**

Representative time-lapse confocal images of co-cultured mock-transduced cells, tmCSF1-transduced cells and BMDMs. Images taken every 2 min are shown.

**Supplementary Video 5. Protein aggregates are anterogradely transported along microtubule.**

Representative time-lapse confocal images of MEF^Baf A1^. Arrows indicate CTDR puncta that move along microtubule. Images taken every 25 sec are shown.

**References**

1	Ip, W. K. E., Hoshi, N., Shouval, D. S., Snapper, S. & Medzhitov, R. Anti-inflammatory effect of IL-10 mediated by metabolic reprogramming of macrophages. *Science* **356**, 513-519 (2017). <https://doi.org:10.1126/science.aal3535>

2	Ye, J. J., Bian, X., Lim, J. & Medzhitov, R. Adiponectin and related C1q/TNF-related proteins bind selectively to anionic phospholipids and sphingolipids. *Proc Natl Acad Sci U S A* **117**, 17381-17388 (2020). <https://doi.org:10.1073/pnas.1922270117>
